# An integrated personal and population-based Egyptian genome reference

**DOI:** 10.1101/681254

**Authors:** Inken Wohlers, Axel Künstner, Matthias Munz, Michael Olbrich, Anke Fähnrich, Verónica Calonga-Solís, Caixia Ma, Misa Hirose, Shaaban El-Mosallamy, Mohamed Salama, Hauke Busch, Saleh Ibrahim

## Abstract

The human genome is composed of chromosomal DNA sequences consisting of bases A, C, G and T – the blueprint to implement the molecular functions that are the basis of every individual’s life. Deciphering the first human genome was a consortium effort that took more than a decade and considerable cost. With the latest technological advances, determining an individual’s entire personal genome with manageable cost and effort has come within reach. Although the benefits of the all-encompassing genetic information that entire genomes provide are manifold, only a small number of *de novo* assembled human genomes have been reported to date ^1–3^, and few have been complemented with population-based genetic variation ^4^, which is particularly important for North Africans who are not represented in current genome-wide data sets ^5–7^. Here, we combine long- and short-read whole-genome next-generation sequencing data with recent assembly approaches into the first *de novo* assembly of the genome of an Egyptian individual. The resulting assembly demonstrates well-balanced quality metrics and is complemented with high-quality variant phasing via linked reads into haploblocks, which we can associate with gene expression changes in blood. To construct an Egyptian genome reference, we further assayed genome-wide genetic variation occurring in the Egyptian population within a representative cohort of 110 Egyptian individuals. We show that differences in allele frequencies and linkage disequilibrium between Egyptians and Europeans may compromise the transferability of European ancestry-based genetic disease risk and polygenic scores, substantiating the need for multi-ethnic genetic studies and corresponding genome references. The Egyptian genome reference represents a comprehensive population data set based on a high-quality personal genome. It is a proof of concept to be considered by the many national and international genome initiatives underway. More importantly, we anticipate that the Egyptian genome reference will be a valuable resource for precision medicine targeting the Egyptian population and beyond.

## Main

With the advent of personal genomics, population-based genetics as part of an individual’s genome is indispensable for precision medicine. Currently, genomics-based precision medicine compares the patients’ genetic make-up to a reference genome ^8^, a genome model inferred from individuals of mostly European descent, to detect risk mutations that are related to disease. However, genetic and epidemiologic studies have long recognized the importance of ancestral origin in conferring genetic risk for disease. Risk alleles and structural variants (SVs) ^9^ can be missing from the reference genome or can have different population frequencies, such that alternative pathways become disease related in patients of different ancestral origin, which motivates the establishment of national or international multi-ethnic genome projects ^6,7,10^. At present, there are several population-based sequencing efforts that aim to map specific variants in the 100,000 genome projects in Asia ^11^ or England ^12^. Furthermore, large-scale sequencing efforts currently explore population, society and history-specific genomic variations in individuals in Northern and Central Europe ^13,14^, North America ^7^, Asia ^15,16^ and, recently, the first sub-Saharan Africans ^17,18^. Nonetheless, there is still little genetic data available for many regions of the world. In particular, North African individuals are not adequately represented in current genetic data sets, such as the 1000 Genomes ^5^, TOPMED ^7^ or gnomAD ^6^ databases. Consequently, imminent health disparities between different world populations have been noted repeatedly for a decade. ^19–22^

In recent years, several high-quality *de novo* human genome assemblies ^1–4^ and, more recently, pan-genomes ^23^ have extended human sequence information and improved the *de facto* reference genome GRCh38 ^8^. Nonetheless, it is still prohibitively expensive to obtain all-embracing genetic information, such as high-quality *de novo* assembled personal genomes for many individuals. Indeed, previous genetic studies assess only a subset of variants occurring in the Egyptian population, e.g., single nucleotide polymorphisms (SNPs) on genotyping arrays ^24,25^, variants in exonic regions via exome sequencing ^26^ or variants detectable by short-read sequencing ^27,28^.

In this study, we generated a *de novo* assembly of an Egyptian individual and identified single nucleotide variants (SNVs) and SVs from an additional 109 Egyptian individuals obtained from short-read sequencing. Those were integrated to generate an Egyptian genome reference. We anticipate that an Egyptian population genome reference will strengthen precision medicine efforts that eventually benefit nearly 100 million Egyptians, e.g., by providing allele frequencies (AFs) and linkage disequilibrium (LD) between variants, information that is necessary for both rare and common disease studies. Likewise, our genome will be of universal value for research purposes, since it contains both European and African variant features. Most genome-wide association studies (GWAS) are performed in Europeans ^29^, but genetic disease risk may differ, especially for individuals of African ancestry ^30^. Consequently, an Egyptian genome reference will be well suited to support recent efforts to include Africans in such genetic studies, for example, by serving as a benchmark data set for SNP array construction and variant imputation or for fine-mapping of disease loci.

Our Egyptian genome is based on a high-quality human *de novo* assembly for one male Egyptian individual (see workflow in Suppl. Fig. 1). This assembly was generated from PacBio, 10x Genomics and Illumina paired-end sequencing data at overall 270x genome coverage (Suppl. Table 1). For this personal genome, we constructed two draft assemblies, one based on long-read assembly by an established assembler, `FALCON` ^31^, and another based on the assembly by a novel assembler, `WTDBG2` ^32^, which has a much shorter run time with comparable accuracy (cf. Suppl. Fig. 1). Both assemblies were polished using short reads and further polishing tools. For the `FALCON`-based assembly, scaffolding was performed, whereas we found that the `WTDBG2`-based assembly was of comparable accuracy without scaffolding (Table 1). Sex chromosomal sequences have not been manually curated. The `WTDBG2`-based assembly was selected as the meta assembly basis, because it performs similarly or better than the `FALCON`-based assembly, according to various quality control (QC) measures. The former did not require scaffolding, and thus there are no N bases or scaffolding errors. Overall, it has about 50% fewer misassemblies. This QC measure holds true even when ignoring misassemblies in centromeres and in segmental duplications and after correction for structural variants (Suppl. Table 2). Where larger gaps outside centromere regions occurred, we complemented this assembly with sequence from the `FALCON`-based assembly (Suppl. Table 3) to obtain a final Egyptian meta-assembly, denoted as EGYPT (for overall assembly strategy, see Suppl. Fig. 1). Both the base assemblies and the final meta assembly are of high quality and complementary and they are comparable to the publicly available assemblies of a Korean ^2^ and a Yoruba (GenBank assembly accession GCA_001524155.4) individual in terms of genome length and various quality control (QC) measures, (Table 1, extended version in Suppl. Table 2). Assembly quality is confirmed by quality control (QC) measures assessed by QUAST-LG ^33^ (Suppl. Table 2), NA-values (Suppl. Fig. 2), k-mer multiplicity with KAT ^34^ (Suppl. Fig. 3, 4 and 5), QV values of more than 40 and by dot plots of alignment with reference GRCh38 (Suppl. Figs. 6-10). We performed repeat annotation and repeat masking for all assemblies (Suppl. Table 4).

**Table 1:**
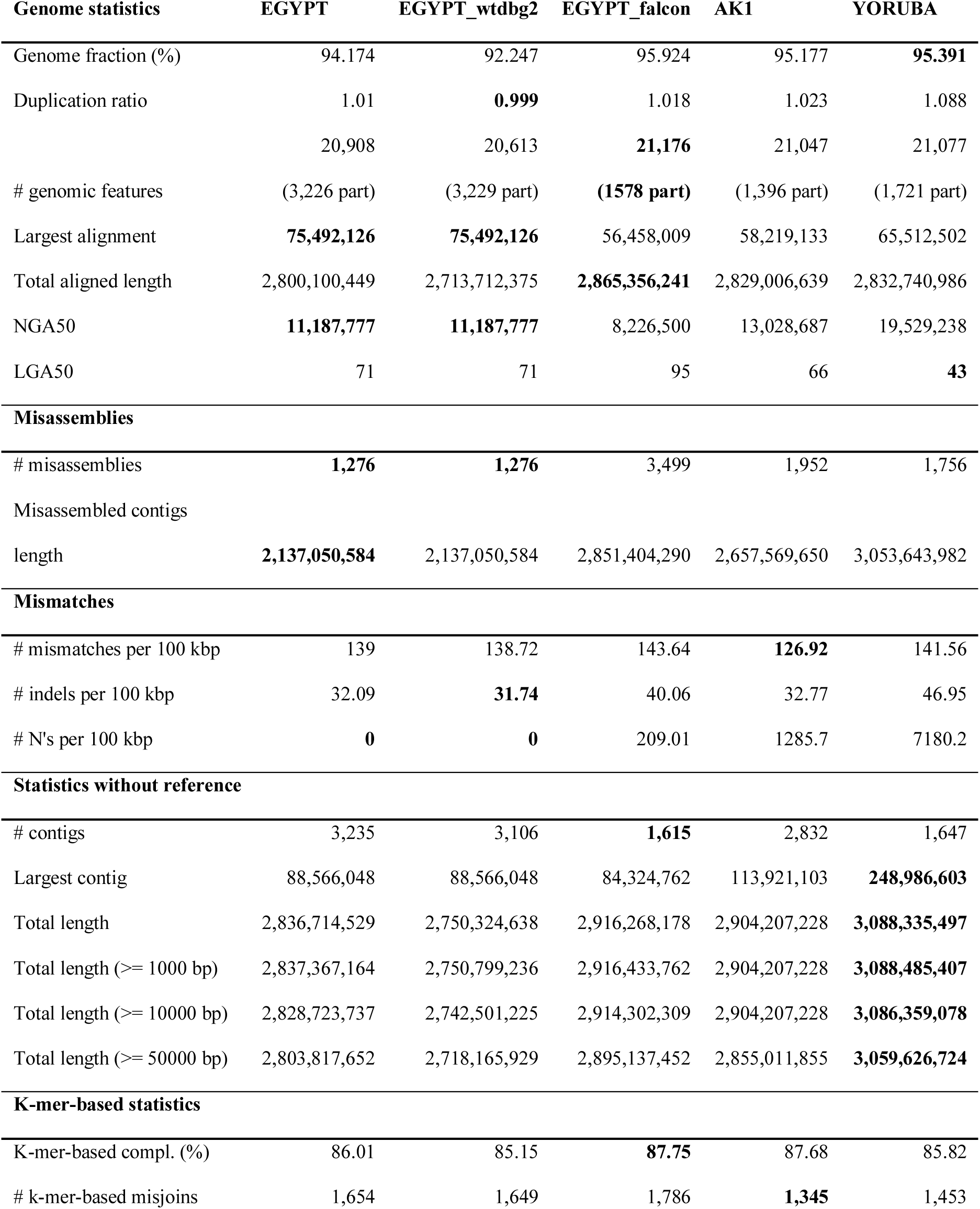
Default assembly quality measures according to `QUAST-LG`. The extended `QUAST-LG` report is provided in Suppl. Table 2. Yoruba is a chromosome-level assembly. Best quality for every measure is denoted in bold.

The meta-assembly was complemented with high-quality phasing information (Suppl. Table 5). EGYPT SNVs and small insertions and deletions (indels) called using short-read sequencing data were phased using high-coverage 10x linked-read sequencing data. This resulted in 3,834,900 of 4,008,080 autosomal variants being phased (95.7%). Furthermore, nearly all (99.41%) of the genes with lengths less than 100 kb and more than one heterozygous SNP were phased into a single phase block. We identified 22 runs of homozygosity (ROH) (Suppl. Table 6), out of which 16 are larger than 5 Mb and sum up to overall 192 MB, which indicates consanguinity at the level of parental third-degree relationship ^35^.

Based on the personal Egyptian genome, we constructed an Egyptian population genome by considering genome-wide SNV AFs in 109 additional Egyptians (Suppl. Table 7). This approach enabled the characterization of the major allele (i.e., the allele with highest AF) in the given Egyptian cohort. To accomplish this, we called variants using short-read data of 13 Egyptians sequenced at high coverage and 97 Egyptians sequenced at low coverage. Although sequence coverage affects variant-based statistics (Suppl. Fig. 11), due to combined genotyping, most variants could also be called reliably in low coverage samples (Suppl. Fig. 12). We called a total of 19,758,992 SNVs and small indels (Suppl. Fig. 13) in all 110 Egyptian individuals (Fig. 1). The number of called variants per individual varied between 2,901,883 to 3,934,367 and was correlated with sequencing depth (see Suppl. Figs. 11-12). This relationship was particularly pronounced for low coverage samples. The majority of variants were intergenic (53.5%) or intronic (37.2%) (Suppl. Fig. 14). Only approximately 0.7% of the variants were located within coding exons, of which 54.4% were non-synonymous and thus cause a change in protein sequence and, possibly, structure (Suppl. Fig. 15).

**Figure 1:**
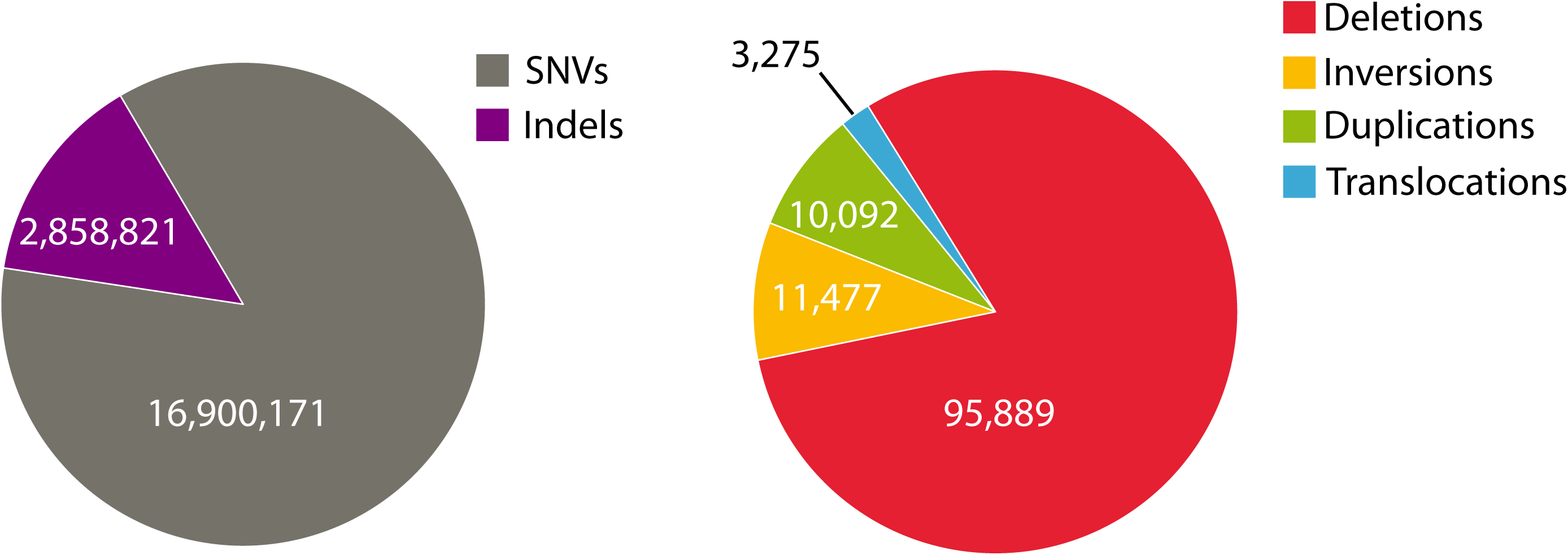
Number of various genetic variant types identified in the Egyptian cohort. Left: The number of SNVs and indels. Right: The number of SV calls: deletions, inversions, duplications and translocations. Additionally, 408 insertions have been called.

Using short-read sequencing data of 110 Egyptians, we called 121,141 SVs, most of which were deletions, but also included inversions, duplications, insertions and translocations of various orders of magnitude (Fig. 1, Suppl. Fig. 16-17). Similar to SNVs, the number of SV calls also varied between individuals (Suppl. Fig. 18) and is slightly affected by coverage (Suppl. Fig. 19). After merging overlapping SV calls, we obtained an average of 2,773 SVs per Egyptian individual (Suppl. Table 8, Suppl. Figs. 20-22).

Using the EGYPT *de novo* assembly, we searched for unique insertions that are common in Egyptians. Towards this, we first mapped all short-read data against the GRCh38 reference genome and to other decoy or alternative haplotype sequences from the GATK bundle. All reads that could not be mapped were subsequently mapped against the EGYPT *de novo* assembly. A similar approach was recently applied to identify novel, unique insertions in *de novo* assemblies of 17 individuals from 5 populations using 10x genomics sequencing ^36^. Altogether we identified 40 unique insertions longer than 500 bp with a total length of 40kb, for which we required for every base in the identified region to have a minimal coverage of 5 reads in at least 10 Egyptian individuals (Suppl. Table 9). Of these sequences, 28 have been mentioned before by Wong *et al*. ^36^, and 10 more in different studies within the last 15 years ^37 38 39 40^. Two out of the 40 insertions are most likely novel. In addition, one region contains three unique insertions, of which two contain additional, novel sequences longer than 500 bases. Closer inspection reveals that these sequences are located within a region of two 50 kb gaps (i.e. N sequences) in the GRCh38 reference genome at chromosome loci chr13:111,703,856-111,753,855 and chr13:111,793,442-111,843,441 with about 40 kb of reference sequence between the gaps. The EGYPT, AK1 and YORUBA assembly sequences that cover this 140 kb reference sequence from chr13:111,703,856 to 111,843,441 are very similar (Suppl. Figs. 23, 24 and 25). They all align about 4 kb from the 40 kb reference sequence between the gaps, only, but at the very beginning of the respective assembly sequence (Suppl. Figs. 26, 27 and 28). Performing a BLAST search of the 140 kb EGYPT assembly sequence reveals an overall 44 kb alignment in five, mainly consecutive, large alignment blocks to “Homo sapiens chromosome 13 clone WI2-2182D8” (AC188786.1) from position 1 to 44,382, see Suppl. Fig. 29. This large reference genome region that contains the largest gap covering sequence reported for AK1 ^2^ is not resolved yet.

Overall, we identified 330 single nucleotide variants and indels in 36 of 40 non-reference sequences (Suppl. Table 10). The percentage of reads that could not be mapped to GRCh38 or GATK bundle sequences, but which were mappable against the *de novo* assembly is on average 8.6%, but for some individuals up to 34.2% (cv. Suppl. Fig. 30). Previously unmapped short reads of 110 Egyptians covered positions for more than 19 Mb of the Egyptian *de novo* assembly. Unique sequences that are commonly shared among Egyptians illustrate that additional reference genomes are needed to capture the genetic diversity that are neither assessable by short read sequencing nor with the current human reference genome.

In addition, the large number of assembly positions to which such short reads map which could not be mapped to the reference genome GRCh38 (including widely used supplementary sequences included in the `GATK` bundle), indicate a need for further assembly-based reference data and for new approaches to better capture genetic diversity.

Genotype principal component analysis of the Egyptian cohort shows a homogeneous group for which the assembly individual is representative (Suppl. Figs. 31-37). We genetically characterized the Egyptian population with respect to 143 other populations of the world using variant data of 5,429 individuals in total. For this, we combined five different data sets: (1) a recently published whole genome sequencing (WGS)-based variant data set from 929 individuals of the Human Genome Diversity Project (HGDP), covering 51 populations ^41^; (2) 2,504 individuals from 26 populations of the 1000 Genomes project for which phase 3 genotypes are available ^5^; (3) WGS-based variant data from 108 Qatari individuals ^42^; (4) SNP array-based variant data of 478 individuals from five countries of the Arabian Peninsula ^25^; (5) 1,305 individuals from 68 African, European, Western and Southern Asian populations that were compiled from 8 different publications into a recent SNP array-based variant data set ^43^. All individuals and their annotations are provided in Suppl. Table 11, data sources are described in Suppl. Table 12. A principal component analysis of the data shows a genetic continuum between Europeans, Africans, East Asians and Americans along the first three principal components, see Suppl. interactive HTML-based Fig. PCA_interactive.html. Egyptians are located on the European-African axis and close to Europeans. Their genetic variance spreads to a small degree in the direction of the Asian axis, akin to further individuals from the Middle East (see Fig. 2c). To preclude a technical bias when intersecting WGS with SNP array data, we compared the analysis results when using whole genome data, only, or when intersecting WGS data with SNP arrays and found comparable results in both cases (Suppl. Fig. 38). The Egyptian PCA location is further supported by an admixture analysis. Our analysis specifies k=24 as the optimal number of genetic components for the entire data set, i.e. having the smallest cross validation error (see Suppl. Fig. 39 for results for k=10 to k=25). Accordingly, the genetics of Egyptian individuals comprises four distinct population components that sum up to 75% on average. Egyptians have a Middle Eastern, a European / Eurasian, a North African and an East African component with 27%, 24%, 15% and 9% relative influence, respectively (see Fig. 2a). According to our cohort, Egyptians show genetically little heterogeneity, with little variance in the proportion of individual components between the individuals (Suppl. Figs 40 and 41). With a focus on populations from the Horn of Africa, the four components we identified have been described before by Hodgeson *et al*. ^44^ in a cohort of 2,194 individuals from 81 populations (mainly 1000 Genomes and HGDP) and substantially fewer variants (n=16,766), but including also 31 Egyptians. They and others hypothesize that most non-African ancestry, i.e. the Eurasian / European and Middle Eastern components in the populations from North Africa and the Horn of Africa is resulting from prehistoric back-to-Africa migration ^44 24^. Recently, Serra-Vidal *et al*. describe North Africa as a “melting pot of genetic components”, attributing most genetic variation in the region also to prehistoric times ^45^. Here, we confirm previously identified genetic components, yet using 2.5 times as many individuals, and using WGS data for the majority of them. This is thus the hitherto most comprehensive data set on genetic diversity world-wide and in this region.

**Figure 2:**
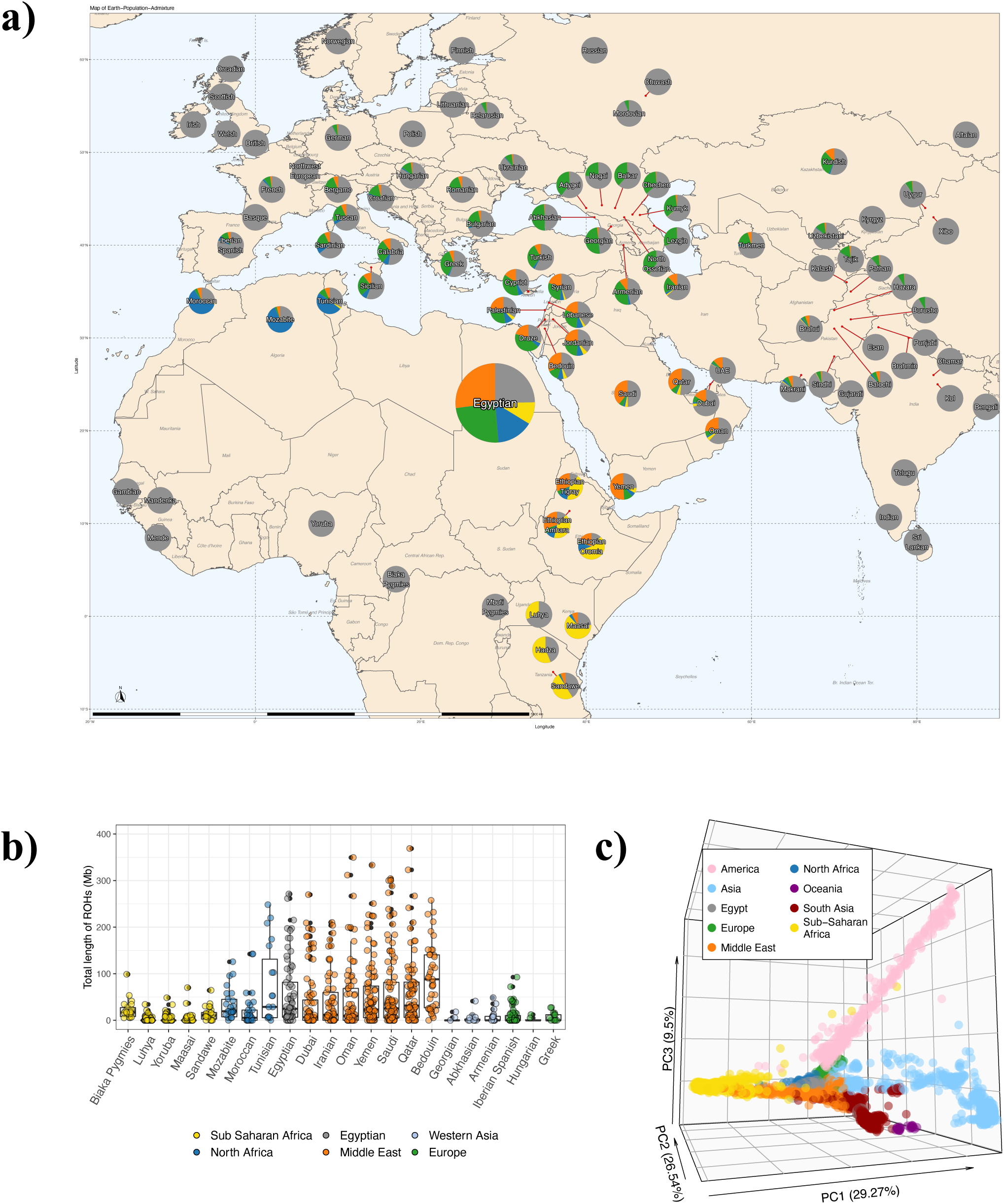
Population genetic characterization of the Egyptian population a) The four largest admixture components in the Egyptian population for African, European and Western Asian populations. b) Box plots for total length of runs of homozygosity for the Egyptians and several populations from relevant world regions (one Qatari not shown). c) Principal component analysis with individuals from populations world-wide.

The European, African and Asian ancestry components of Egyptians are further supported by mitochondrial haplogroup assessment from mtDNA sequencing of 227 individuals in additiona to 100 available from the literature ^27^. mtDNA sequencing revealed that Egyptians have haplogroups most frequently found in Europeans (e.g., H, V, T, J, etc.; >60%), African (e.g., L with 24.8%) or Asian/East Asian haplogroups (e.g., M with 6.7%). Overall, this supports the admixture and PCA analysis and the notion that Egypt’s transcontinental geographical location shaped Egyptian genetics (Suppl. Fig. 42).

Lastly, we characterized the Egyptian population with respect to runs of homozygosity. The distribution of overall length of ROHs larger than 5 Mb is comparable for the Egyptian population and Middle Eastern populations and, to lesser extent, also for other North African and Western Asian populations. In comparison, Europeans and Sub-Saharan Africans have usually shorter ROHs, see Fig. 2b. Abundance of long ROHs is typical for the Greater Middle East ^26^ and reflects the common practice of consanguineous marriages in this region.

In total, we identified 6,599,037 common Egyptian SNVs (minor allele frequency (MAF) > 5%, genotypes in a minimum of 100 individuals), of which 1,198 are population-specific; i.e., they are either rare (MAF < 1%) or not detected in any other population in the 1000 Genomes ^5^, gnomAD database ^6^ or TOPMed ^7^ (Suppl. Table 13). These numbers are comparable to population-specific variant numbers reported previously for 1000 Genomes populations ^46^. Four SNVs likely have a molecular impact (Suppl. Table 14), indicated by a CADD ^47^ deleteriousness score greater than 20. SNP rs143563851 (CADD 24.2) has recently been identified in 1% of individuals of a cohort of 211 Palestinians in a study that performed targeted sequencing of blood group antigen synthase GBGT1 ^48^. SNP rs143614333 (missense variant in gene CR2, CADD 23.6) is in ClinVar ^49^, with three submitters reporting that the variant is of uncertain clinical significance. Additionally, we obtained 49 variants with no dbSNP ^50^ rsID (Suppl. Table 15). These numbers of population-specific SNPs, of which some are likely to have an immediate impact on clinical characteristics and diagnostics, indicate insufficient coverage of the genetic diversity of the world’s population for precision medicine and thus the need for local genome references. To detect a putative genetic contribution of Egyptian population-specific SNPs towards molecular pathways, phenotypes or disease, we performed gene set enrichment analysis for all 461 protein-coding genes that were annotated to population-specific SNPs by Ensembl VEP ^51^. Enrichr, a gene list enrichment tool incorporating 153 gene sets and pathway databases ^52^, reports that genes from obesity-related traits of the GWAS catalog 2019 collection are over-represented (adj. p-value: 1.02E-6; 49 of 804 genes), which might hint at population-specific metabolism regulation that is linked to body weight.

Variants that are not protein coding may have a regulatory effect that affects gene and eventually protein expression. Using blood expression data obtained from RNA sequencing for the EGYPT assembly individual in conjunction with 10x sequencing-based phased variant data, we identified genes with haplotype-dependent expression patterns (see Suppl. Fig. 43 for the analysis overview and Suppl. Figs. 44-45 for the results). We report 1,180 such genes (Suppl. Table 16). Of these, variants contained in haplotypes of 683 genes (58%) have previously reported expression quantitative trait loci (eQTLs) in blood according to Qtlizer ^53^, for 380 genes supported by multiple studies. For 370 genes (31%), the strongest associated blood eQTL SNV is haplotypically expressed, and for 131 genes, the best eQTL has been previously reported by multiple studies. Concordance of haplotypic expression with eQTLs indicates that a common variant may affect gene expression; discordance hints towards a rare variant.

We investigated the impact of Egyptian ancestry on disease risk by integrating Egyptian variant data with the GWAS catalog ^54^, a curated database of GWAS. According to the GWAS catalog, most published GWAS are performed on Europeans ^29^, and only a single study has been performed on Egyptians ^55^ (by one of the co-authors). Furthermore, only 2% of individuals included in GWAS are of African ancestry ^29^. AFs, LD and genetic architecture can differ between populations, such that results from European GWAS cannot necessarily be transferred ^30^. This lack of transferability also compromises the prediction of an individual’s traits and disease risk using polygenic scores: such scores are estimated to be approximately one-third as informative in African individuals compared to Europeans ^56^. From the GWAS catalog, we constructed a set of 4,008 different, replicated, high-quality tag SNPs (i.e., one strongest associated SNP per locus) from European ancestry GWAS for 584 traits and diseases. We compared the tag SNPs’ AFs and proxy SNPs in the Egyptian cohort (n=110) and Europeans from 1000 Genomes (n=503) (Suppl. Table 17). Egyptian AFs of tag SNPs are comparable to European AFs, with a tendency to be lower (Fig. 3a). There are variants common in Europeans (AF>5%) but rare in Egyptians (AF<5%) (Suppl. Fig. 46). A total of 261 tag SNPs are not present in the Egyptian cohort (∼7%), clearly indicating a need to perform GWAS in non-European populations to further elucidate disease risk conferred by these loci. We investigated differences in LD structure using an approach that is used for fine-mapping of GWAS data, which identifies proxy variants (illustrated in Fig. 3c). Proxy variants are variants correlated with the tag GWAS SNP, i.e., in high LD (here, R^2^>0.8). The post-GWAS challenge is the identification of a causal variant from a set of variants in LD (tag SNP and proxy variants). We found that the number of proxy variants was much lower in the Egyptian cohort (Fig. 3b), likely due to shorter haplotype blocks known from African populations. This indicates that LD differences between Egyptians and Europeans may compromise GWAS transferability and European ancestry-based polygenic scores. However, Egyptian proxy variants are usually included in the larger set of European proxy variants (Fig. 3d). An example is variant rs2075650 (a locus sometimes attributed to gene TOMM40), which has been linked to Alzheimer’s disease in seven GWASs (cf. Suppl. Fig. 47). This tag SNP has seven proxy variants in Europeans but only two proxy variants in Egyptians. One European proxy, rs72352238, has also been reported as a GWAS tag SNP, but it is not a proxy of rs2075650 in Egyptians and may thus fail replication and transfer of GWAS results from the European to the Egyptian population.

**Figure 3:**
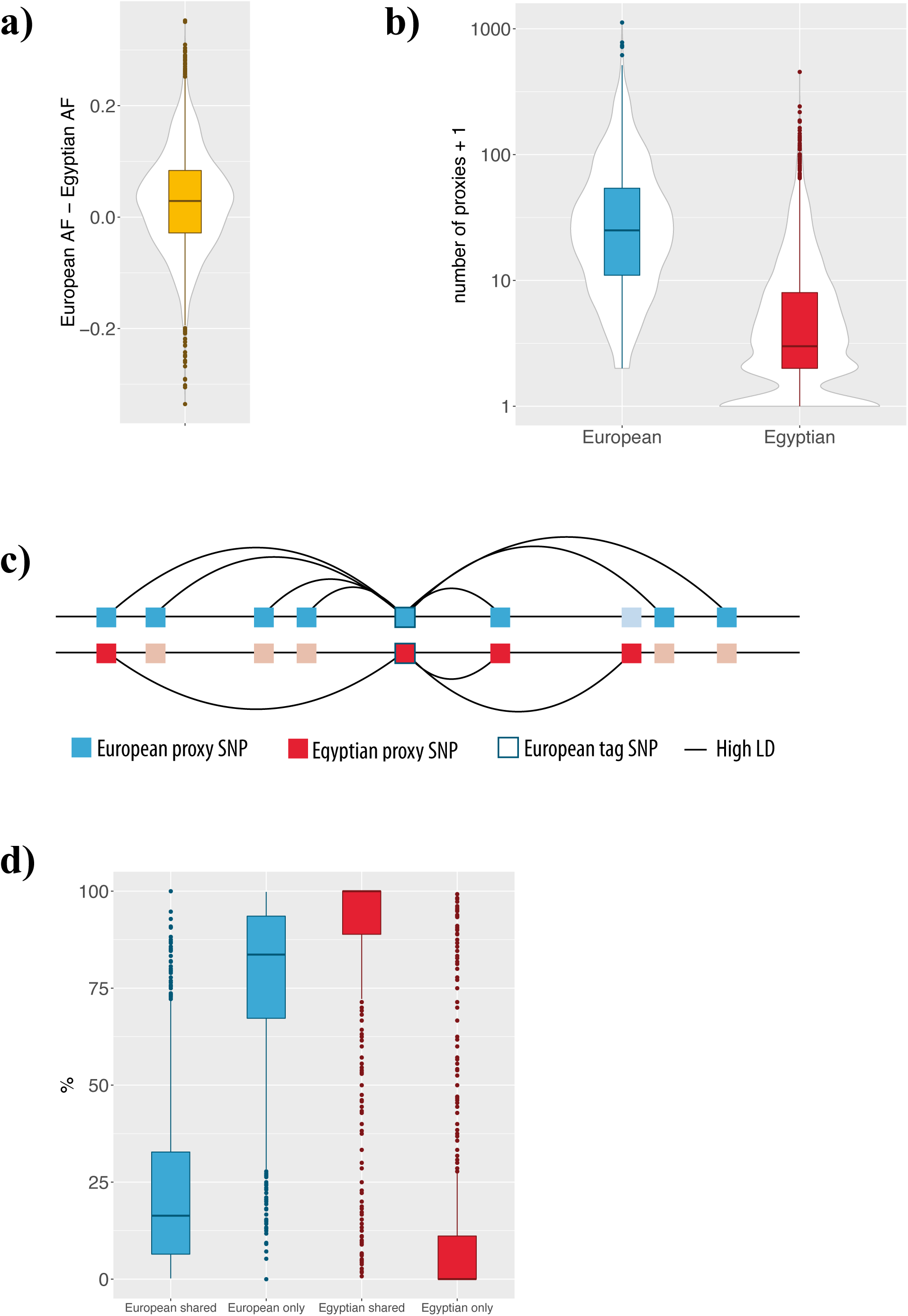
AF and proxy SNP comparisons for 3,698 GWAS tag SNPs called in a minimum of 100 Egyptians. a) AF differences. b) Number of proxies. c) Illustration of the proxy SNP comparison. A European GWAS tag SNP (center) and variants in Europeans (top) and Egyptians (bottom). Lines denote variants in high LD. The tag SNP has 7 proxy variants in Europeans and 3 in Egyptians. Light blue/red variants are no proxy variants in Europeans/Egyptians. Two proxy variants are shared. Thus 2 of 7 European (∼29%) and 2 of 3 Egyptian (∼67%) variants are shared. Further 5 of 7 European proxies are European-only (∼71%) and 1/3 Egyptian proxies are Egyptian-only (∼33%). d) European shared: Percentage of European proxy SNPs shared with Egyptian proxy SNPs. European only: Percentage of European proxy SNPs not shared with Egyptian proxies. Egyptian shared / Egyptian only respectively.

With our Egyptian genome reference, it will be possible to perform comprehensive integrated genome and transcriptome comparisons for Egyptian individuals in the future. This will shed light on personal as well as population-wide common genetic variants. As an example for personalized medicine for Egyptian specific genetics we visualize the complete genetic information of the DNA repair-associated gene BRCA2 from our study in the integrative genomics viewer ^57^ (IGV) and the variant phasing information within the 10x Genomics browser `LOUPE` in Fig. 4 and Suppl. Fig 48, respectively. BRCA2 is linked to the progression and treatment of breast cancer and other cancer types ^58^, if mutated. The IGV depicts the sample coverage based on sequencing data from PacBio, 10x Genomics and Illumina (whole genome as well as RNA) for the personal EGYPT genome together with common Egyptian SNPs. Variants previously assessed in a breast cancer GWAS ^58^ are displayed as Manhattan plot; note the three significant GWAS SNPs between positions 32,390 and 32,400 kb. The bottom compares the identified SNVs and indels from the Korean and Yoruba *de novo* assembly with our *de novo* EGYPT assembly. Visual inspection of both small and structural variations at the personal and population-based genome levels already yields significantly different variants, which might be important for genetic counselling and detection of inherited risks for cancer.

**Figure 4:**
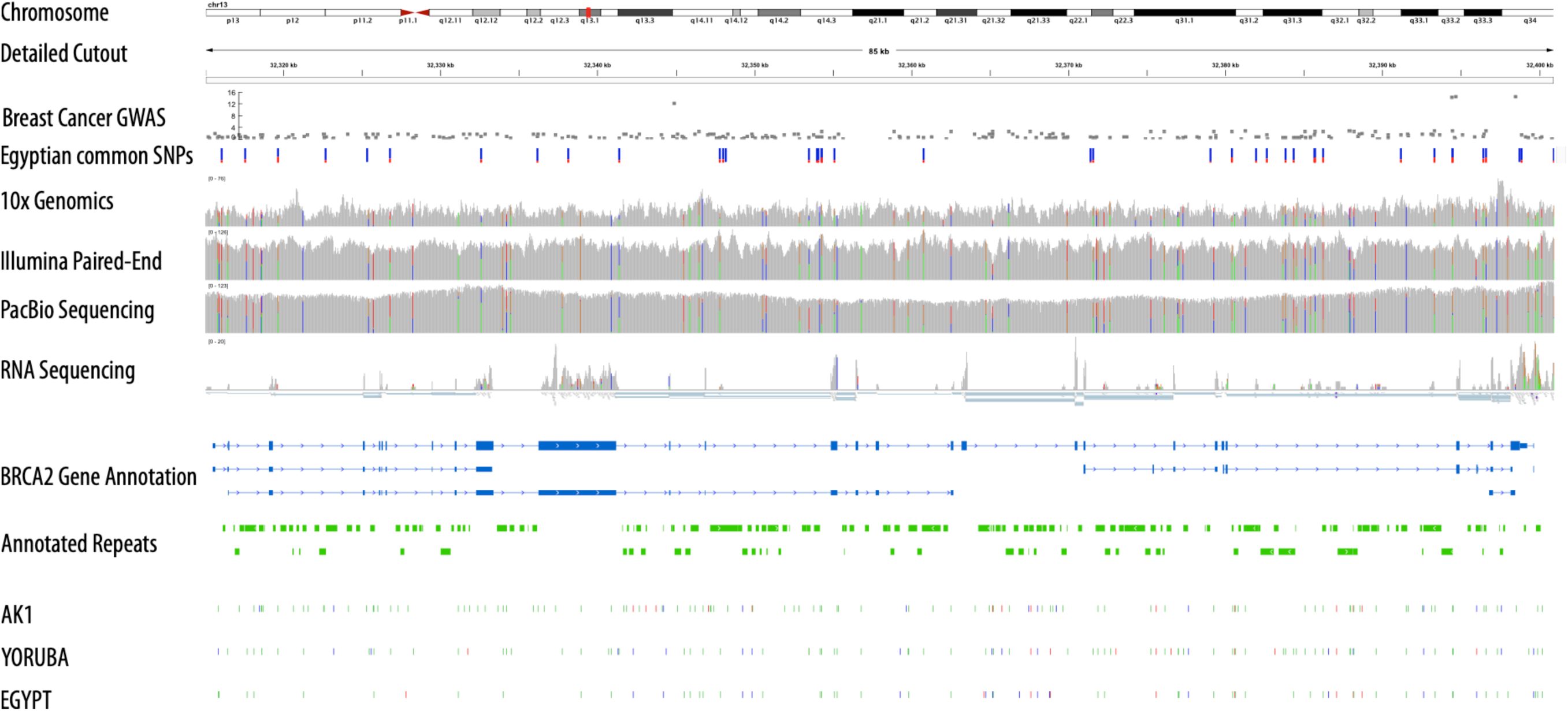
Integrative view of Egyptian genome reference data for the gene BRCA2, which is associated with breast cancer. The rows denote from top to bottom: Genome location on chromosome 13 of the magnified region for BRCA2 (first and second row); GWAS data for breast cancer risk ^58^; Variants that are common in the cohort of 110 Egyptians; Read coverage of genetic region based on 10x Genomics, Illumina paired-end and PacBio sequencing data; Coverage and reads of RNA sequencing data; BRCA2 gene annotation from Ensembl; Repeats annotated by; SNVs and indels identified by comparison of assemblies AK1, YOURUBA and EGYPT with GRCh38. The colors denote base substitutions (green), deletions (blue) and insertions (red). The corresponding variant phasing for the EGYPT individual is displayed in Suppl. Fig. 48.

In conclusion, we constructed the first Egyptian – and North African – genome reference, which is an essential step towards a comprehensive, genome-wide knowledge base of the world’s genetic variations. The wealth of information it provides can be immediately utilized to study in-depth personal genomics and common Egyptian genetics and its impact on molecular phenotypes and disease. This reference will pave the way towards a better understanding of the Egyptian, African and global genomic landscape for precision medicine.

## Methods

### Sample acquisition

Samples were acquired from 10 Egyptian individuals. For nine individuals, high-coverage Illumina short-read data were generated. For the assembly individual, high-coverage short-read data were generated as well as high-coverage PacBio data and 10x data. Furthermore, we used public Illumina short-read data from 100 Egyptian individuals from Pagani *et al*. ^27^. See Supplementary Tables 1 and 7 for an overview of the individuals and the corresponding raw and result data generated in this study.

### PacBio data generation

For PacBio library preparation, the SMRTbell DNA libraries were constructed following the manufacturer’s instructions (Pacific Bioscience, www.pacb.com). The SMRTbell DNA libraries were sequenced on the PacBio Sequel and generated 298.2 GB of data. Sequencing data from five PacBio libraries were generated at overall 99x genome coverage.

### Illumina short-read data generation

For 350 bp library construction, the genomic DNA was sheared, and fragments with sizes of approximately 350 bp were purified from agarose gels. The fragments were ligated to adaptors and amplified using PCR. The generated libraries were then sequenced on the Illumina HiSeq X Ten using PE150 and generated 312.8 GB of data.

For the assembly individual, sequencing data from five libraries was generated at overall 90x genome coverage. For nine additional individuals, one library each was generated, amounting to an overall 305x coverage of sequencing data. For the 100 individuals of Pagani *et al*. ^27^, three were sequenced at high coverage (30x) and 97 at low coverage (8x). The average coverage over SNV positions for all 110 samples is provided in Supplementary Table 7.

### RNA sequencing data generation

For RNA sequencing, ribosomal RNA was removed from total RNA, double-stranded cDNA was synthesized, and then adaptors were ligated. The second strand of cDNA was then degraded to generate a directional library. The generated libraries with insert sizes of 250-300 bp were selected and amplified and then sequenced on the Illumina HiSeq using PE150. Overall, 64,875,631 150 bp paired-end sequencing reads were generated.

### 10x sequencing data generation

For 10x genomic sequencing, the Chromium Controller was used for DNA indexing and barcoding according to the manufacturer’s instructions (10x Genomics, www.10xgenomics.com). The generated fragments were sheared, and then adaptors were ligated. The generated libraries were sequenced on the Illumina HiSeq X Ten using PE150 and generated 272.7 GB of data. Sequencing data from four 10x libraries was generated at overall 80x genome coverage.

### Construction of draft *de novo* assemblies and meta-assembly

We used `WTDBG2` ^32^ for human *de novo* assembly followed by its accompanying polishing tool `WTPOA-CNS` with PacBio reads and in a subsequent polishing run with Illumina short reads. This assembly was further polished using PILON ^59^ with short-read data (cf. Suppl. Methods: *-based assembly*). An alternative assembly was generated by using FALCON ^60^, QUIVER ^61^, SSPACE-LONGREAD ^62^, `PBJELLY` ^63^, `FRAGSCAFF` ^64^ and PILON ^59^ (cf. Suppl. Methods: *-based assembly*).

Proceeding from the `WTDBG2`-based assembly, we constructed a meta-assembly. Regions larger than 800 kb that were not covered by this base assembly and were not located within centromere regions were extracted from the alternative `FALCON`-based assembly (Suppl. Table 3). See Suppl. Fig. 1 for an overview of our assembly strategy, including meta-assembly construction (cf. Suppl. Methods: *Meta-assembly construction*).

Assembly quality and characteristics were assessed with QUAST-LG ^33^. Additionally, we removed misassemblies in centromeres or in segmental duplication regions from the `QUAST-LG` report and furthermore removed structural variants from misassemblies (cf. Suppl. Methods: *Assembly comparison and QC*). The extraction of coordinates for meta-assembly construction was performed using `QUAST-LG` output. K-mer multiplicity was assessed with KAT ^34^. Following Porubsky *et al*. ^65^, we computed QV as the number of homozygous variants divided by the effective genome size. Towards this, we mapped all short reads to the assembly using `BWA MEM` and perform variant calling using `FREEBAYES` with default parameters. We kept only homozygous variants with a minimum quality of 10 using `VCFTOOLS`. Single-nucleotide differences were counted as difference of 1 bp, indel differences as the length differences between reference and alternative allele. Based on `SAMTOOLS` command “stats”, we computed the sum of bases with short read coverage as effective genome size.

### Repeatmasking

Repeatmasking was performed by using REPEATMASKER ^66^ with RepBase version 3.0 (Repeatmasker Edition 20181026) and Dfam_consensus (http://www.dfam-consensus.org) (cf. Suppl. Methods: *Repeat annotation*).

### Unique inserted sequences

We trimmed Illumina short sequencing reads of 110 Egyptian individuals using `FASTP` 0.20.0 with default parameters, mapped the output reads to GRCh38 and `GATK` bundle sequences using `BWA` 0.7.15-r1140 and sorted by chromosomal position using `SAMTOOLS` 1.3.1. Subsequently, we extracted reads that did not map to GRCh38 using `SAMTOOLS` with parameter F13 (i.e. read paired, read unmapped, mate unmapped) and repeated the mapping and sorting using the Egyptian *de novo* assembly. We merged the read-group specific BAM files for each sample and calculated the per base read depth using `SAMTOOLS`. Afterwards, we aggregated the results via custom scripts and extracted uniquely inserted sequences from the Egyptian *de novo* assembly. Insertions were defined as contiguous regions of at least 500 bp having a coverage of more than 5 reads per base in 10 or more samples. Lastly, we `BLAST`ed the obtained sequences against the standard databases (option nt) for highly similar sequences (option megablast) using a custom script. For the uniquely inserted sequences that we identified, we created a pileup over all BAM files containing the reads that did not map to GRCh38 using `SAMTOOLS`. Based on these pileups, we then called the variants using `BCFTOOLS`. Variants with quality of more than 10 were kept.

### Phasing

Phasing was performed for the assembly individual’s SNVs and short indels obtained from combined genotyping with the other Egyptian individuals, i.e., based on short-read data. These variants were phased using 10x data and the 10x Genomics `LONGRANGER WGS` pipeline with four 10x libraries provided for one combined phasing. See Supplementary Methods *Variant phasing* for details.

### SNVs and small indels

Calling of SNVs and small indels was performed with `GATK` 3.8 ^67^ using the parameters of the best practice workflow. Reads in each read group were trimmed using `Trimmomatic` ^68^ and subsequently mapped against reference genome hg38 using `BWA-MEM` ^69^ version 0.7.17. Then, the alignments for all read groups were merged sample-wise and marked for duplicates. After the base recalibration, we performed variant calling using `HaplotypeCaller` to obtain GVCF files. These files were input into GenotypeGVCFs to perform joint genotyping. Finally, the variants in the outputted VCF file were recalibrated, and only those variants that were flagged as “PASS” were kept for further analyses. We used `FastQC` ^70^, `Picard Tools` ^71^ and `verifyBamId` ^72^ for QC (cf. Suppl. Methods: *Small variant QC*).

### Variant annotation

Variant annotation was performed using `ANNOVAR` ^73^ and `VEP` ^51^ (cf. Suppl. Methods: *Small variant annotation*)

### Structural variants

SVs were called using `DELLY2` ^74^ with default parameters as described on the `DELLY2` website for germline SV calling (https://github.com/dellytools/delly) (cf. Suppl. Methods: *Structural variant QC*). Overlapping SV calls in the same individual were collapsed by the use of custom scripts. See Supplementary Methods *Collapsing structural variants* for details.

### Population genetics

For population genetic analyses, we compared the Egyptian variant data with variant data from five additional sources specified in Suppl. Table 12. Individuals together with their annotations are listed in Suppl. Table 11. Variant data was merged to contain only variants present in all data sets and subsequently filtered and LD pruned. Genotype principal component analysis was computed using SMARTPCA ^75^ from the `EIGENSOFT` package. Admixture was computed with ADMIXTURE ^76^ (cf. Suppl. Methods: *Population genetics* and *SNP array-based Egyptian variant data*). Runs of homozygosity were computed on the same files that were used for PC computation and admixture using `PLINK –homozyg`. ROHs with size larger than 5 Mb were summed to obtain overall length of ROHs per individual.

### Mitochondrial haplogroups

Haplogroup assignment was performed for 227 individuals using `HAPLOGREP` 2 ^77^. Furthermore, mitochondrial haplogroups were obtained from Pagani *et al*. ^27^ for 100 individuals. See Supplementary Methods *Mitochondrial haplogroups* for details.

### Population-specific variants

Our set of common Egyptian SNVs comprises variants with genotypes in a minimum of 100 individuals whose alternative allele has a frequency of more than 5%. Those common Egyptian SNVs that are otherwise rare, i.e., have an AF of less than 1% in the 1000 Genomes, and gnomAD populations as well as in TOPMed were considered Egyptian-specific. AFs were annotated using the Ensembl API. Furthermore, a list of Egyptian common variants without dbSNP rsID was compiled, see Supplementary Methods *Small variant annotation* for details.

### Haplotypic expression analysis

RNA sequencing reads were mapped and quantified using `STAR` (Version 2.6.1.c) ^78^. Haplotypic expression analysis was performed by using `PHASER` and PHASER GENE AE (version 1.1.1) ^79^ with Ensembl version 95 annotation on the 10x-phased haplotypes using default parameters. See Supplementary Methods *Haplotypic expression* for details.

### GWAS catalog data integration

GWAS catalog associations for GWAS of European ancestry were split into trait-specific data sets using Experimental Factor Ontology (EFO) terms. For every trait, a locus was defined as an associated variant +/-1 MB, and only loci that were replicated were retained. For proxy computation, we used our Egyptian cohort (n=110) and the European individuals of 1000 Genomes (n=503). For details, see Supplementary Methods *Data integration with the GWAS catalog*.

### Integrative genomics view

We implemented a workflow to extract all Egyptian genome reference data for view in the IGV ^57^. This includes all sequencing data mapped to GRCh38 (cf. Suppl. Methods *Sequencing read mapping to GRCh38*) as well as all assembly differences (cf. Suppl. Methods *Alignment to GRCh38* and *Assembly-based variant identification*) and all Egyptian variant data. See Supplementary Methods *Gene-centric integrative data views* for details.

### Ethics statement

This study was approved by the Mansoura Faculty of Medicine Institutional Review Board (MFM-IRB) Approval Number RP/15.06.62. All subjects gave written informed consent in accordance with the Declaration of Helsinki. This study and its results are in accordance with the Jena Declaration (https://www.uni-jena.de/unijenamedia/Universit%C3%A4t/Abteilung+Hochschulkommunikation/Presse/Jenaer+Erkl%C3%A4rung/Jenaer_Erklaerung_EN.pdf).

## Supporting information

An_Egyptian_genome_reference_supplement_v2.pdf

An_Egyptian_genome_reference_supplementary_tables_v2.xlsx

PCA_interactive.html.zip

## Supplementary information

*Supplementary Tables 1-17:* An_Egyptian_genome_reference_supplementary_tables.xlsx *Supplementary Methods and Supplementary Figures 1-48:* An_Egyptian_genome_reference_supplement.pdf

## Acknowledgements

We acknowledge support on coordination of the project and assembly work w.r.t the `FALCON`- based assembly through Ms. Lu Wang from the Novogene (UK) Company Limited. IW, HB and SI acknowledge funding by the Deutsche Forschungsgemeinschaft (DFG, German Research Foundation) under Germany’s Excellence Strategy – EXC 22167-390884018. Verónica Calonga-Solís was supported by a DAAD scholarship. All authors acknowledge computational support from the OMICS compute cluster at the University of Lübeck.

## Contributions

H.B, S.I. and M.S. conceived the study. I.W, A.K, M.M., H.B. and S.I. designed the study. I.W., A.K., M.M., M.O, A.F. and V. C.-S. performed data analysis. C.M. constructed the `FALCON`- based assembly. M.S. and S.E-M. compiled the Egyptian cohort and provided samples. M.H. performed mtDNA library preparation and sequencing. I.W., H.B. and S.I. wrote the manuscript. All authors read and approved the final manuscript.

## Data availability

All summary data of the Egyptian genome reference are available at www.egyptian-genome.org, where also variant allele frequencies can be queried online. Raw sequencing data and variant data are available at EGA under study ID EGAS00001004303. De novo assemblies are available at NCBI under BioProject ID PRJNA613239.

## Code availability

Computational tools used are specified in the Supplementary Methods. Workflows use Snakemake and Conda (especially Bioconda) for reproducible data analysis and are provided on request.

## Corresponding authors

Correspondence to Hauke Busch or Saleh Ibrahim.

