## Supplementary material for "An integrated personal and population-based Egyptian genome reference": An_Egyptian_genome_reference_supplement_v2.pdf

*Inken Wohlers, Axel Künstner, Matthias Munz, Michael Olbrich, Anke Fähnrich, Verónica Calonga-Solís, Caixia Ma, Misa Hirose, Shaaban El-Mosallamy, Mohamed Salama, Hauke Busch* & Saleh Ibrahim**

* These authors contributed equally to this work

**Supplementary Figures** **12**

**Supplementary References** **53**

### Supplementary Methods

#### WTDBG2-based assembly

The WTDBG2-based assembly was constructed with WTDBG2 [1] using preset option "-x sq", which applies default options for PacBio sequencing data-based assembly and "-g 3g" to specify an approximate genome size of 3 Gb. WTPOA-CNS, a consensuser for WTDBG2, was applied as described on the WTDBG2 website. Briefly, WTPOA-CNS was applied once for obtaining a draft assembly, then for polishing with PacBio data (after mapping reads to the draft assembly with MINIMAP2 [2] and option "-x map-pb") and finally for polishing with Illumina short-read data using preset "-x sam-sr" for using default options for short read polishing (after mapping to the PacBio-polished assembly with BWA-MEM [3]). We remapped the short reads to this polished assembly using again BWA-MEM to apply another round of polishing using PILON [4], an established short-read polishing tool. This slightly improved the already very good overall assembly quality.

#### FALCON-based assembly

The FALCON-based assembly was generated with FALCON (version 0.7), followed by consensusing using QUIVER (version smrtlink_5.0.1). Program SSPACE-LONGREAD (version 1-1) was used to assemble to scaffolds and PBJELLY (version 15.8.24) to fill gaps. For scaffolding using 10x data, FRAGSCAFF (version 140324) was used. The assembly has been polished using PILON (version 1.22) [4].

#### Meta assembly construction

Centromere regions have been obtained from the UCSC Genome browser (URL https://bit.ly/2IYfpgW) by selecting "group: all tables" and "table: centromeres". GRCh38 coordinates of regions greater than 800 kb of the GRCh38 reference genome which were not covered by the WTDBG2-based assembly and were not located within centromere regions were identified and coordinates of aligned segments of the FALCON-based assembly which overlap these gaps were obtained. The aligned regions of the FALCON-based assembly were extracted and added to the WTDBG2-based assembly as individual, additional contigs. The considered assembly gaps and corresponding novel FALCON-based contigs which were added in the meta assembly are given in Suppl. Table 3. By filling 31 gaps, which affect 10 chromosomes, this approach slightly improves the GRCh38-covered genome fraction and the k-mer-based completeness of the meta assembly in comparison to the WTDBG2-based assembly (see also Suppl. Table 2).

### Assembly comparison and QC

We compared the assembly with the latest version of the reference genome assembly GRCh38, which we obtained from ftp://ftp.ensembl.org/pub/release-93/fasta/homo_sapiens/dna/Homo_sapiens.GRCh38.dna.primary_assembly.fa.gz. In the following, we refer to this reference assembly version as GRCh38.

For assembly quality assessment, we compared our two assemblies, EGYPT_wtdbg2 and EGYPT_falcon and the final meta assembly EGYPT with the published assembly of a Korean individual [5] obtained from https://www.ncbi.nlm.nih.gov/assembly/GCA_001750385.2 as well as a chromosome-level assembly of a Yoruba individual (1000 Genomes ID NA19420) downloaded from https://www.ncbi.nlm.nih.gov/assembly/GCA_001524155.4. We assessed all quality measures available by QUAST_LG [6] using default options and, as suggested for large genomes, the options "—large" and "—memory-efficient". We also computed k-mer based statistics using QUAST_LG with the option "—k-mer-stats". For QUAST_LG analysis concerning the number of genes contained in the assembly, we used gene annotation from Ensembl version 95 obtained from ftp://ftp.ensembl.org/pub/release-95/gff3/homo_sapiens/Homo_sapiens.GRCh38.95.gff3.gz. Interactive overview of QUAST_LG statistics as well as the ICARUS [7] contig viewer displaying all assemblies is provided at www.egyptian-genome.org. We used an approach utilized in two recent assembly publications [8] [9] to better characterize QUAST_LG misassemblies and correct for structural variants. With minor adaptations, we used the script of Shafin *et al.* [8] from https://github.com/kishwarshafin/helen/blob/master/helen/modules/python/helper/quast_sv_extractor.py, together with the centromere and segmental duplication files that the authors used, which were downloaded from https://www.ncbi.nlm.nih.gov/grc/human and https://github.com/mvollger/segDupPlots/, respectively. Structural variants that we removed were obtained from the DELLY short read-based SV calls for the assembly individual. For the AK1 assembly, we obtained structural variants from Suppl.Table 16 of Seo *et al.* [5]. Resulting stratified misassembly numbers for all assemblies are provided together with the QUAST_LG QC numbers in Suppl. Table 2.

### Repeat annotation

We performed repeat annotation with the tool REPEATMASKER version 4.0.7 (Smit *et al.* (2015)) using human repeat databases from Repbase (RepBaseRepeatMaskerEdition-20181026.tar.gz obtained from www.girinst.org) and Dfam_consensus (http://www.dfam-consensus.org, distributed with REPEATMASKER). REPEATMASKER was run with option "-q" for increased speed at slightly lower sensitivity and "-xsmall", which returns a file with sequences masked such that repetitive regions are denoted in lower case letters and non-repetitive regions in capital letters.

### Variant phasing

Variant phasing was performed based on 10x sequencing data from four libraries by using `LONGRANGER WGS` (version 2.2.2) with the GRCh38 reference files provided by 10x genomics at `http://cf.10xgenomics.com/supp/genome/refdata-GRCh38-2.1.0.tar.gz`. We have four 10x libraries available and barcodes may be shared between libraries, but denote different molecules. Thus, we adjusted the configuration MRO file of the 10x pipeline management framework Martian as described on the 10x genomics web site. This results in a suffix being appended to barcodes indicating the source library number.

### Small variant QC

We summarized and QC'ed SNV and small indels with respect to number of SNV, percentage of missing SNV calls, mean depth and heterozygosity per individual (Suppl. Fig. 12) to detect potential outliers. Sequencing depth did not substantially influence variant calling, but the aforementioned numbers are correlated with (and likely driven by) sequencing depth (Suppl. Fig. 11). Small indels are most frequent, and insertions of a given size have been called about as often as deletions, which is denoted by a symmetric size distribution (Suppl. Fig. 13).

### Small variant annotation

SNVs and small indels have been annotated using `ANNOVAR` [10] as well as `VEP` [11] and classified according to variant location (Suppl. Fig. 15). Exonic variants have further been classified according to variant consequence types (Suppl. Fig. 15). Deleterious effects of protein-coding variants were predicted using `CADD` [12], `POLYPHEN` [13] and `SIFT` [14] scores provided by the annotation tools. The stand-alone `VEP` tool used specifies the internally used data sources as VEP="v95" time="2019-04-12 09:14:39" ensembl-io=95.78ccac5 ensembl=95.4f83453 ensembl-funcgen=95.94439f4 ensembl-variation=95.858de3e 1000genomes="phase3" COSMIC="86" ClinVar="201810" ESP="V2-SSA137" HGMD-PUBLIC="20174" assembly="GRCh38.p12" dbSNP="151" gencode="GENCODE 29" genebuild="2014-07" gnomAD="170228" polyphen="2.2.2" refseq="2018-07-10 14:50:52 - GCF_000001405.38_GRCh38.p12_genomic.gff" regbuild="1.0" sift="sift5.2.2". Allele frequency information has been obtained via the Ensembl API (accessed 09/2019). The subset of Egyptian common variants, which has not been assigned an rsID has been annotated additionally with the newest available VEP version via the Ensembl web interface (https://www.ensembl.org/Tools/VEP). The data sources here are "v98" time="2019-10-24 12:37:40" cache="/nfs/public/release/ensweb-data/latest/tools/www/e98/vep/cache/homo_sapiens/98_GRCh38" db="homo_sapiens_core_98_38@hh-mysql-ens-species-web-1" 1000genomes="phase3" COSMIC="89" ClinVar="201907" ESP="V2-SSA137" HGMD-PUBLIC="20184" assembly="GRCh38.p13" dbSNP="152" gencode="GENCODE 32" genebuild="2014-07" gnomAD="r2.1" polyphen="2.2.2" regbuild="1.0" sift="sift5.2.2". This reduces the number of common Egyptian SNPs without rsID to 48 (cf. Suppl. Table 12).

### Structural variant QC

We computed for the 135,819 SVs called by `DELLY2` [15] in the cohort of 110 Egyptians the number of SVs, percentage of missing SV calls and heterozygosity per individual (see Suppl. Fig. 18). These numbers are correlated (Suppl. Fig. 19). Most of 121,141 `DELLY` filter passing SV calls thereof are deletions (n=95,889), but also inversions (n=11,477), duplications (n=10,092), translocations (n=3,275) and insertions (n=408) have been called. Corresponding SV lengths are up to $10^8$ with most SV calls affecting sequence up to 10kB (Suppl. Fig. 17).

### Collapsing structural variants

Structural variant calls of type deletion, insertion, inversion or duplication were collapsed per individual by dividing variants calls into groups with respect to their chromosomal region (chromosome, start position, end position). That is, each group only contains variant calls that are overlapping with each other and with none of the variant calls in any other group. Translocations were collapsed per individual by merging variants with the same original chromosomal position and the same new chromosomal position. The number of SVs per individual after collapsing and a boxplot of the corresponding SV sizes are provided in Suppl. Figs 20,21 and 22 for deletions, inversions and duplications.

### Population genetics

For PCA of 110 Egyptian individuals only (Suppl. Figs. 31-37), we kept biallelic variants with minor allele frequency greater 5% that do not violate Hardy-Weinberg equilibrium ($p = 10^{-6}$) and have no missing genotypes. Variants in high LD regions or known inversions have been removed (chr6:25 Mb–33.5 Mb, chr5:44 Mb–51.5 Mb, chr8:8 Mb–12 Mb, chr11:45 Mb–57 Mb) [16]. LD pruning was performed using `PLINK` with "–indep-pairwise 1000 10 0.2". For population genetics analyses with other populations, we used the cohort of 110 whole genome sequenced Egyptian individuals and 5,319 individuals from overall 143 other populations, compiled from five additional sources, Bergström *et al.* [17], 1000 genomes (1000G) [18], Fernandes *et al.* [19], Rodriguez-Flores *et al.* [20], and Busby 2020 (`http://dx.doi.org/10.17632/ckz9mtgrjj.3`) for details on these data sets see Suppl. Table 12. From the Qatar WGS data of Rodriguez-Flores *et al.*, we removed variants with filter flag LowQual followed by `PICARD` LiftoverVcf from hg37 to hg38. The array data from Fernandes *et al.* and Busby *et al.* was converted from plink to VCF format using plink with options ”–recode vcf-iid –real-ref-alleles –alleleACGT” followed by PICARD LiftoverVcf to GRCh38 (Busby *et al.* from hg36, Fernandes *et al.* from 37). From the Busby data set, 30 duplicate individuals were removed. We intersected the datasets using command ”bcftools isec -c none -n 11111” and merged with command ”bcftools merge –merge all”. From the merged data sets we kept all individuals, except five outlier in the Fernandes data set (individual IDs H9,IR134,IR84,N39,S60) and from the Busby data set we kept only the African, European, Western and South Asian populations which consist of individuals that are not part of the 1000G or Bergström data set already (68 Busby populations are added). Dataset inersection results in $n = 280,566$ variants. Further, using VCFTOOLS we exclude (i) indels (–remove-indels), (ii) variants with more than 5% missing genotypes (–max-missing), (iii) multiallelic variants (–min-alleles 2 and –max-alleles 2), (iv) variants significantly violating Hardy Weinberg equilibrium (–hwe 0.000001), (v) variants not on the autosomes (vi) variants that are not PASS in the filter column. After filtering, $n = 127,261$ variants remain. The resulting VCF file is converted to plink format and LD pruning is performed with PLINK using parmater ”–indep-pairwise 100 10 0.4”, which results in $n = 97,940$ variants. The resulting LD pruned variants are used for genotype principal component analysis (PCA) and admixture analysis. PCA uses SMARTPCA [21] from the EIGENSOFT software package. Admixture analysis uses ADMIXTURE [22], which is run for $k = 2, ..., 25$ components with parameter ”–cv=10” for assessing cross-validation error to select the best $k$. Visualization of admixture results uses the R package POPHELPER [23]. In order to confirm Egyptian PC location for a different cohort of Egyptians, we used SNP array-based variant data from 398 Egyptians, for details see the next Suppl. Section. Because the number of overlapping variants is too low with the Fernandes and Busby data set, we performed this comparative analysis only for the two large whole genome sequencing-based data sets, Bergström and 1000G. From the Egyptian WGS variant data, we removed rare variants with minor allele frequency less than 5%. Before SMARTPCA computation, we performed LD pruning for both settings using “–indep-pairwise 1000 10 0.2”. This resulted in $n = 171,329$ variants used for the SNP-array setting and $n = 201,707$ variants used for the WGS setting.

### SNP array-based Egyptian variant data

We used PLINK 1.9, PLINK 2.0 and BCFTOOLS 1.8 for the quality control of the Egyptian genotypes. The genotypes were previously used in a genome-wide association study of psoriasis [24] and belong to Egyptian individuals in a case and control cohort. For cases and controls separately, we removed non-autosomal genotypes and genotypes consisting of allele pairs AT or GC. Then we filtered out genotypes with a per variant call rate < 98%, a Hardy-Weinberg-Equilibrium P-value (PHWE) < 0.001 for controls and PHWE < 10-5 for controls and a minor allele count < 5. For the remaining variants, we flipped all genotypes to the forward strand. After that we normalized the variant identifiers and merged the genotypes of the case and the control cohorts, followed by another call rate filtering step in which we removed variants with a call rate < 98% and a samples with a call rate < 95%. Subsequently, we removed individuals with a heterozygosity rate deviating more than 2 standard deviations from the mean. Next, we checked for cryptic relatedness using the KING [25] implementation in PLINK 2.0 and removed one individual for each pair of individuals with a relationship of second degree and closer (> 0.0884). Lastly, we performed a principal component analysis to check for outliers. In an iterative manner (https://gist.github.com/matmu/97f91283e57c68c2aa4cfd83483a17de), we removed individuals that exceeded more than 10 standard deviations from the mean in at least one of the first 5 principal components. This resulted in 206 cases and 398 controls, of which we only used the control individuals for population genetics analyses.

### Mitochondrial haplogroups

Genomic DNA samples were processed for library preparation, as previously described in the Human mtDNA Genome protocol for Illumina Sequencing Platform (http://emea.support.illumina.com/content/dam/illumina-support/documents/documentation/chemistry_documentation/samplepreps_legacy/human-mtdna-genome-guide-15037958-01.pdf). In brief, two primer sets [MTL-F1 (AAA GCA CAT ACC AAG GCC AC) and MTL-R1 (TTG GCT CTC CTT GCA AAG TT); MTL-F2 (TAT CCG CCA TCC CAT ACA TT) and MTL-R2 (AAT GTT GAG CCG TAG ATG CC)] were used to amplify the mtDNA by long-range PCR. Library preparation was performed using a Nextera XT DNA Library Preparation Kit (Illumina Inc., CA, USA), and the 10-pM library was sequenced on the Illumina MiSeq sequencing platform (2x150 bp paired-end reads) (Illumina Inc.). Haplogroup assignment was performed using HaploGrep 2 [26]. In brief, HaploGrep weights each polymorphism present in PhyloTree17 (a phylogenetic tree of worldwide human mitochondrial DNA variation) based on its informativeness to define haplogroups. The set of SNPs in the input file are classified as informative or remaining (not informative). A score is given based on the weights of the "informative SNPs" but it is "penalized" by the number of remaining SNPs. For quality control, our recomputed haplogroup frequencies of 100 Egyptian individuals from Pagani *et al.* [27] have been compared with frequencies reported in the corresponding publication. Recomputed and previously reported frequencies were identical.

### Haplotypic expression

The tool PHASER [28] (version 1.1.1) was used to calculate haplotypic expression for the Egyptian assembly individual using our 10x-phased variants as gold-standard phasing, i.e. without the option to perform phasing from the RNA sequencing data. Suppl. Fig. 43 provides an overview of the analysis. RNA sequencing data obtained from blood was pre-processed with FASTP [29]. STAR version 2.6.1.c [30] was used to align reads to GRCh38 using Ensembl version 95 annotation. RNA-Seq QC was performed with QUALIMAP [31]. In total, expression of 58,738 genes was quantified. In a two-step process, PHASER was first used to calculate haplotypic counts which resulted in 16,566 genes with non-zero counts being analyzed (Suppl. Fig. 44). Of those, we excluded genes with less than 30 reads mapped to both haplotypes. In a second step, for the remaining 7,202 genes, allelic fold change was computed and significance tested using a Binomial test (Suppl. Fig. 45). Both steps were performed as described in the HowTo provided by the PHASER authors (`https://stephanecastel.wordpress.com/2017/02/15/how-to-generate-ase-data-with-phaser`). Multiple testing correction using FDR was performed. Allelic expression results for 1,180 significant genes at FDR of 5% are provided in Suppl. Table 12.

### Data integration with the GWAS catalog

We downloaded associations and ancestry information from the GWAS catalog (https://www.ebi.ac.uk/gwas/, version 2019/08/27), a curated database of published genome-wide association studies. We split associations into 3,064 trait-specific data sets according to the mapped terms from the Experimental Factor Ontology (EFO). For every disease, only associations from studies of individuals from European descent according to column "BROAD ANCESTRAL CATEGORY" were kept. In a next screen we kept only associations of variants for which there is another variant within 1 MB that is associated with the same trait. In order to select a set of common, high-quality European GWAS SNPs we used the genome-wide genotypic data of 503 individuals of European descent from the state-of-the-art 1000 Genomes data and keep bi-allelic variants with more than 5% minor allele frequency, not violating Hardy-Weinberg Equilibrium (p=0.000001), from which we selected variants with rsIDs matching associations in the GWAS catalog. For every trait we computed all-versus-all linkage disequilibrium of the remaining associated SNPs and output all pairs of SNPs within 1 MB with LD of $R^2 \geq 0.8$. Associations of SNPs with at least one other associated SNP in LD were kept and can be considered a replicated association signal. At this step we thus exclude associations reported in multiple studies, but always for the exactly same SNP and not for at least one other SNP in LD. For 585 trait ontology terms we kept a minimum of 2 associations. In a next step, we kept one associated SNP per trait and locus, where we define a locus as a region of 1 MB with a tag SNP. We automatically selected the tag SNP based on the number of studies reporting the rsID position for the respective trait. Note that we may have exclude additional, independent association signals that are closer than 1 MB from a SNP selected as tag SNP. The resulting tag SNPs were combined over all traits; there are 4,008 such SNPs. Of these, 261 (6.5%) have not been called in the Egyptian cohort. We investigated these closer and 42 are located within the MHC locus (chr6:28510020-33480577) and very few have other variants (largely indels) with start position +/-15 BP, possibly compromising the variant calling. For the tag SNPs we computed proxy SNPs that are in LD ($R^2 \geq 0.8$) within 1 MB using genotypes of the European and Egyptian cohort, respectively. In visualizations we display data for 3,959 tag SNPs with missing genotypes in at most 10% of Egyptian individuals.

### Sequencing read mapping to GRCh38

We performed quality control on raw FASTQ files using FASTQC (https://www.bioinformatics.babraham.ac.uk/projects/fastqc). All NGS data was mapped to GRCh38. For PacBio long reads, we used MINIMAP2 [2] with option "-x map-pb" for mapping PacBio genomic reads. For Illumina short read data, we used BWA-MEM [3] with default options. QC for mapped read data was performed using SAMTOOLS STATS [32]. Linked 10x genomics reads were processed by LONGRANGER BASIC (version 2.2.2) to generate barcoded FASTQ files, which were subsequently processed by LONGRANGER WGS, which outputs also a BAM mapping file of linked reads.

### Alignment to GRCh38

We aligned all assemblies with GRCh38 using NUCMER from the recently updated MUMMER4 suite [33] with default options, except additionally using "—mum" which denotes that alignment anchor matches need to be unique in both GRCh38 and assembly. We filtered alignments using MUMMER4's delta-filter with option "-q", which maps each assembly position onto the best hit in GRCh38, allowing for GRCh38 overlaps, as well as option "-1", which performs 1-to-1 alignment allowing for rearrangements. Alignments from 1-to-1 mapping have been further used for assembly-based variant detection using NUCDIFF [34]. MUMMER4's mummerplot script has been used for dotplot visualization (Suppl. Figs. 6-10).

### Assembly-based variant identification

We identified SNVs, indels and structural variants present in the assembly by aligning it with the human reference genome GRCh38 and detecting sequence differences. This was achieved by using the tool NUCDIFF [34] on selected genomic regions utilizing genome-wide EGYPT-to-GRCh38 alignments from 1-to-1 mapping (see last section).

### Gene-centric integrative data views

Most users of the Egyptian genome reference will be interested in a specific genetic region or a specific gene and would like to investigate, if and where this region or gene is affected by personal or population-specific variation. To facilitate such analysis, we implemented a workflow to extract all relevant Egyptian data within a specified region centered around a gene of interest (by default +/- 100 kB). The resulting files can be viewed in the Integrative Genomics Viewer (IGV) [35]. The gene-centric data contains

- EGYPT PacBio long reads mapped to GRCh38 (BAM)
- EGYPT Illumina short reads mapped to GRCh38 (BAM)

- EGYPT 10x linked reads mapped to GRCh38 (BAM)
- EGYPT blood RNA-Seq reads mapped to mapped to Ensembl genes version 95 given with GRCh38 coordinates (BAM)
- EGYPT assembly differences to GRCh38 (BED)
- AK1 assembly differences to GRCh38 (BED)
- YORUBA assembly differences to GRCh38 (BED)
- All, common and population-specific small variants of 110 Egyptian individuals including EGYPT (VCF)
- VEP annotations of small variants (flat file)
- Small variants from all 1000 Genomes phase 3 individuals with genotypes (VCF)
- Ensembl gene annotation version 95 (GTF)
- Variant data from dbSNP (VCF)

### Supplementary Figures

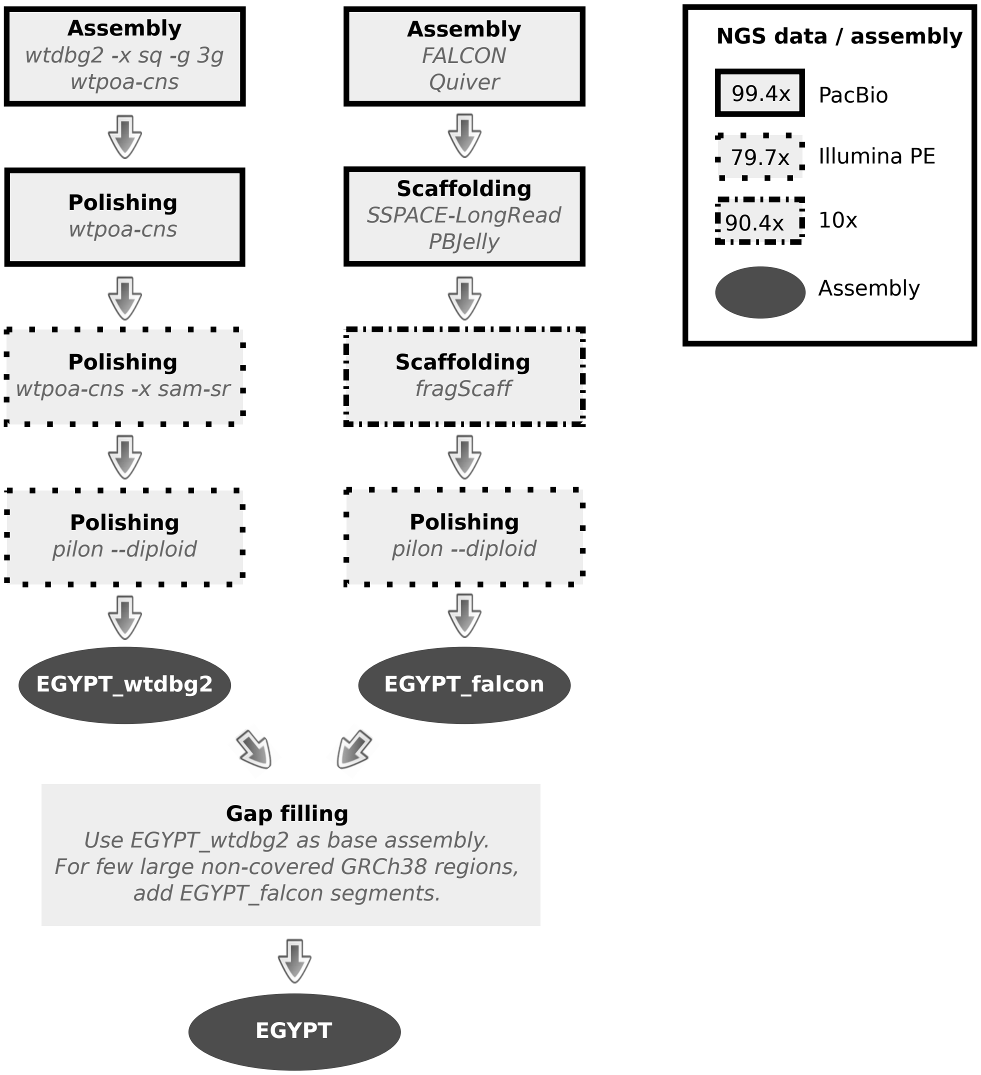

Supplementary Figure 1: Overview of the *de novo* assembly strategy with the NGS data types used in every step. EGYPT_wtdbg2 and EGYPT_falcon are two individual, alternative high quality *de novo* assemblies, which we combined into a final meta assembly that we refer to as EGYPT.

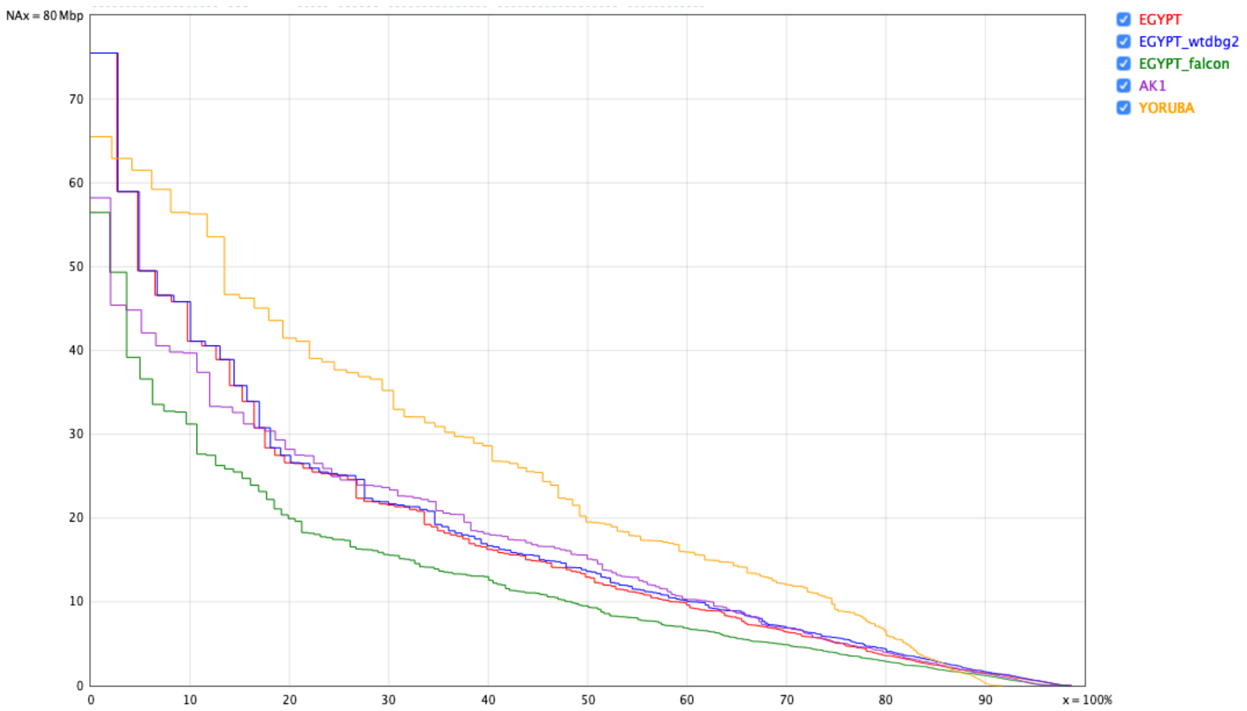

Supplementary Figure 2: NAx display of the N-values of the five assemblies; NA-values are like N-values, but contigs are split at misassembly sites.

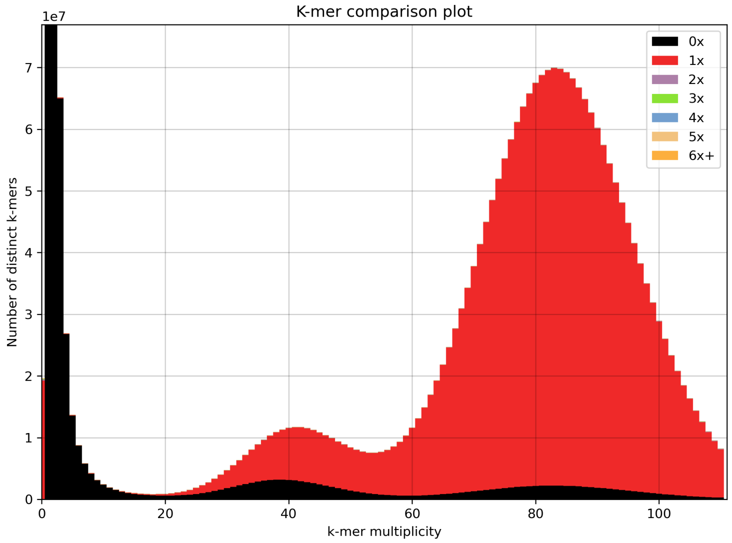

Supplementary Figure 3: K-mer multiplicity with KAT [36] for the `WTDBG2`-based assembly EGYPT_wtdbg2. The second, high peak is at about 90, which is the sequencing coverage. The first, smaller peak is caused by heterozygous variants, k-mers including such variant positions have a coverage of approximately 45x.

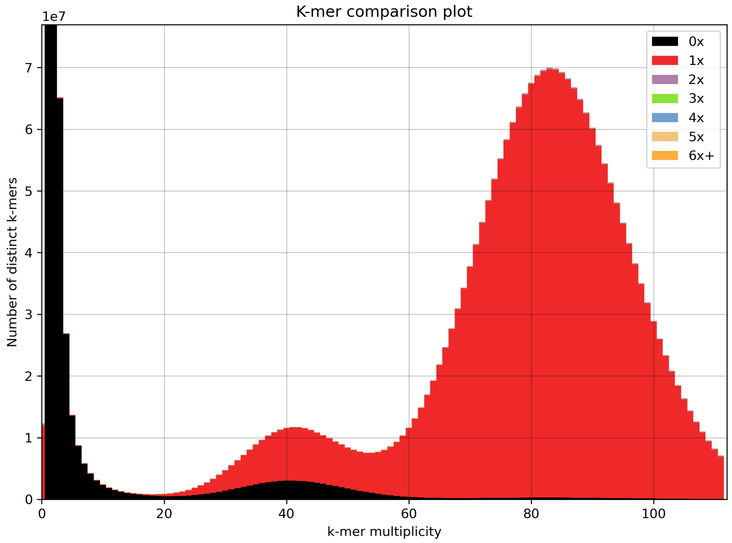

Supplementary Figure 4: K-mer multiplicity with KAT [36] for the `FALCON`-based assembly EGYPT_falcon. The second, high peak is at about 90, which is the sequencing coverage. The first, smaller peak is caused by heterozygous variants, k-mers including such variant positions have a coverage of approximately 45x.

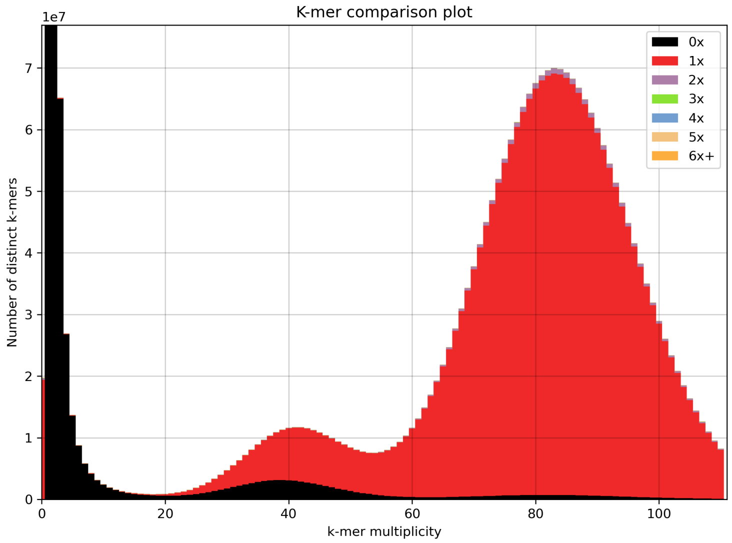

Supplementary Figure 5: K-mer multiplicity with KAT [36] for the final meta assembly EGYPT. The second, high peak is at about 90, which is the sequencing coverage. The first, smaller peak is caused by heterozygous variants, k-mers including such variant positions have a coverage of approximately 45x. There is very slightly more k-mers occuring twice in the meta assembly compared to the draft assemblies.

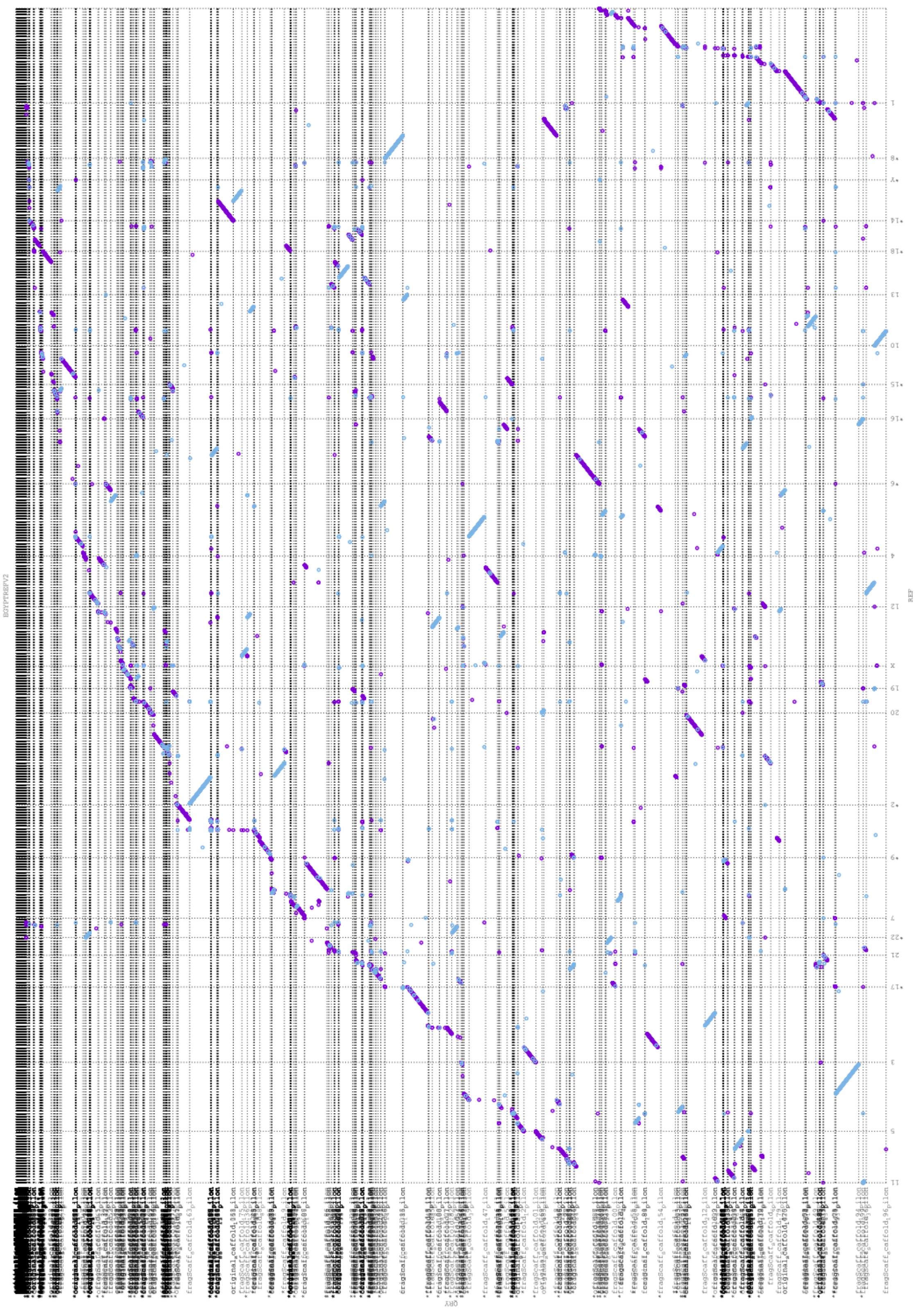

Supplementary Figure 6: EGYPT_falcon assembly dotplot for 1-to-1 alignment with GRCh38.

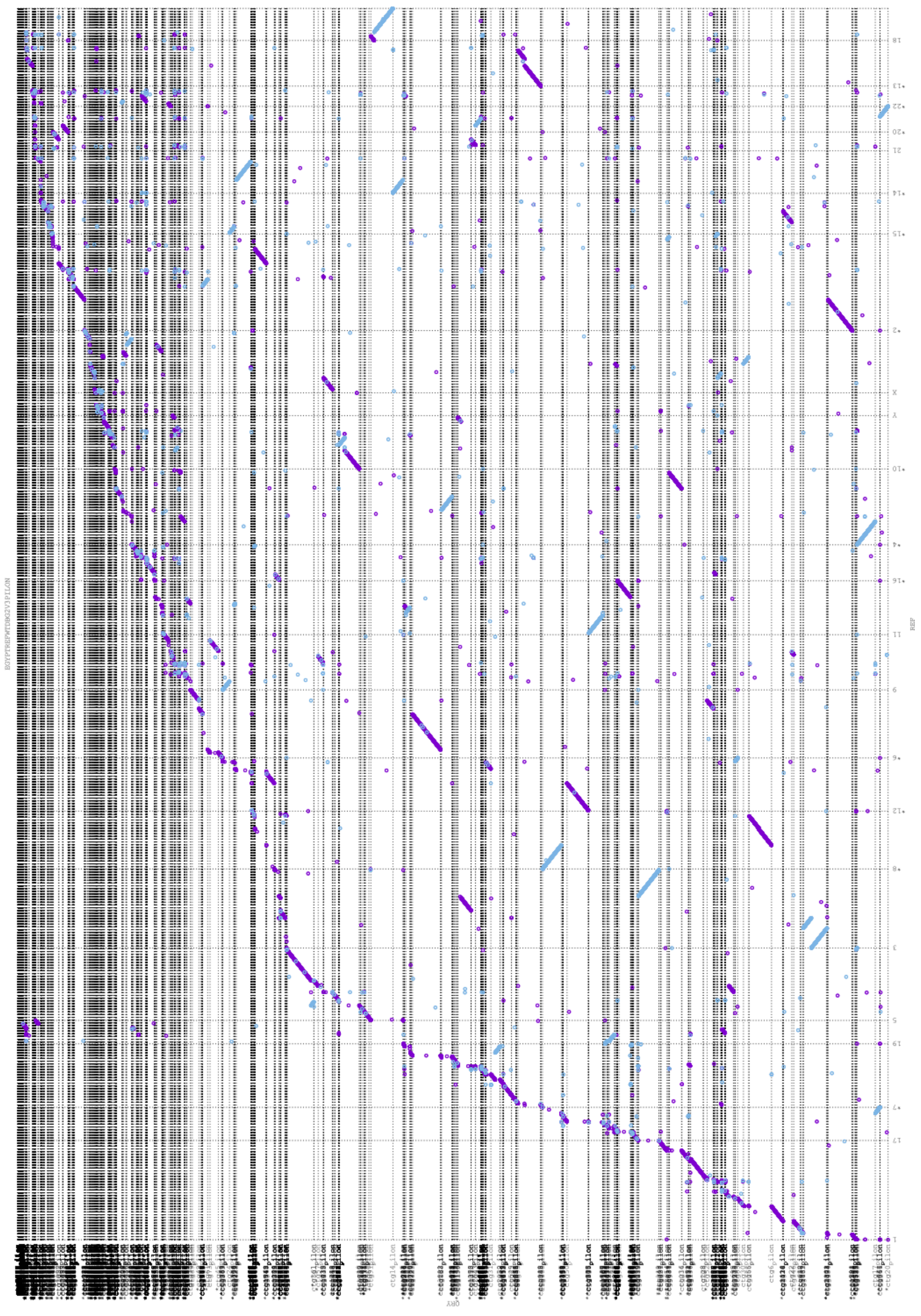

Supplementary Figure 7: EGYPT_wtdbg2 assembly dotplot for 1-to-1 alignment with GRCh38.

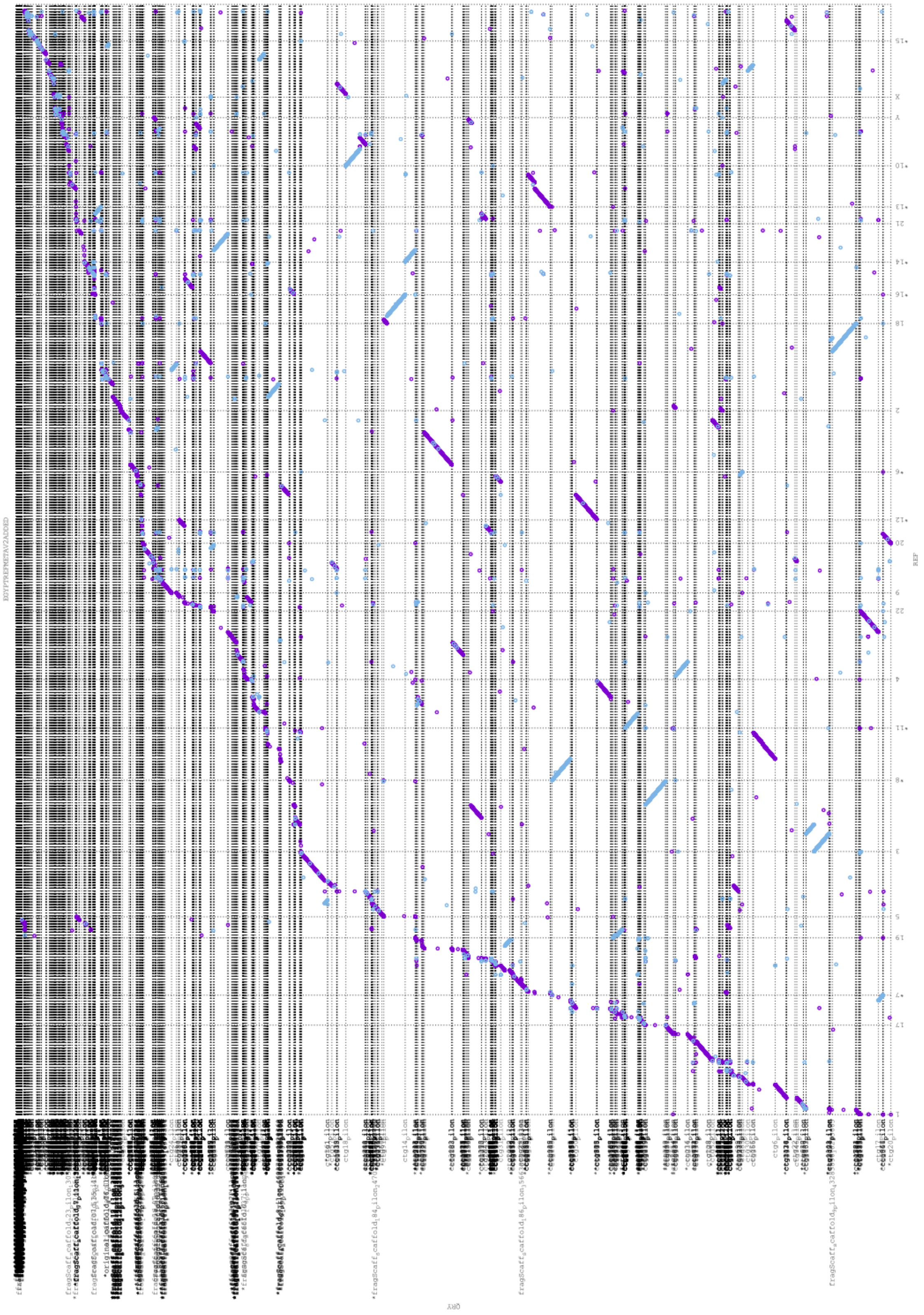

Supplementary Figure 8: EGYPT assembly dotplot for 1-to-1 alignment with GRCh38.

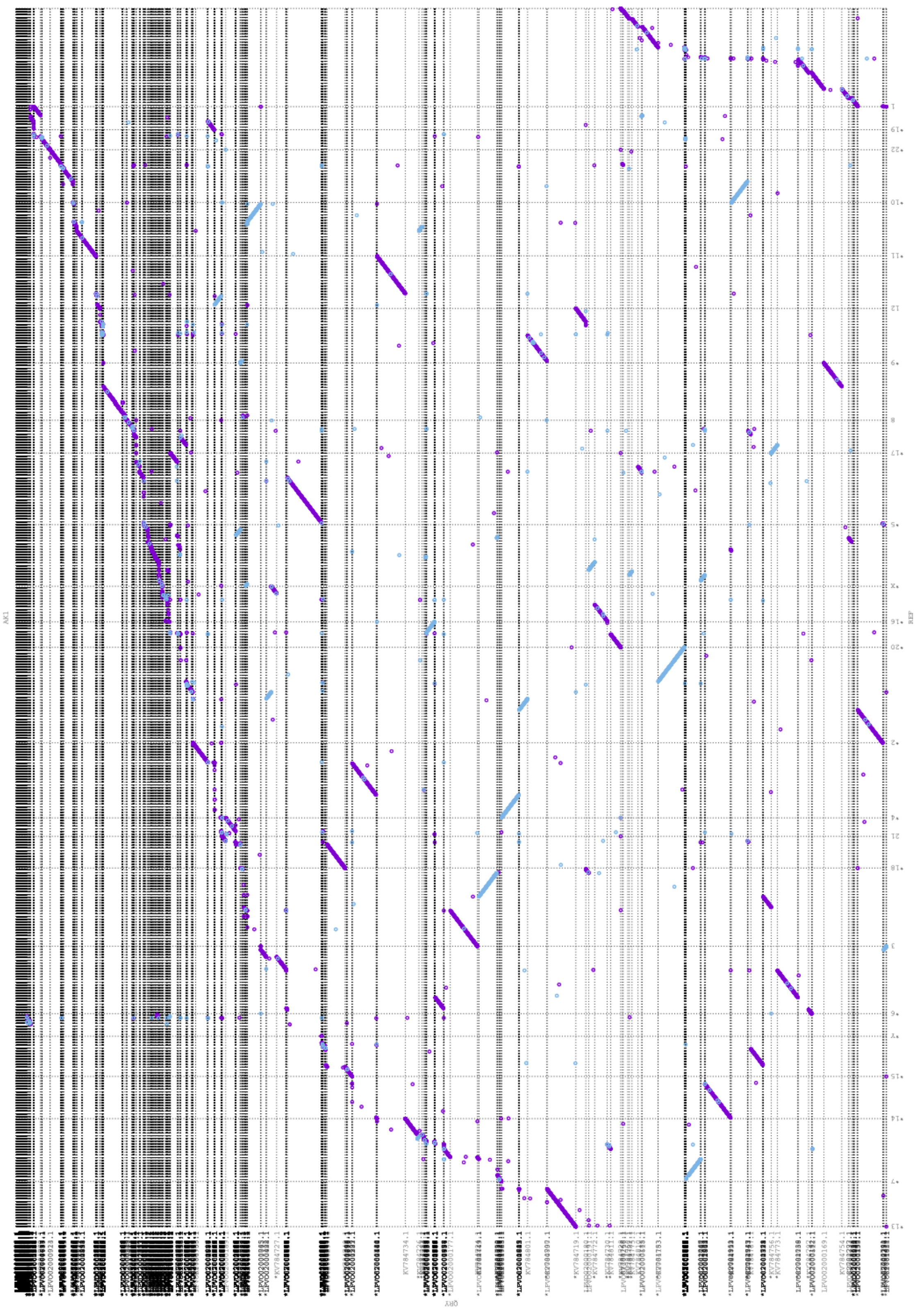

Supplementary Figure 9: AK1 assembly dotplot for 1-to-1 alignment with GRCh38.

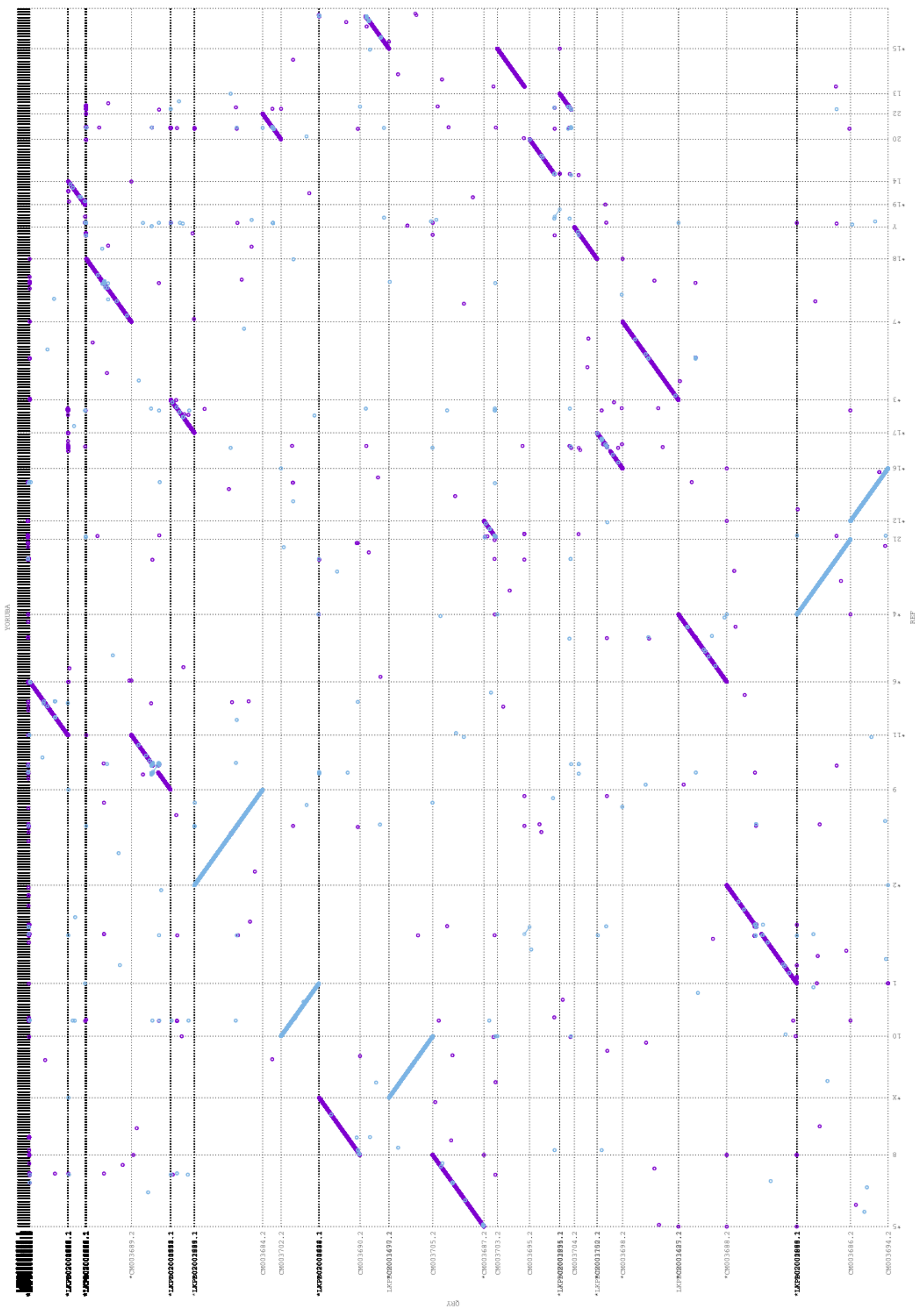

Supplementary Figure 10: YORUBA assembly dotplot for 1-to-1 alignment with GRCh38.

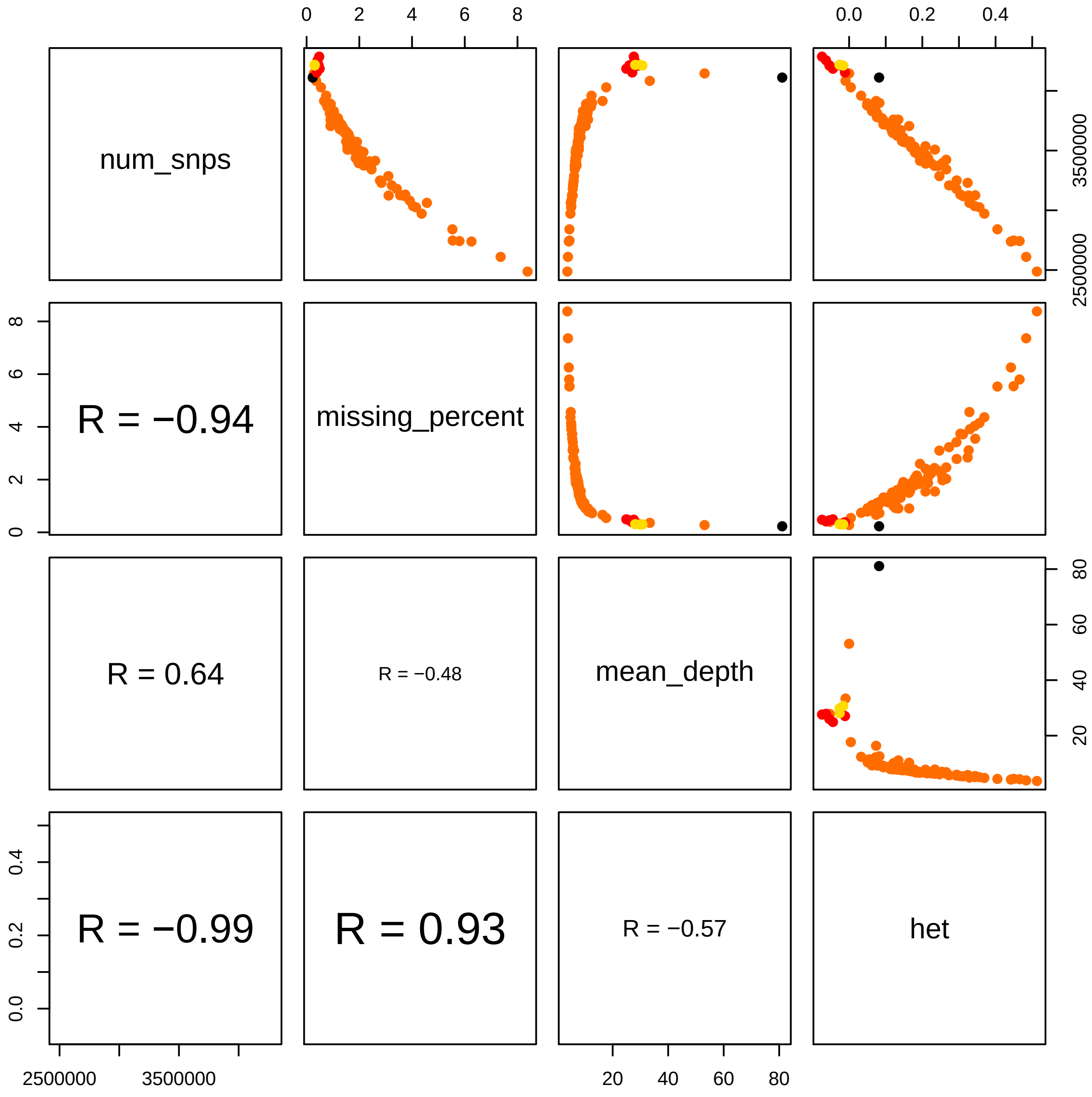

Supplementary Figure 11: Correlations between variant statistics. Pagani *et al.* individuals are orange, Egyptians from Delta red, Upper Egyptians yellow and the EGYPT individual black.

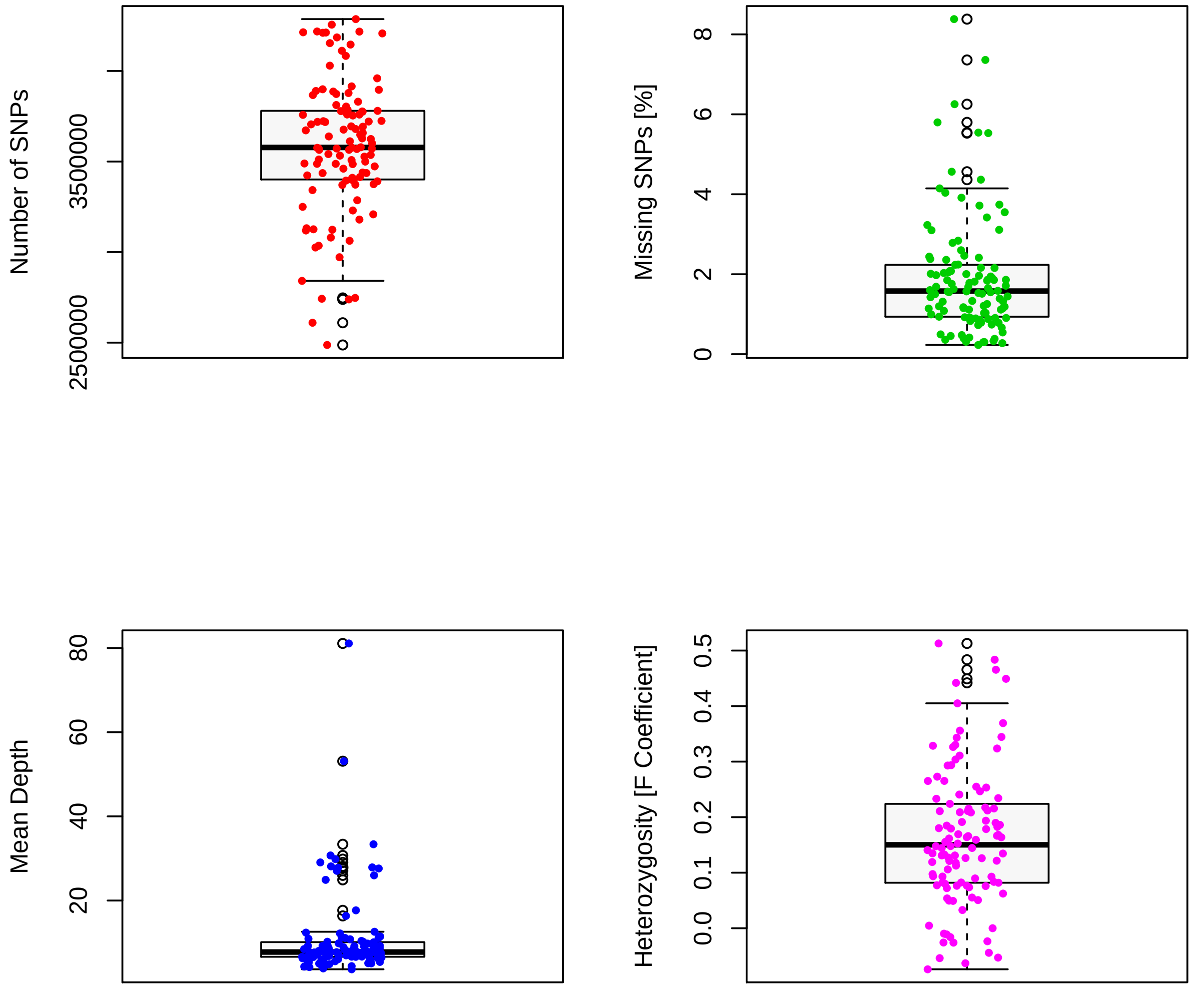

Supplementary Figure 12: Boxplots of number of SNV, percentage of missing SNV calls, mean depth and heterozygosity, each for the cohort of 110 Egyptians.

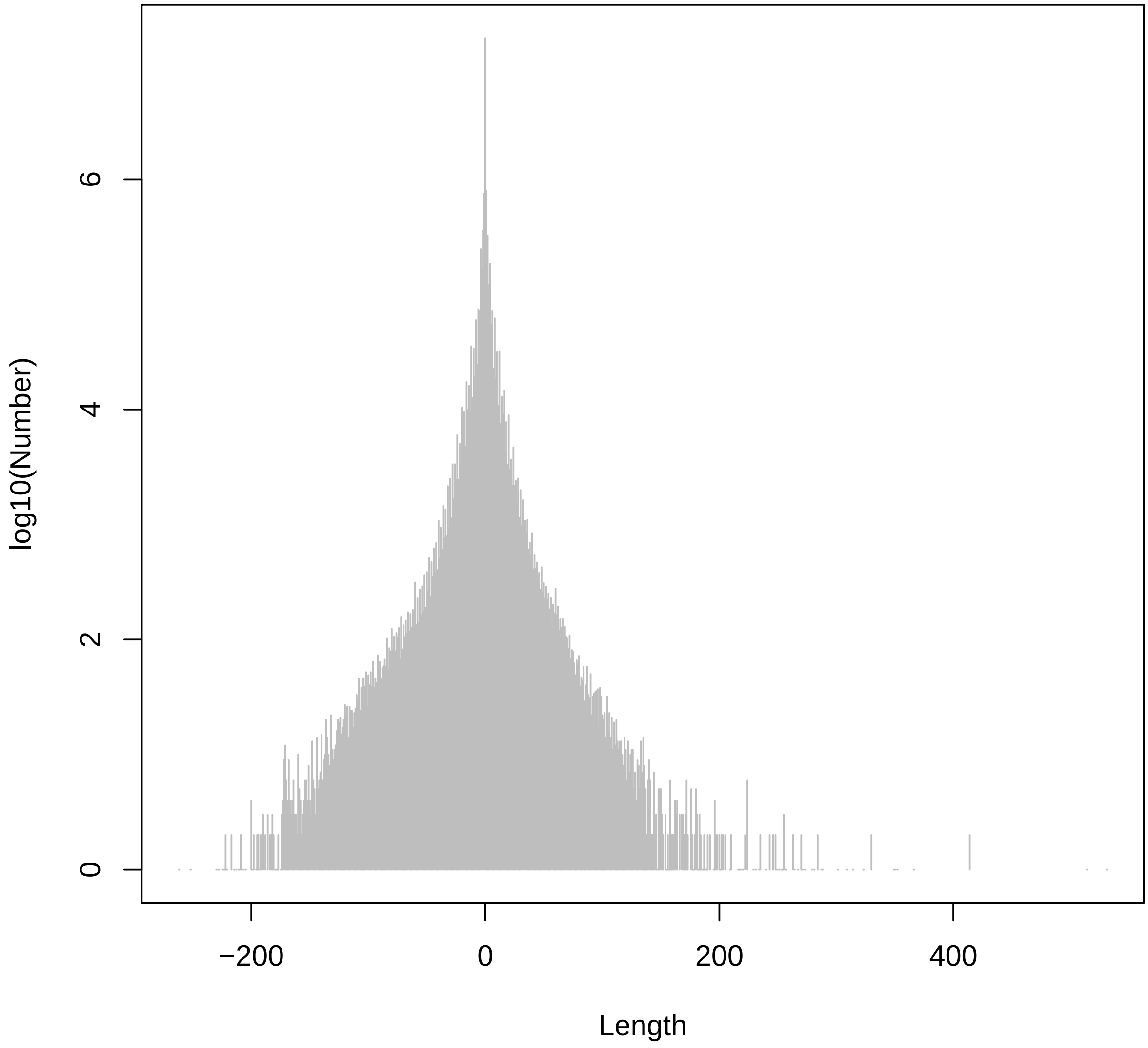

Supplementary Figure 13: Histogram of indel sizes: negative length refers to deletions, positive length to insertions

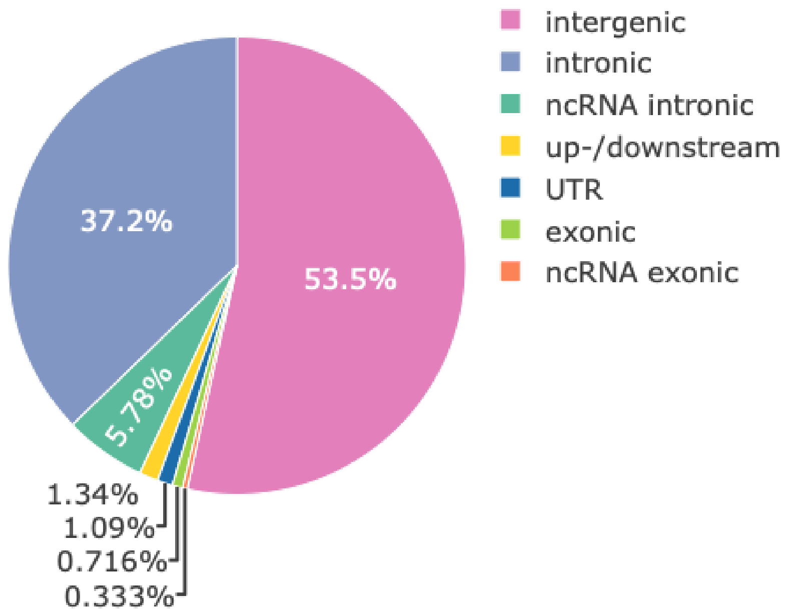

Supplementary Figure 14: Classification of small variants according to genomic location according to `Annovar` annotations.

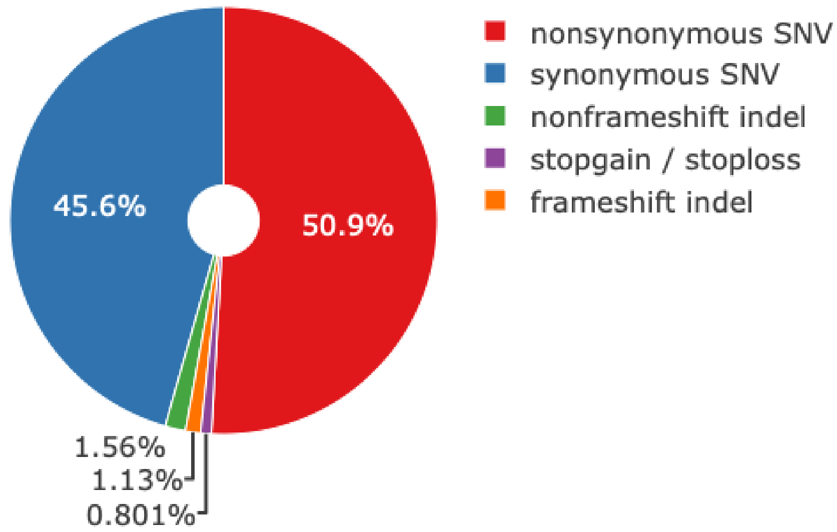

Supplementary Figure 15: Classification of small variants according to exonic variant consequences.

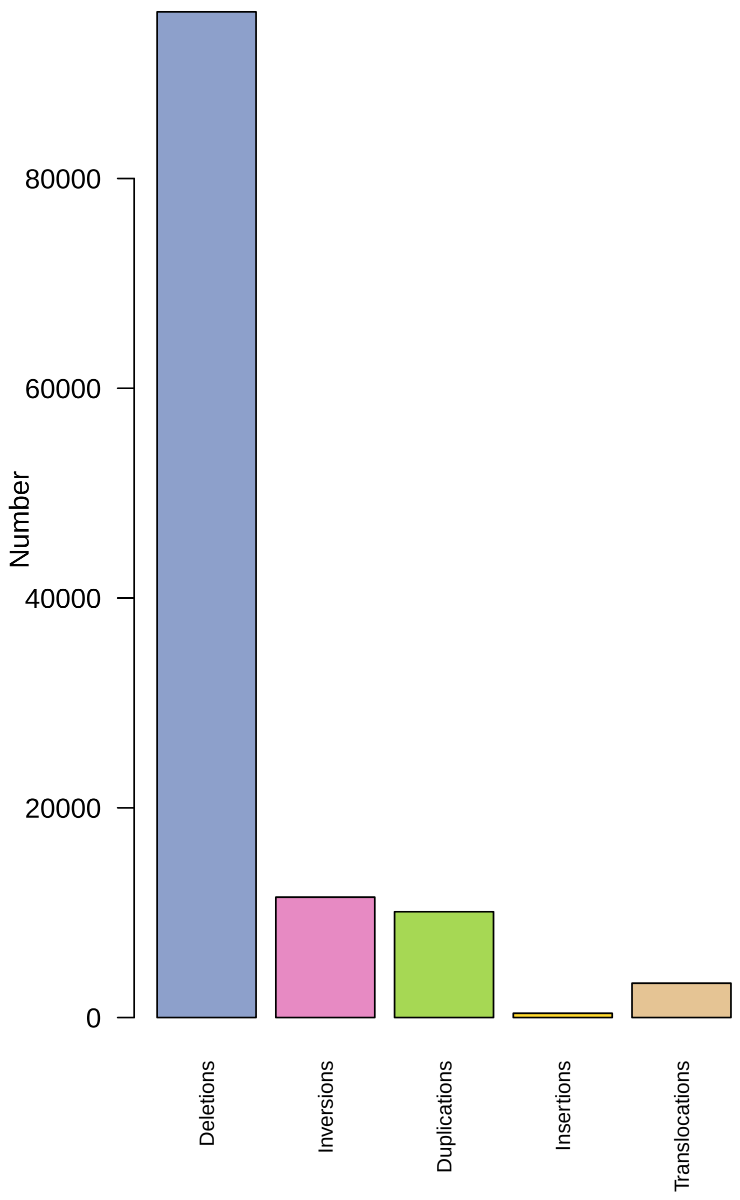

Supplementary Figure 16: Histogram of types of SV calls.

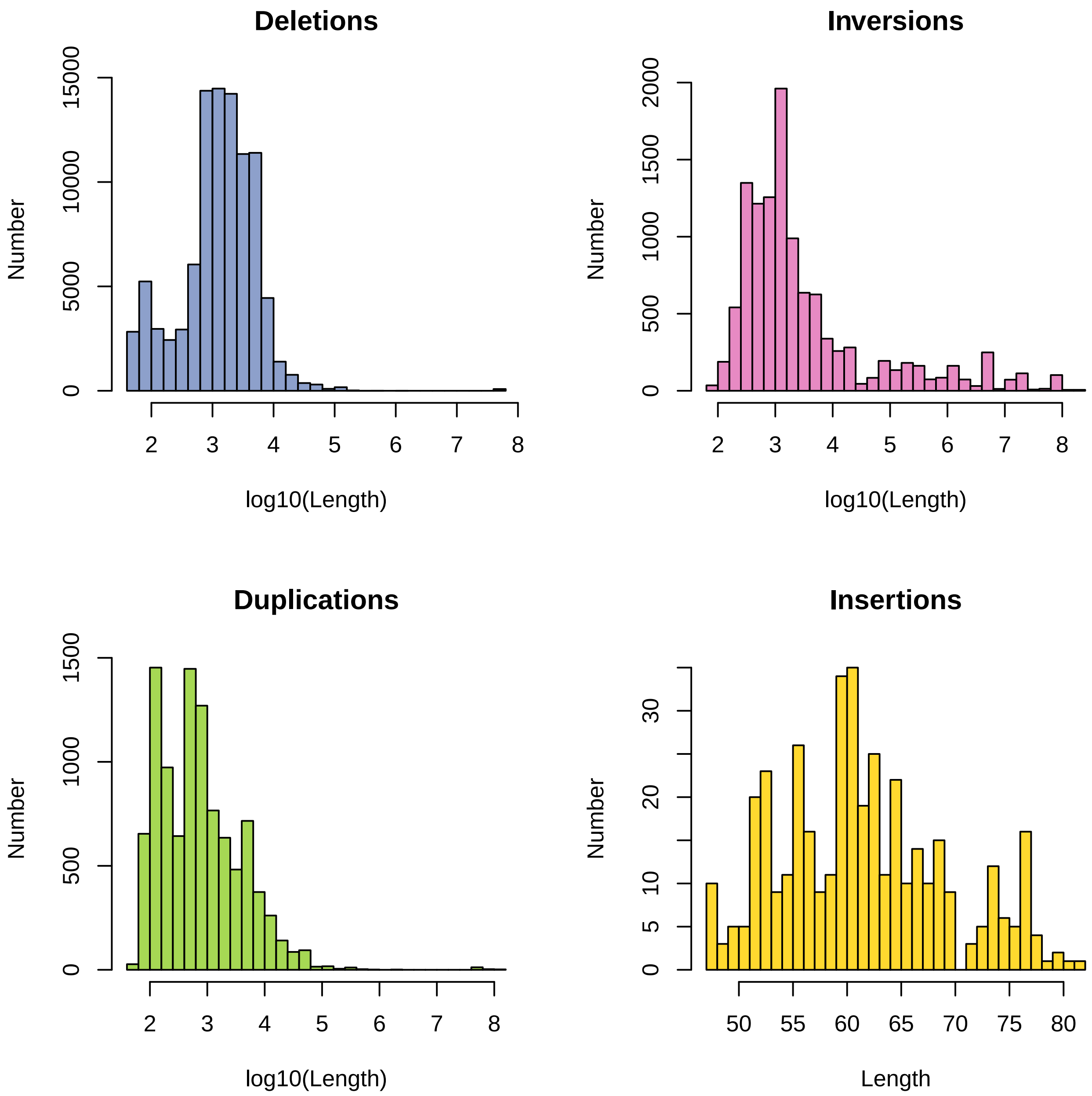

Supplementary Figure 17: Histogram of lengths of deletions, inversions, duplications and insertions passing `DELLY` filter.

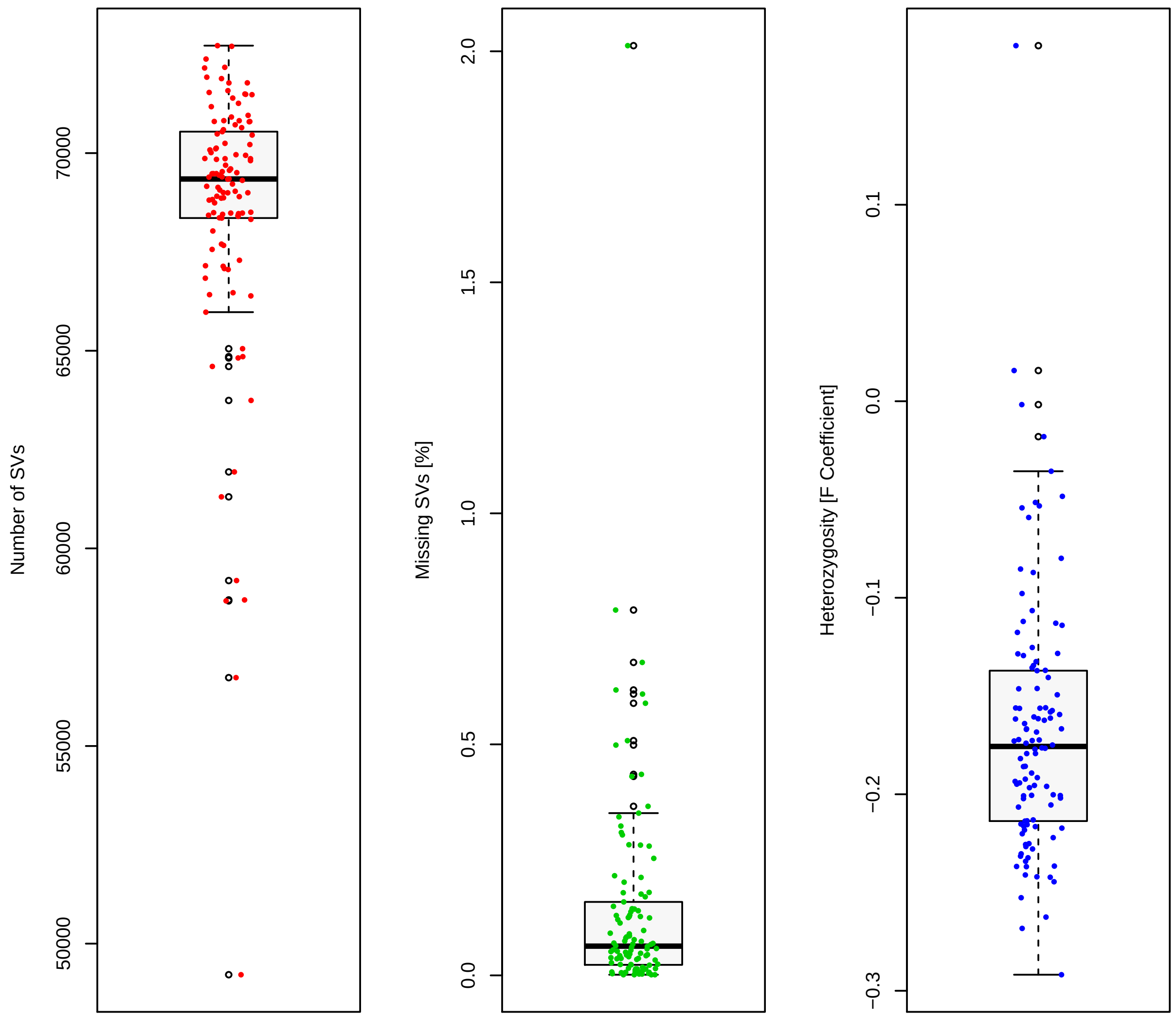

Supplementary Figure 18: Boxplots of number of SV calls, missing SV calls and SV-based heterozygosity for 110 Egyptian individuals.

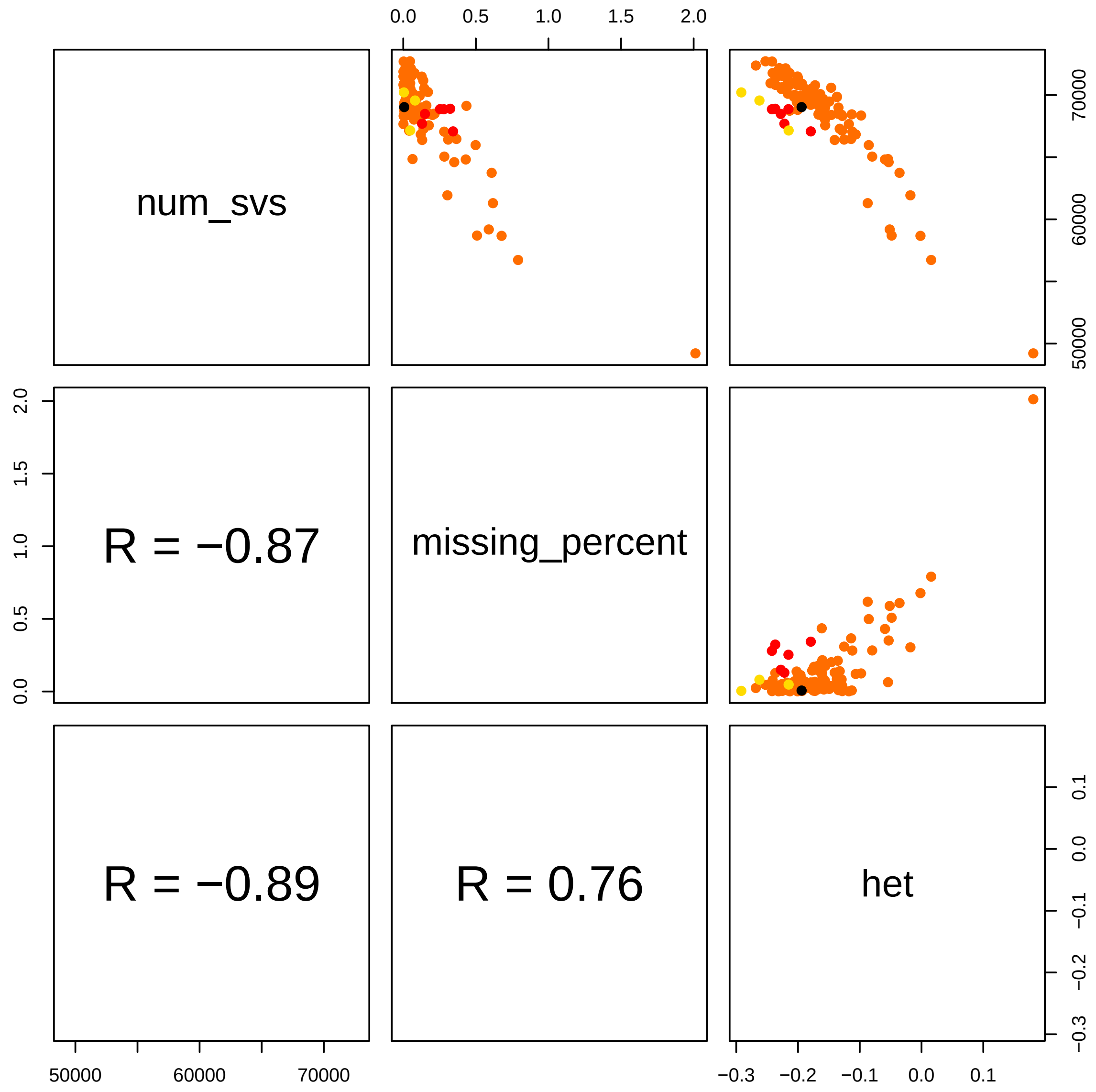

Supplementary Figure 19: Scatterplots and correlation of SV call numbers, missing SV calls and heterozygosity. Pagani *et al.* individuals are orange, Egyptians from Delta red, Upper Egyptians yellow and the EGYPT individual black.

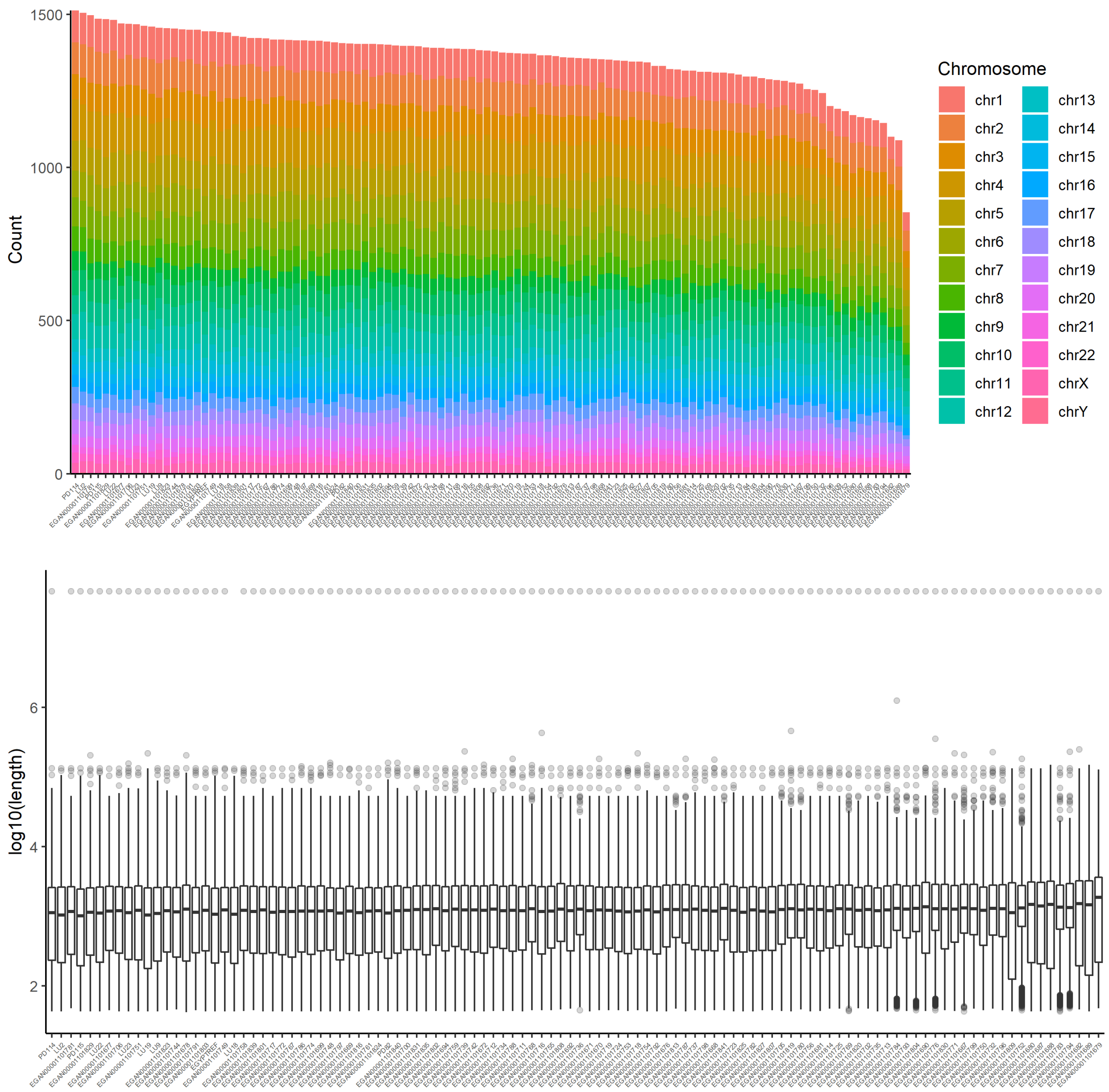

Supplementary Figure 20: Number of deletions and boxplots of deletion lengths per individual after collapsing SVs.

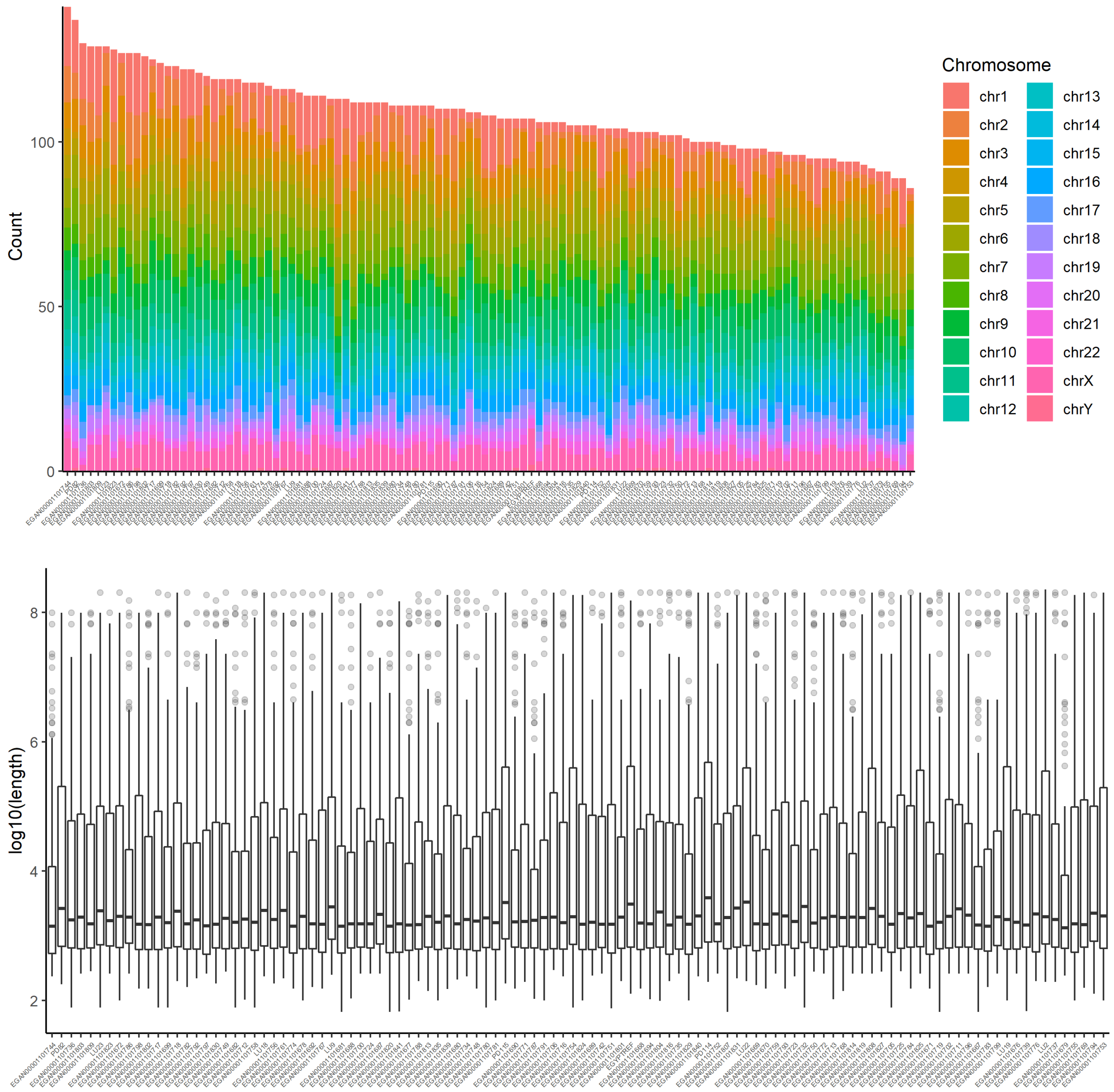

Supplementary Figure 21: Number of inversions and boxplots of inversion lengths per individual after collapsing SVs.

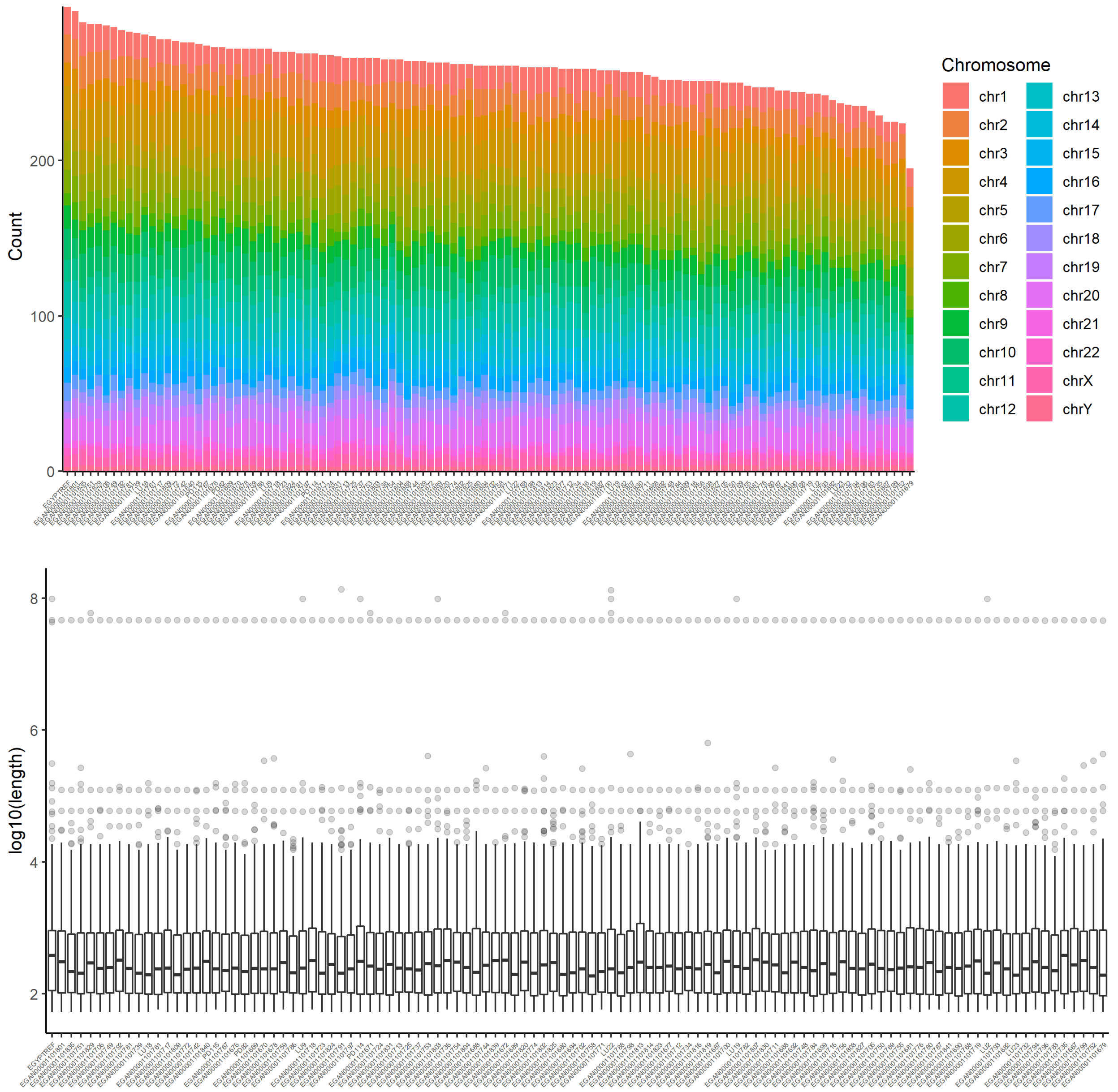

Supplementary Figure 22: Number of duplications and boxplots of duplication lengths per individual after collapsing SVs.

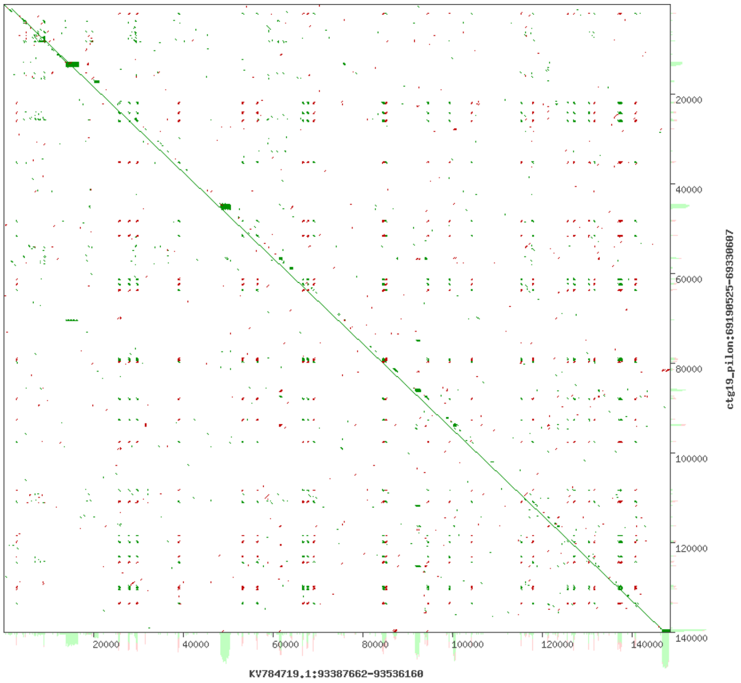

Supplementary Figure 23: Dot plot for GRCh38 gap covering sequence at chr13:111,703,856-111,843,441 for AK1 versus EGYPT assembly sequence; generated with `YASS` [37]

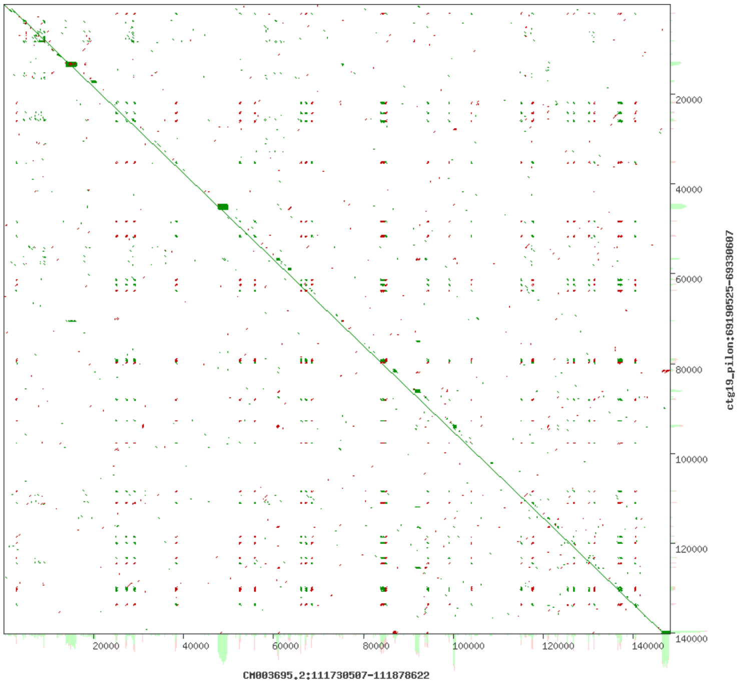

Supplementary Figure 24: Dot plot for GRCh38 gap covering sequence at chr13:111,703,856-111,843,441 for YORUBA versus EGYPT assembly sequence; generated with `YASS` [37]

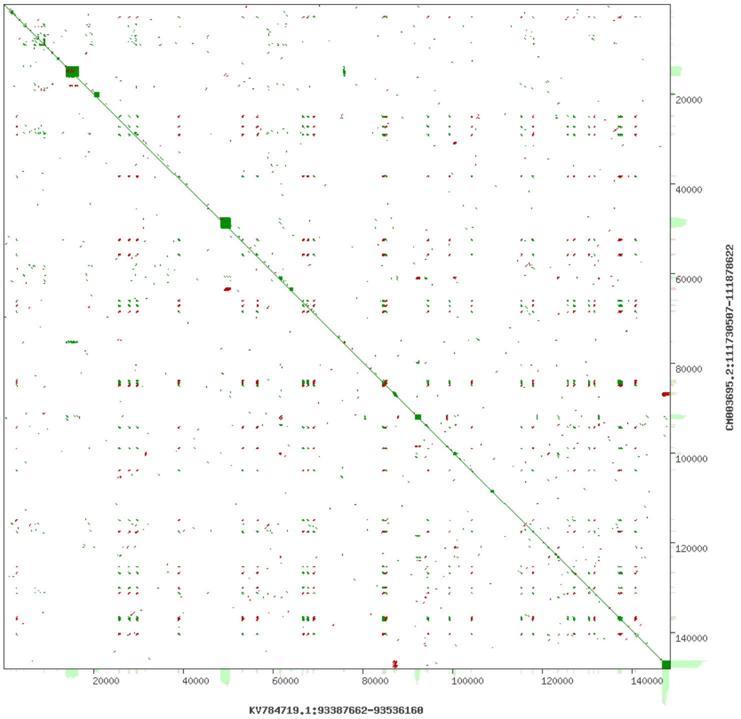

Supplementary Figure 25: Dot plot for GRCh38 gap covering sequence at chr13:111,703,856-111,843,441 for AK1 versus YORUBA assembly sequence; generated with `YASS` [37]

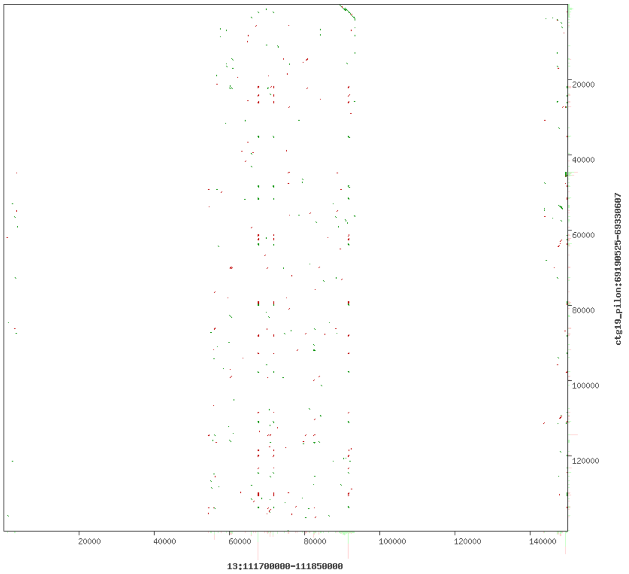

Supplementary Figure 26: Dot plot for GRCh38 gaps chr13:111,703,856-111,753,855 and chr13:111,793,442-111,843,441 and GRCh38 sequence in between versus EGYPT sequence; generated with `YASS` [37]

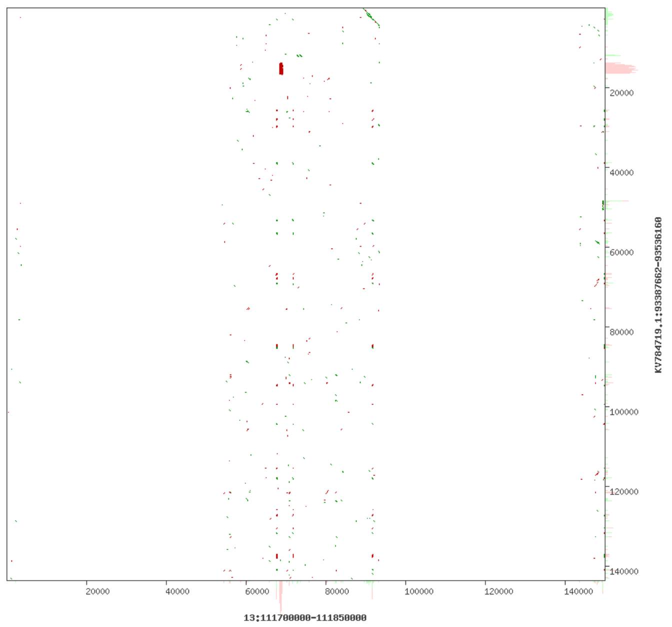

Supplementary Figure 27: Dot plot for GRCh38 gaps chr13:111,703,856-111,753,855 and chr13:111,793,442-111,843,441 and GRCh38 sequence in between versus EGYPT sequence; generated with `YASS` [37]

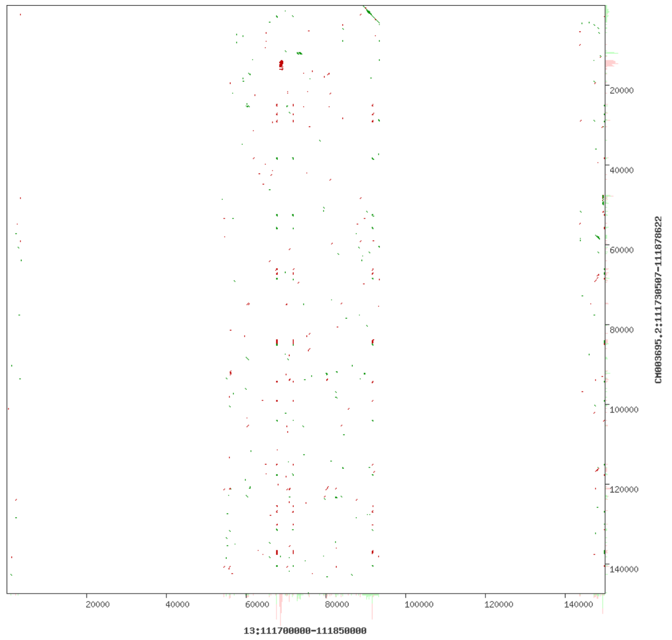

Supplementary Figure 28: Dot plot for GRCh38 gaps chr13:111,703,856-111,753,855 and chr13:111,793,442-111,843,441 and GRCh38 sequence in between versus YORUBA sequence; generated with `YASS` [37]

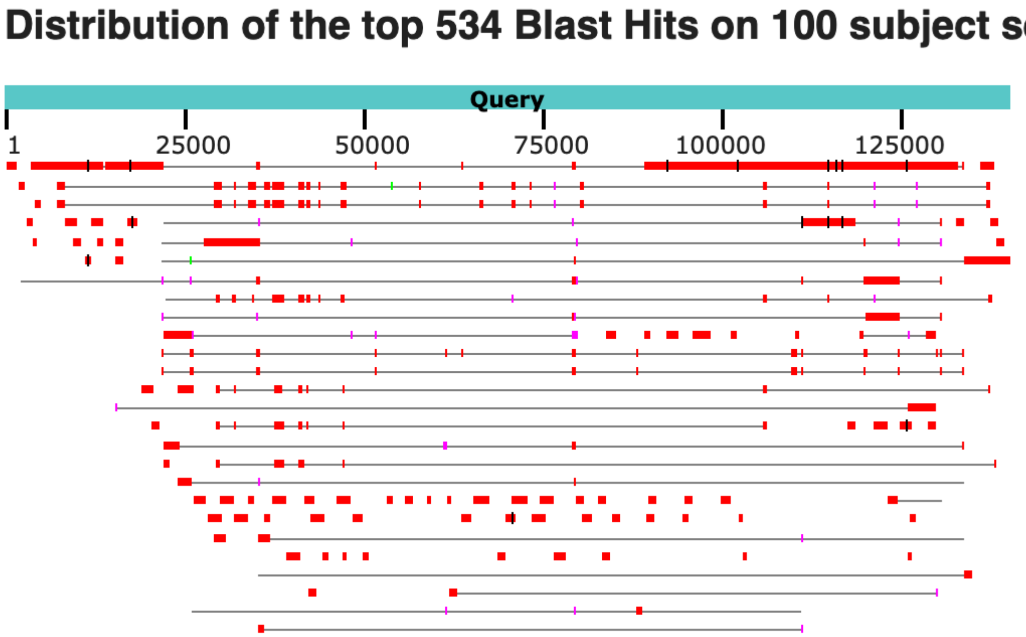

Supplementary Figure 29: A graphic BLAST result displays database sequences similar to the 140 kb gap covering assembly sequence. Alignment with "Homo sapiens chromosome 13 clone WI2-2182D8" (AC188786.1) covers 32% of the assembly sequence (top row right), alignment with "Homo sapiens clone 059F breakpoint junction genomic sequence" (KY503279.1) 12% (top row left).

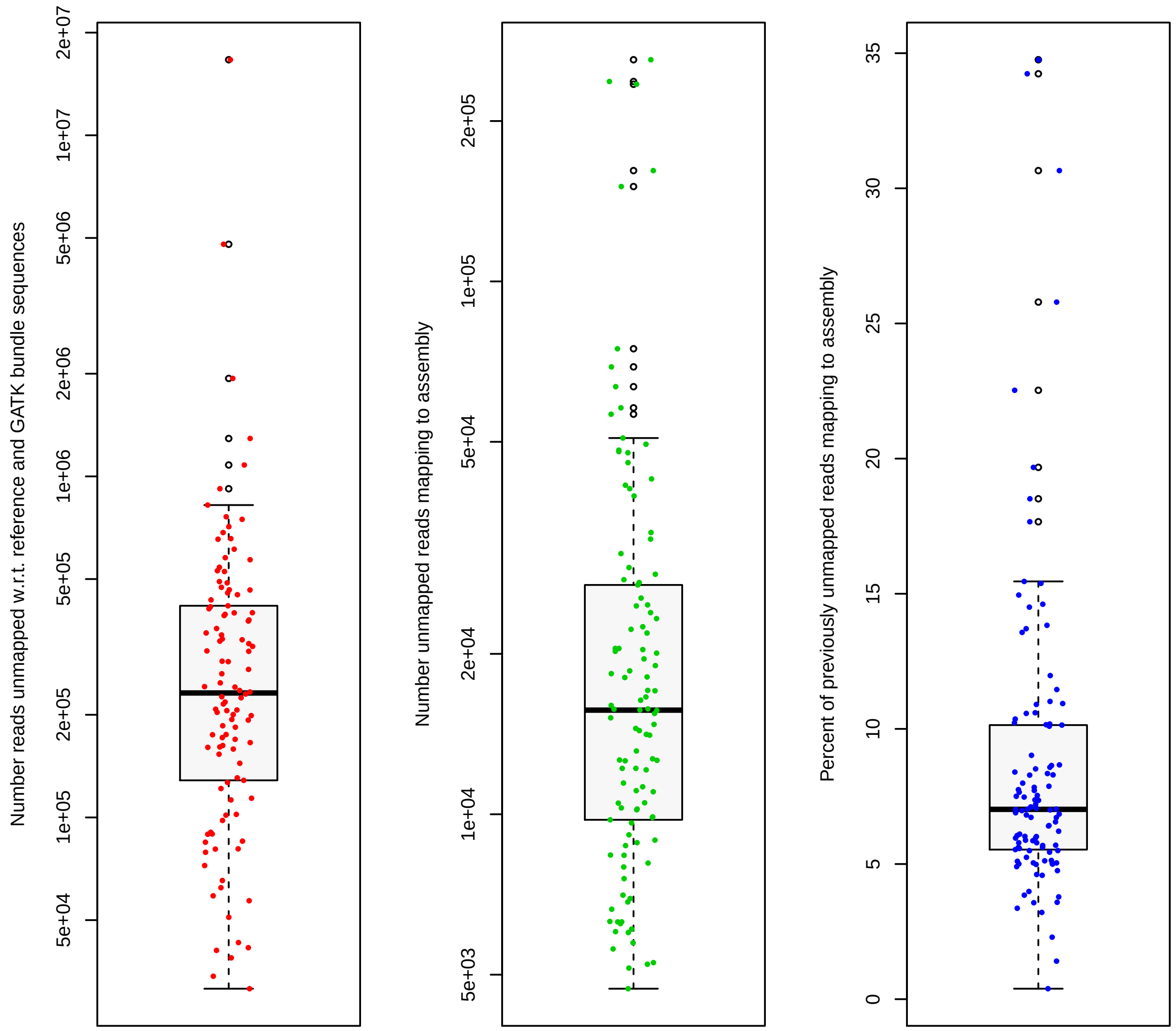

Supplementary Figure 30: Boxplots for short read data of 110 Egyptian individuals. Left: Number of reads not mapped to the reference genome or other GATK bundle sequences. Center: Number of unmapped reads that map to the EGYPT assembly. Right: Percentages of previously unmapped reads mapping to the EGYPT assembly.

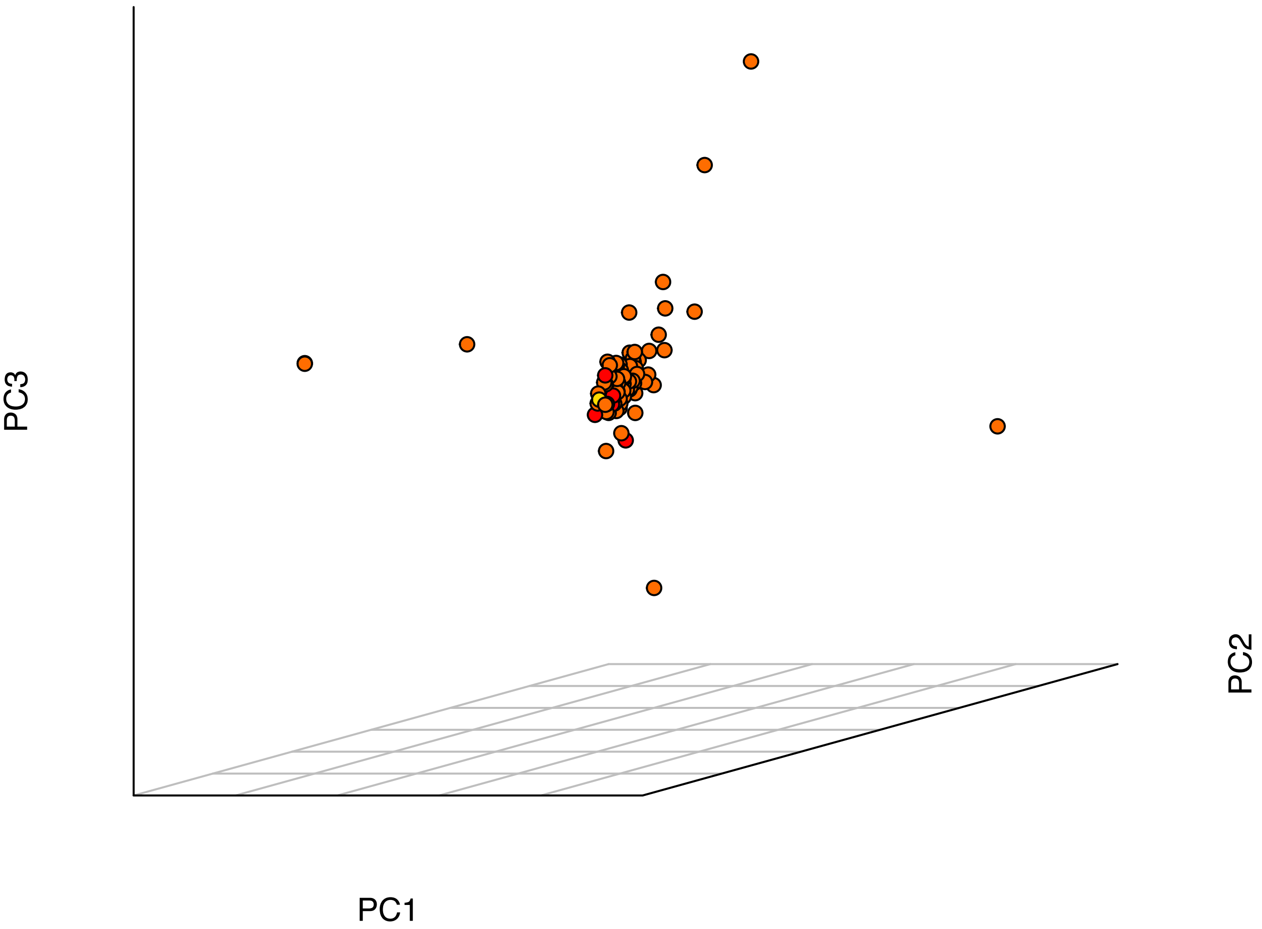

Supplementary Figure 31: Genotype principal components 1, 2 and 3 from Egyptian-only PCA. Red: Egyptian - Nile Delta; Yellow: Egyptian - Upper Egypt; Orange: Egyptian from Pagani *et al.*

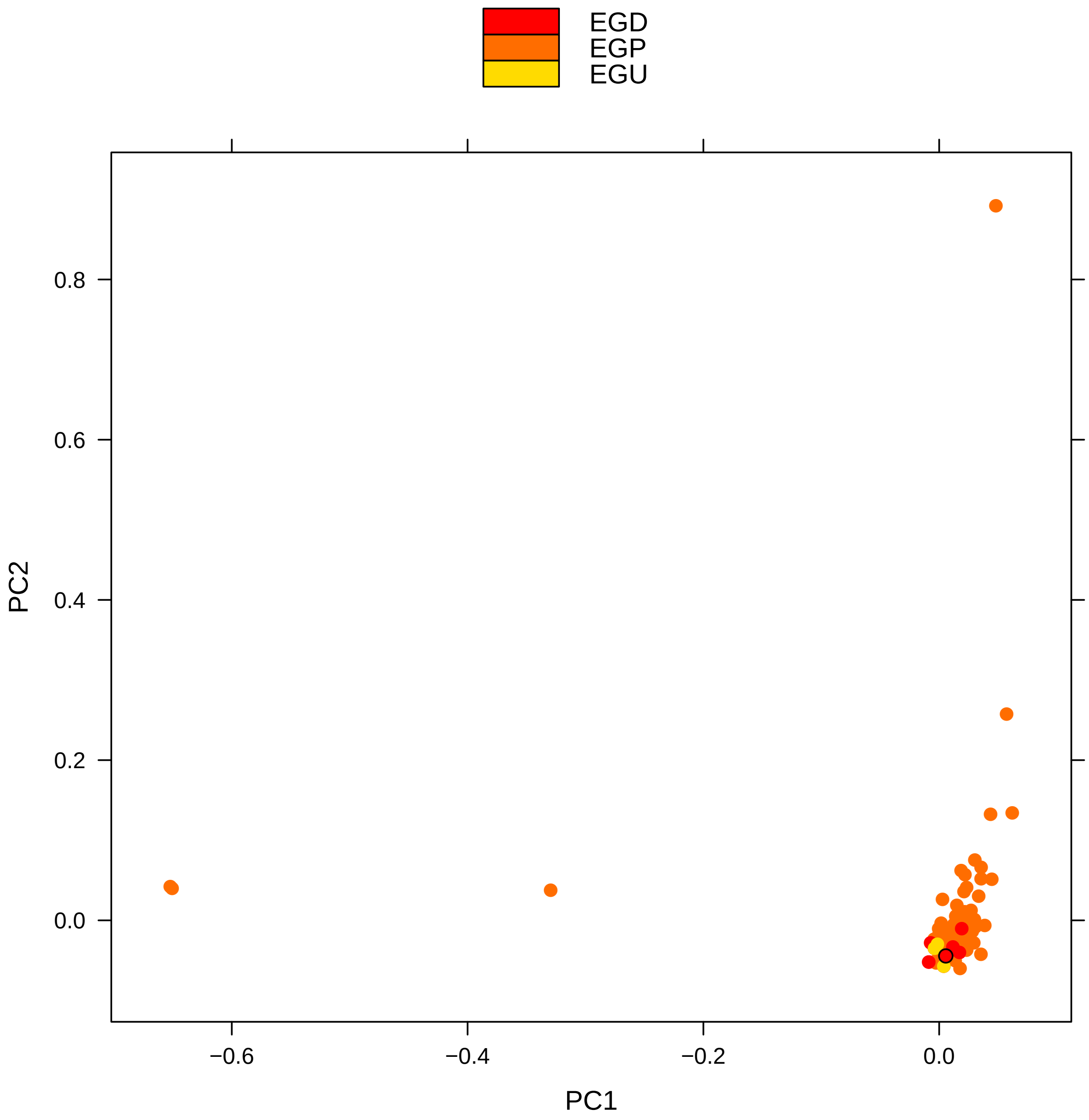

Supplementary Figure 32: Genotype principal components 1 versus 2 from Egyptian-only PCA. EGD: Egyptian - Nile Delta; EGU: Egyptian - Upper Egypt; EGP: Egyptian from Pagani *et al.* A black circle denotes the assembly individual EGYPT.

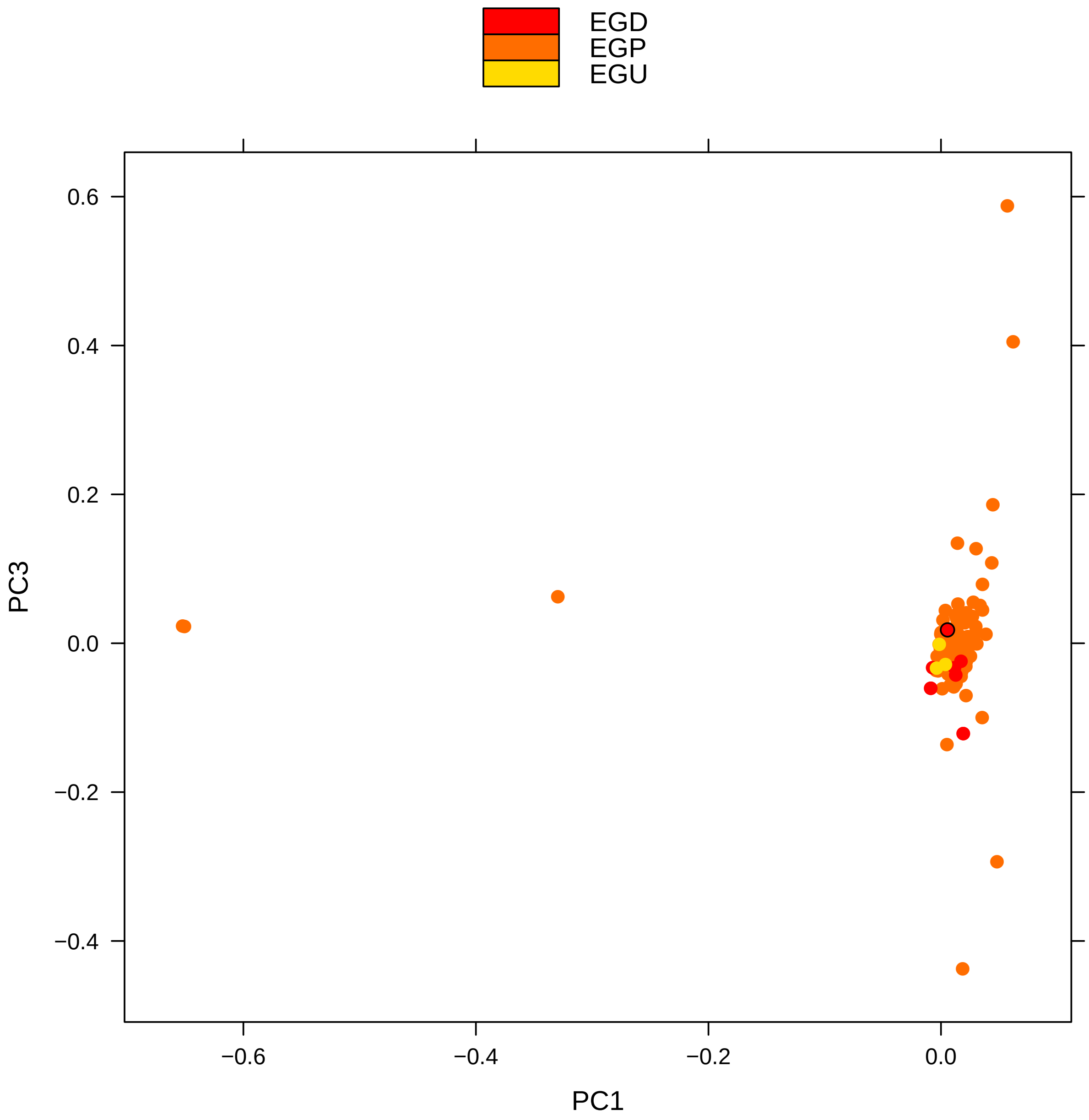

Supplementary Figure 33: Genotype principal components 1 versus 3 from Egyptian-only PCA. For label description see Suppl. Fig. 32

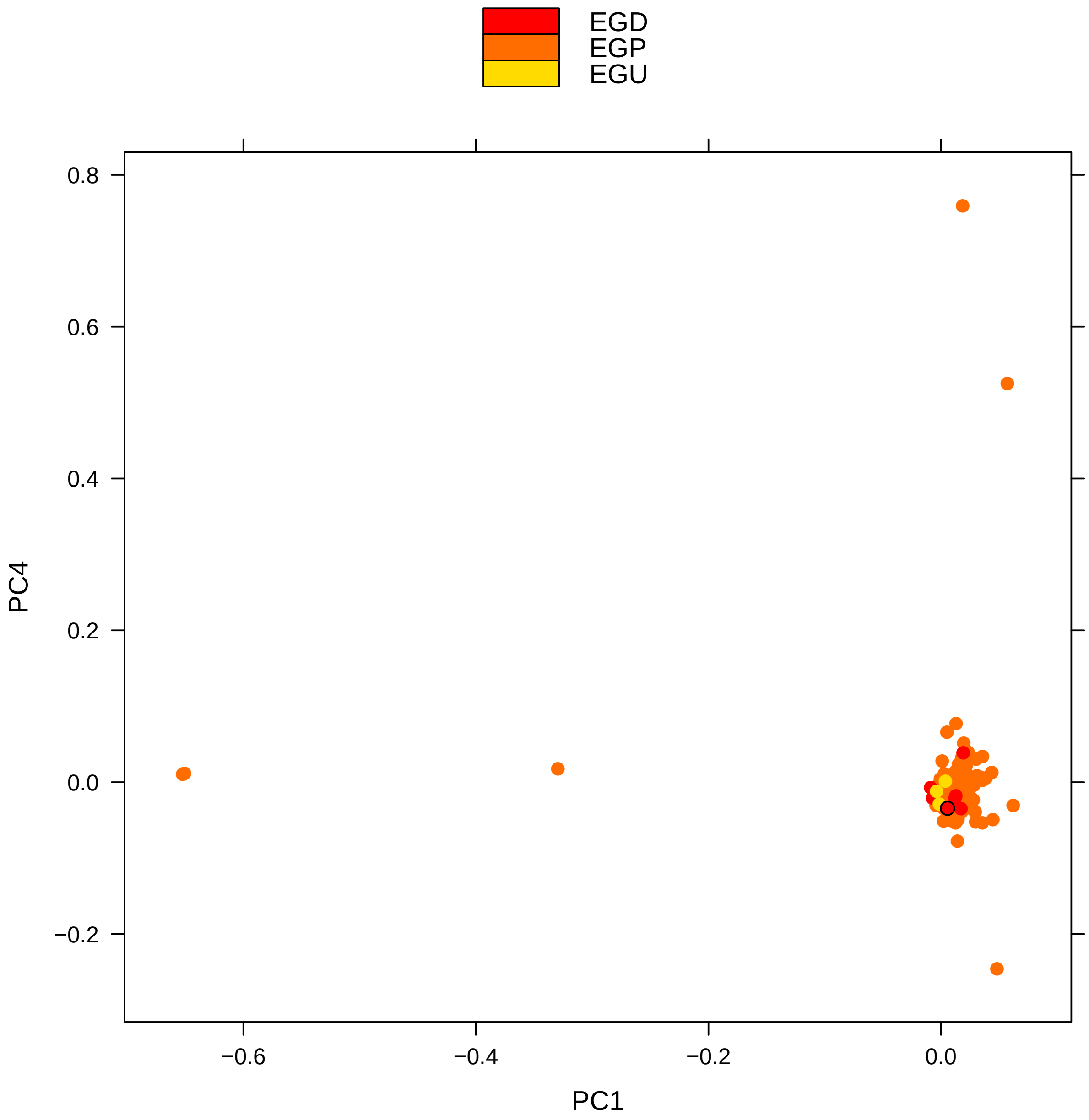

Supplementary Figure 34: Genotype principal components 1 versus 4 from Egyptian-only PCA. For label description see Suppl. Fig. 32

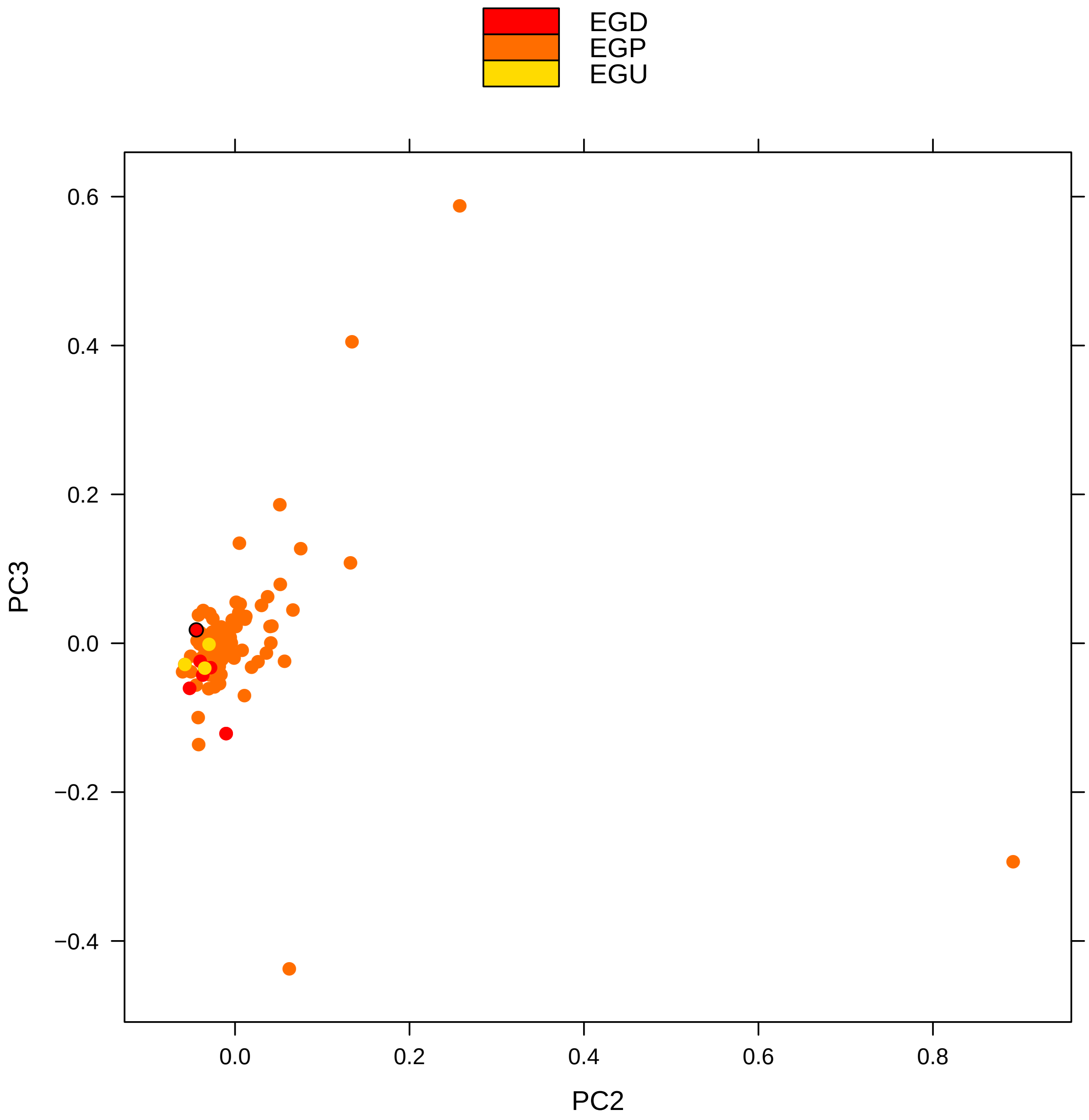

Supplementary Figure 35: Genotype principal components 2 versus 3 from Egyptian-only PCA. For label description see Suppl. Fig. 32

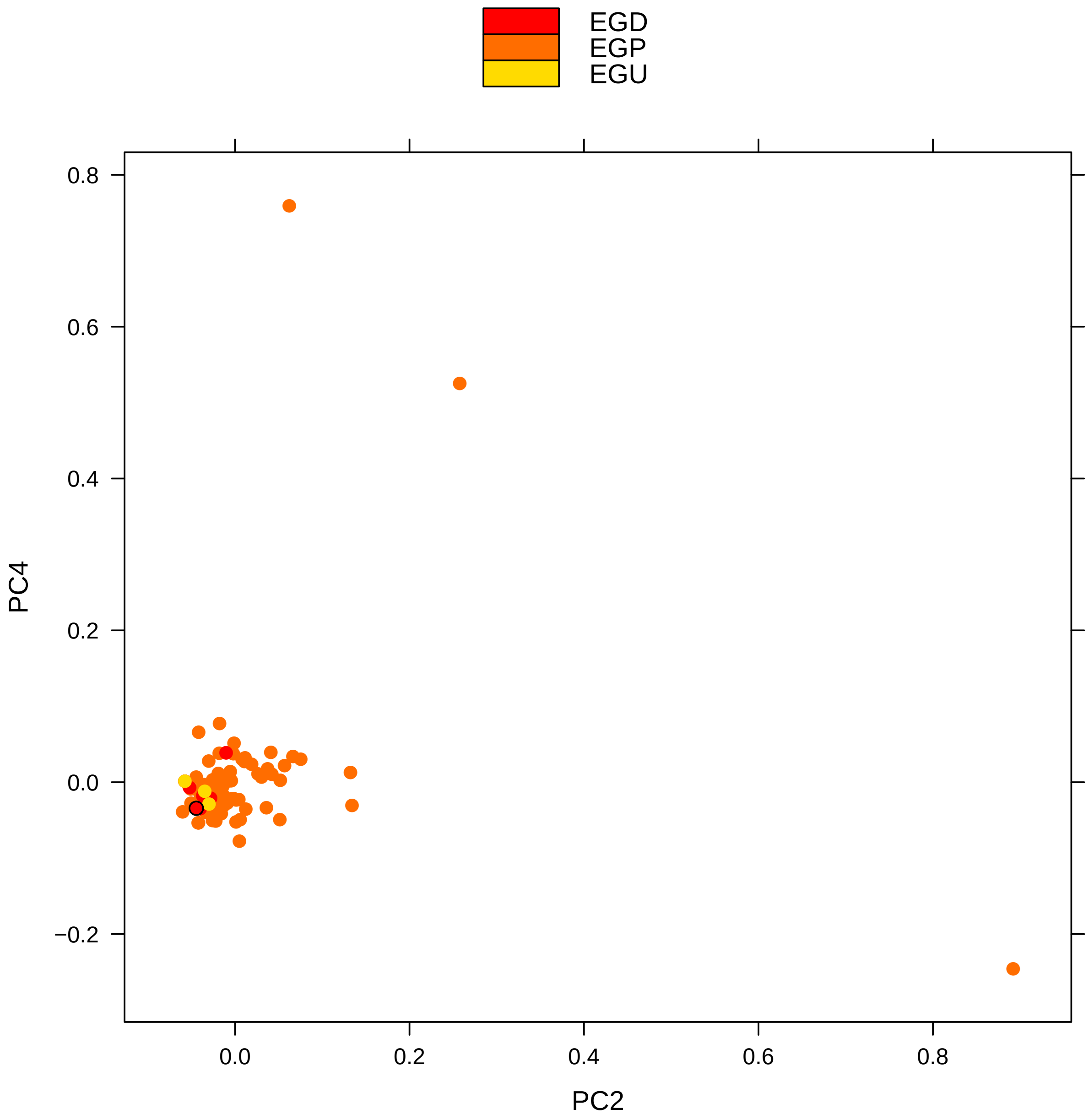

Supplementary Figure 36: Genotype principal components 2 versus 4 from Egyptian-only PCA. For label description see Suppl. Fig. 32

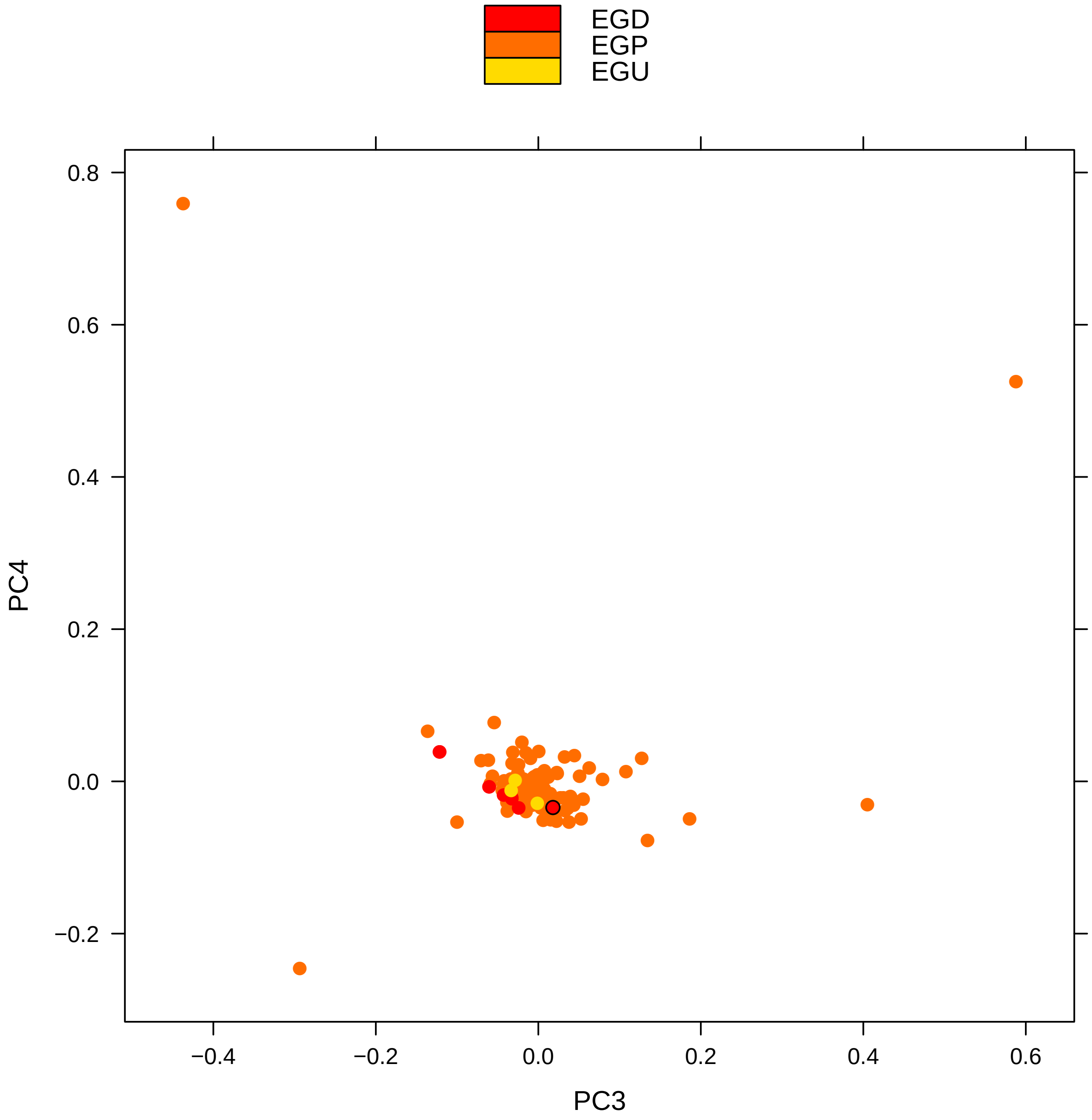

Supplementary Figure 37: Genotype principal components 3 versus 4 from Egyptian-only PCA. For label description see Suppl. Fig. 32

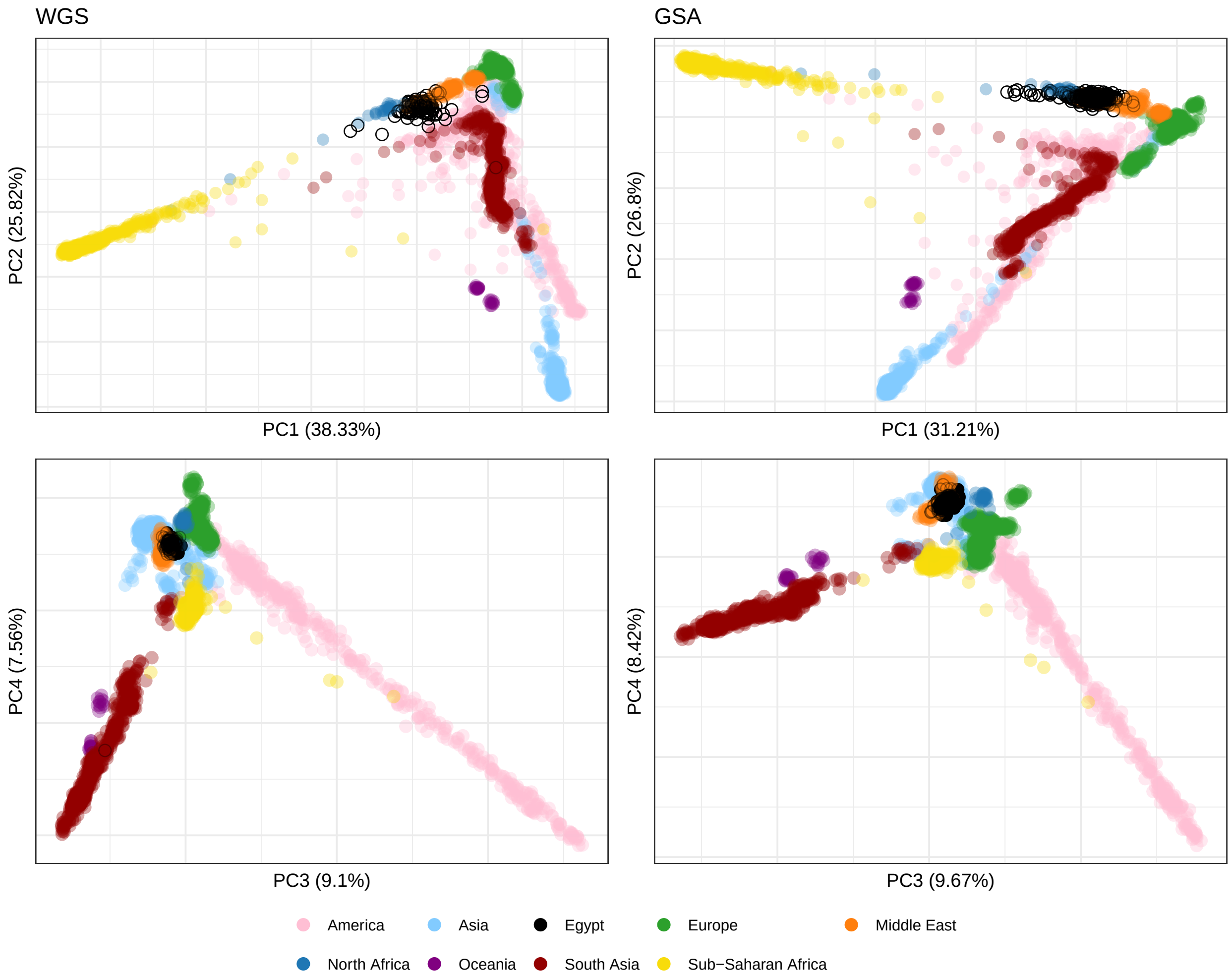

Supplementary Figure 38: Comparison of admixture analyis with WGS-based 1000G and Bergström *et al.* data for 110 WGS Egyptians (left) and 398 SNP array Egyptians (right). Egyptians are circled black. Top row: PC 1 versus PC 2. Bottom row: PC 3 versus PC 4.

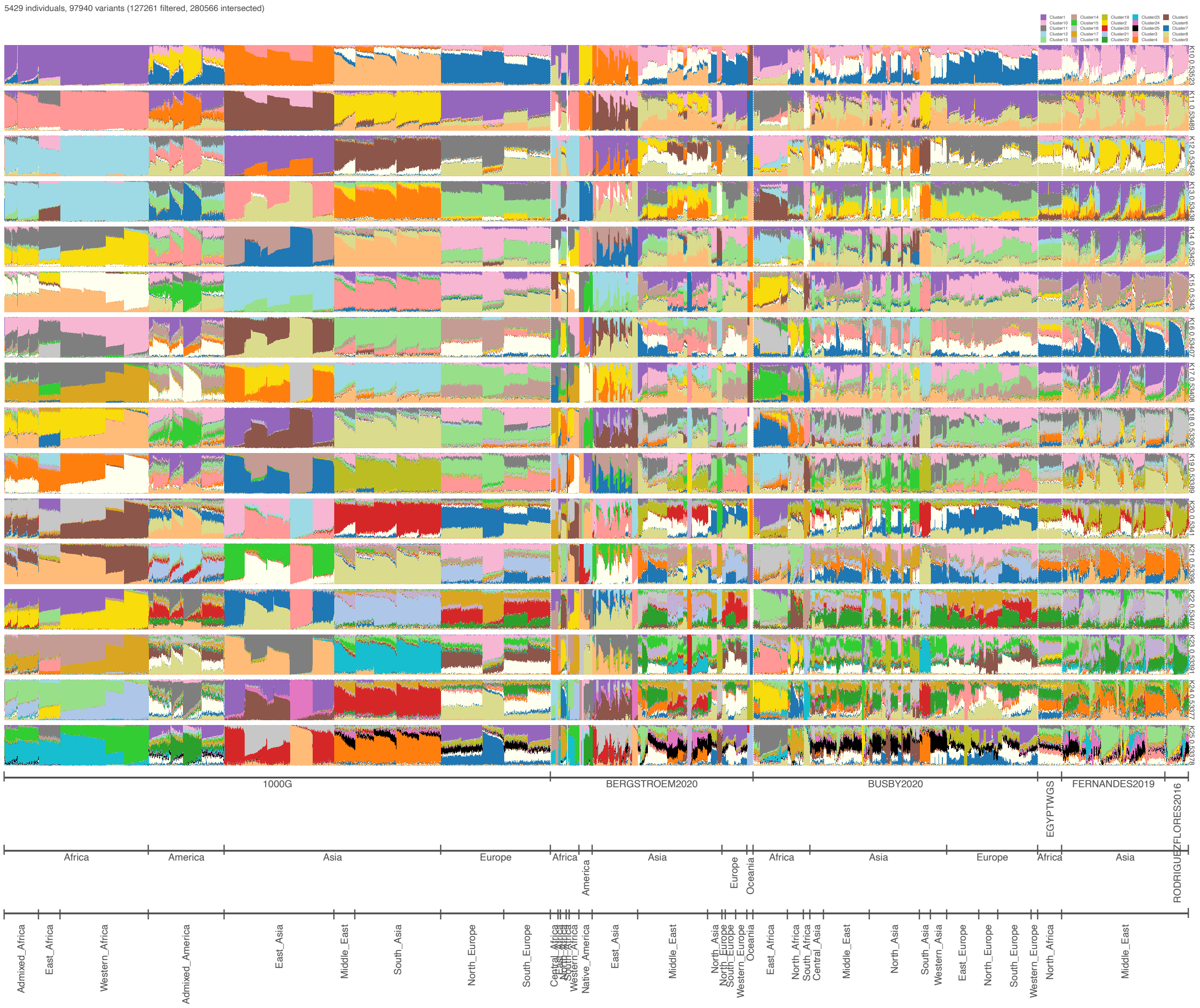

Supplementary Figure 39: Results of admixture analysis for different number of components $k$.

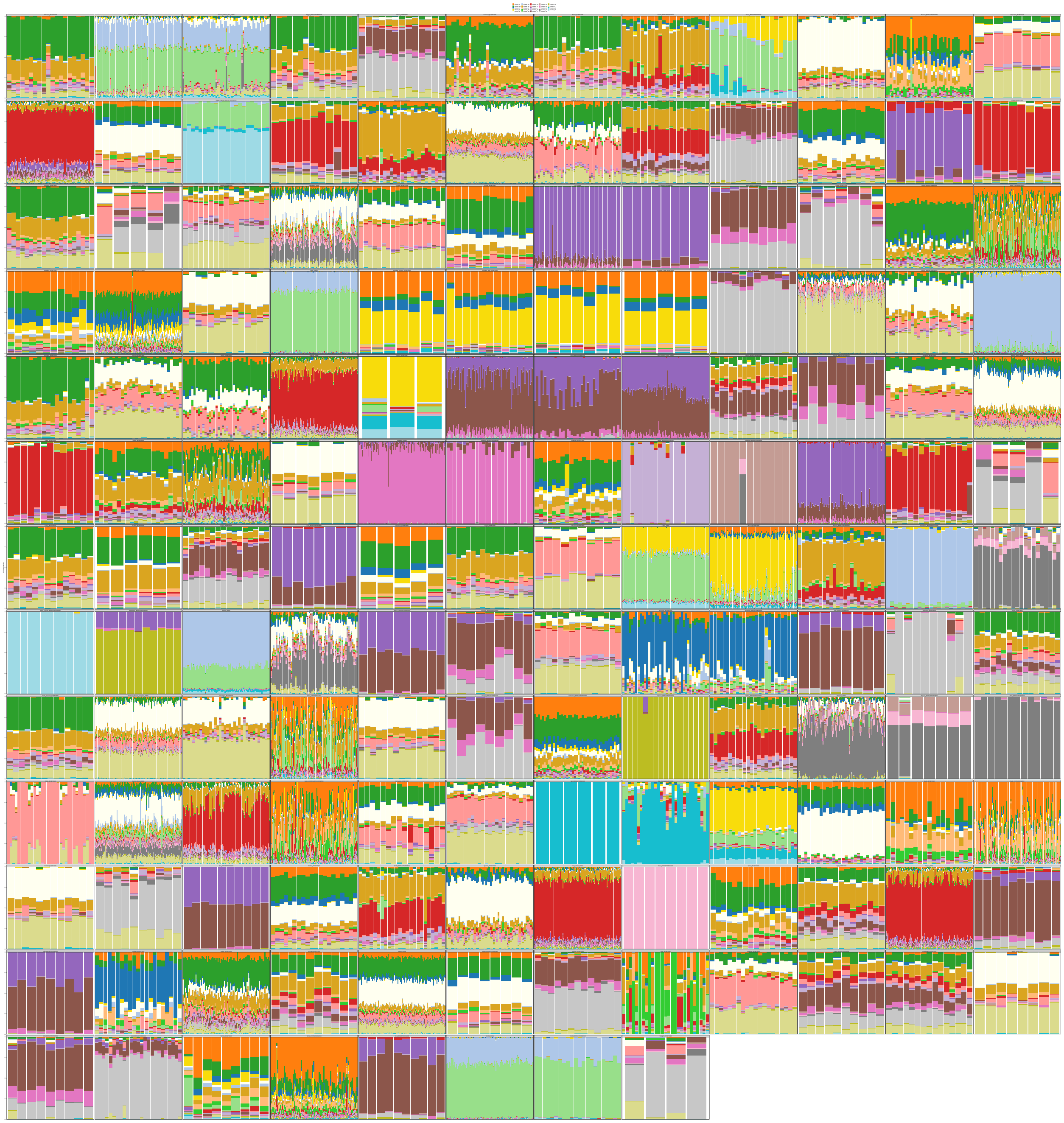

Supplementary Figure 40: Individual-level admixture results for all analyzed populations. Eight populations are part of two data sets. Components have been ordered from top to bottom in the order of their contribution in the Egyptian WGS data set. This Suppl. Fig. refers to admixture analysis for $k = 24$ visualized in Suppl. Fig. 39 (with same color-code, but here components re-ordered by Egyptian contribution)

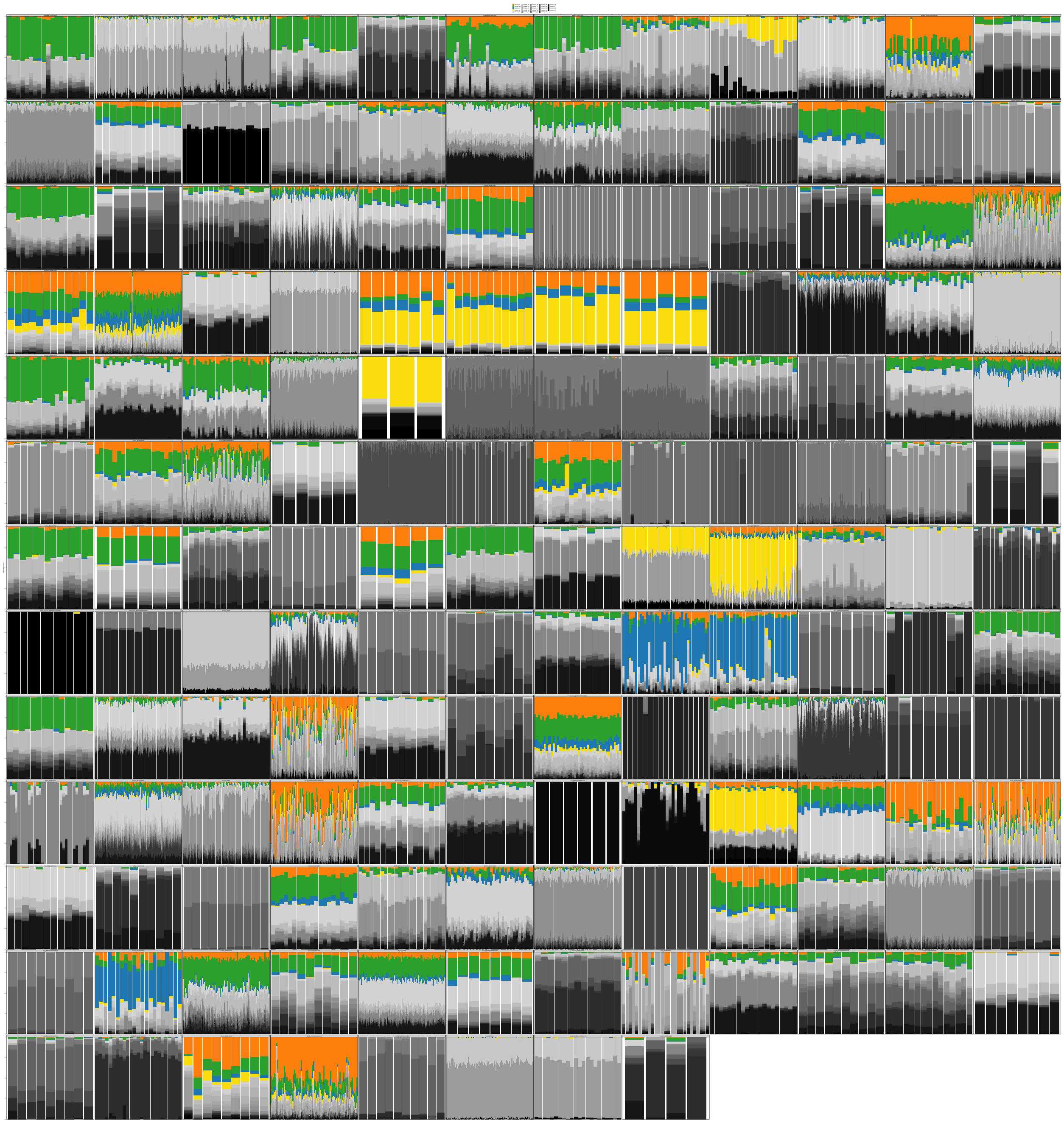

Supplementary Figure 41: Individual-level admixture results for all analyzed populations. Eight populations are part of two data sets. Components have been ordered from top to bottom in the order of their contribution in the Egyptian WGS data set. Only the four largest Egyptian components are shown in color and the remaining components in grayscale. See Suppl. Fig. 40 for distinguishing all components.

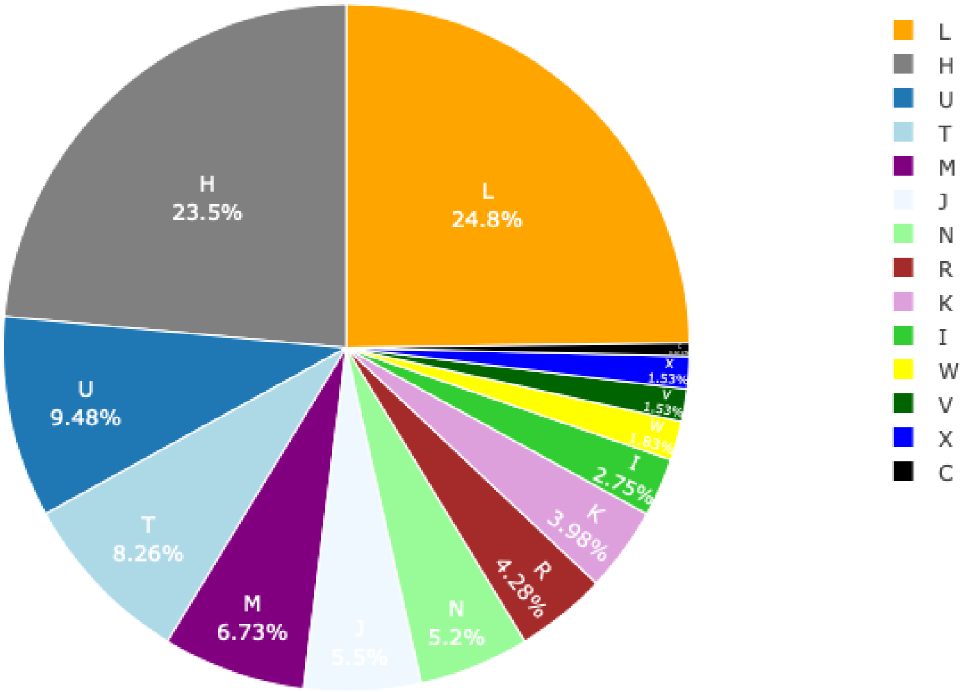

Supplementary Figure 42: Pie chart of mitochondrial haplogroups of 327 Egyptian individuals.

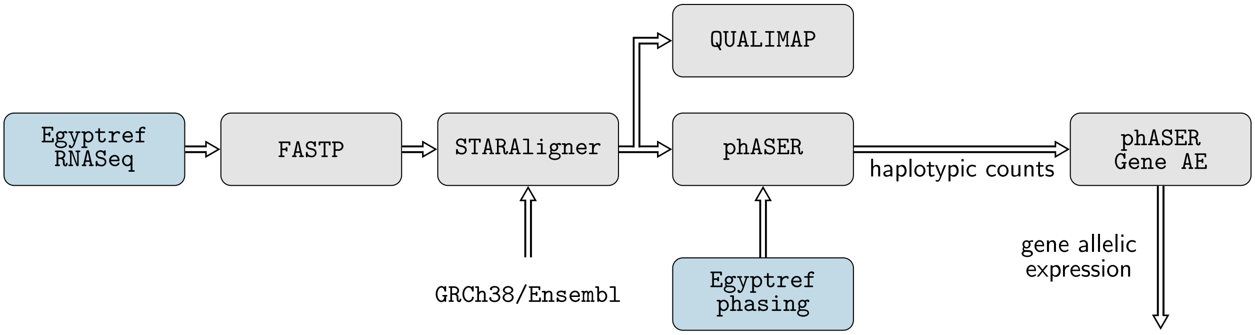

Supplementary Figure 43: Overview of haplotypic expression analysis using PHASER.

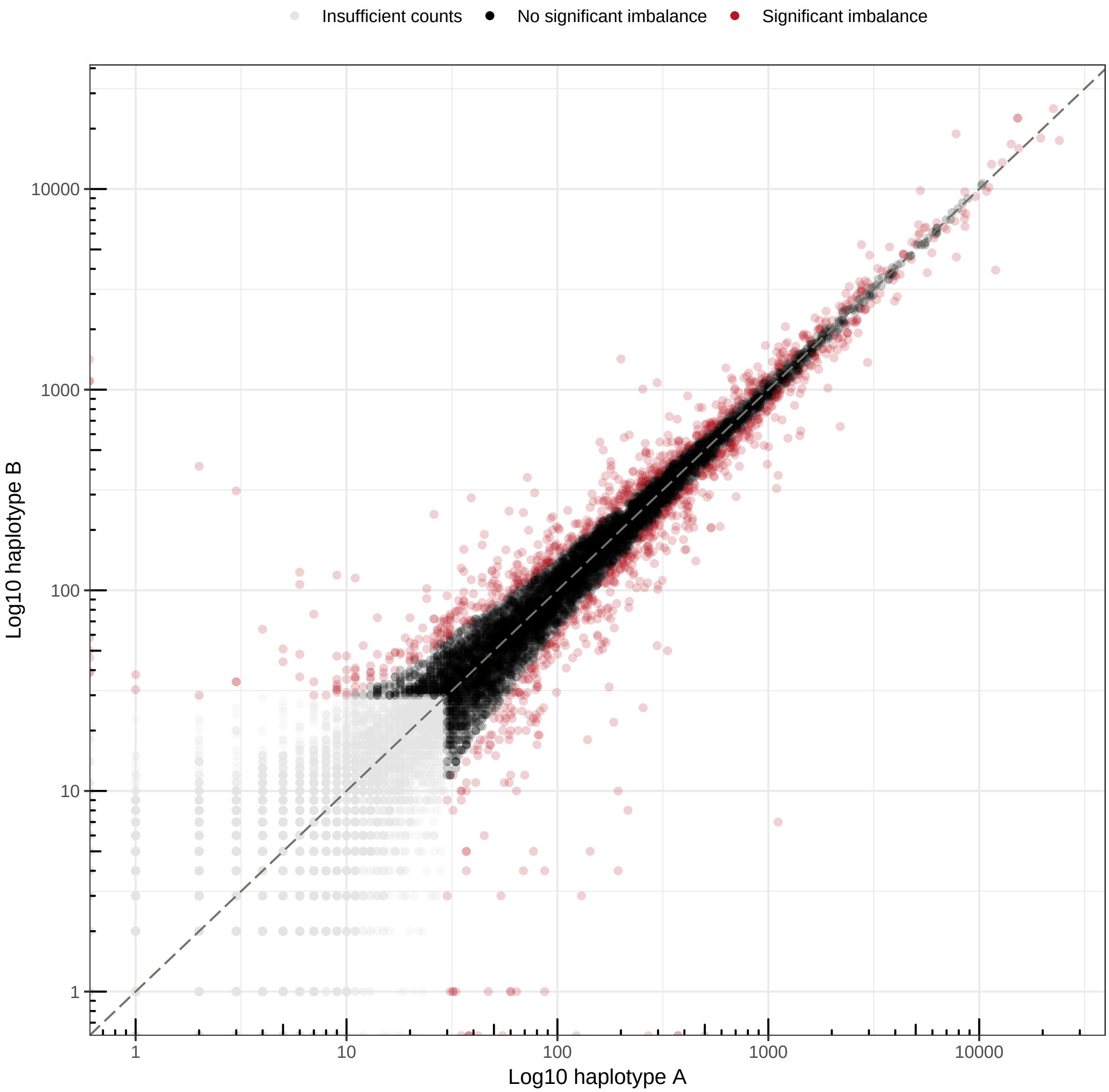

Supplementary Figure 44: Haplotypic counts for 16,566 genes. Genes with insufficient number of reads are displayed light gray and have not been used in allelic expression analysis. Significant genes are displayed red.

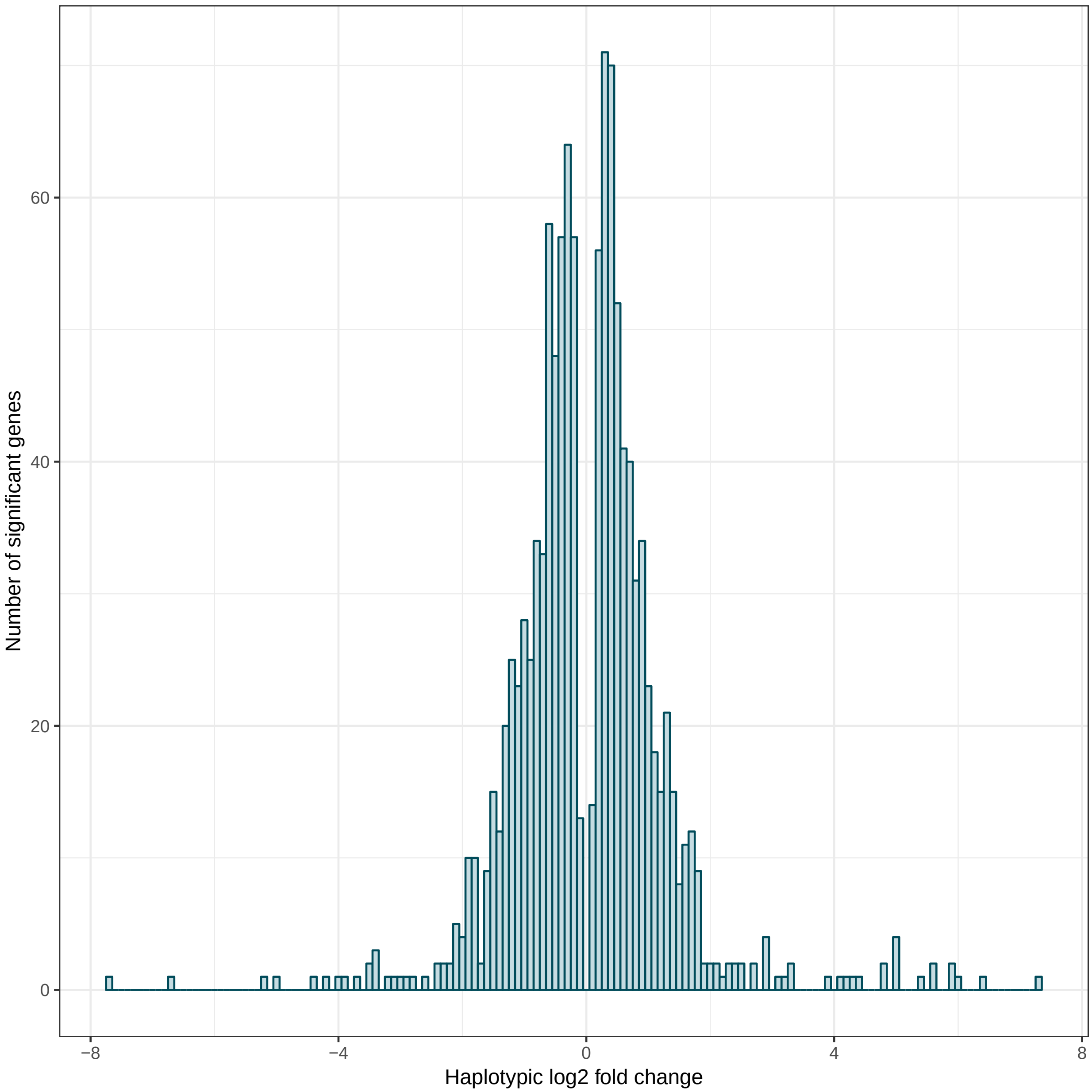

Supplementary Figure 45: Histogram of haplotypic fold changes for 1,180 genes with significant haplotypic expression.

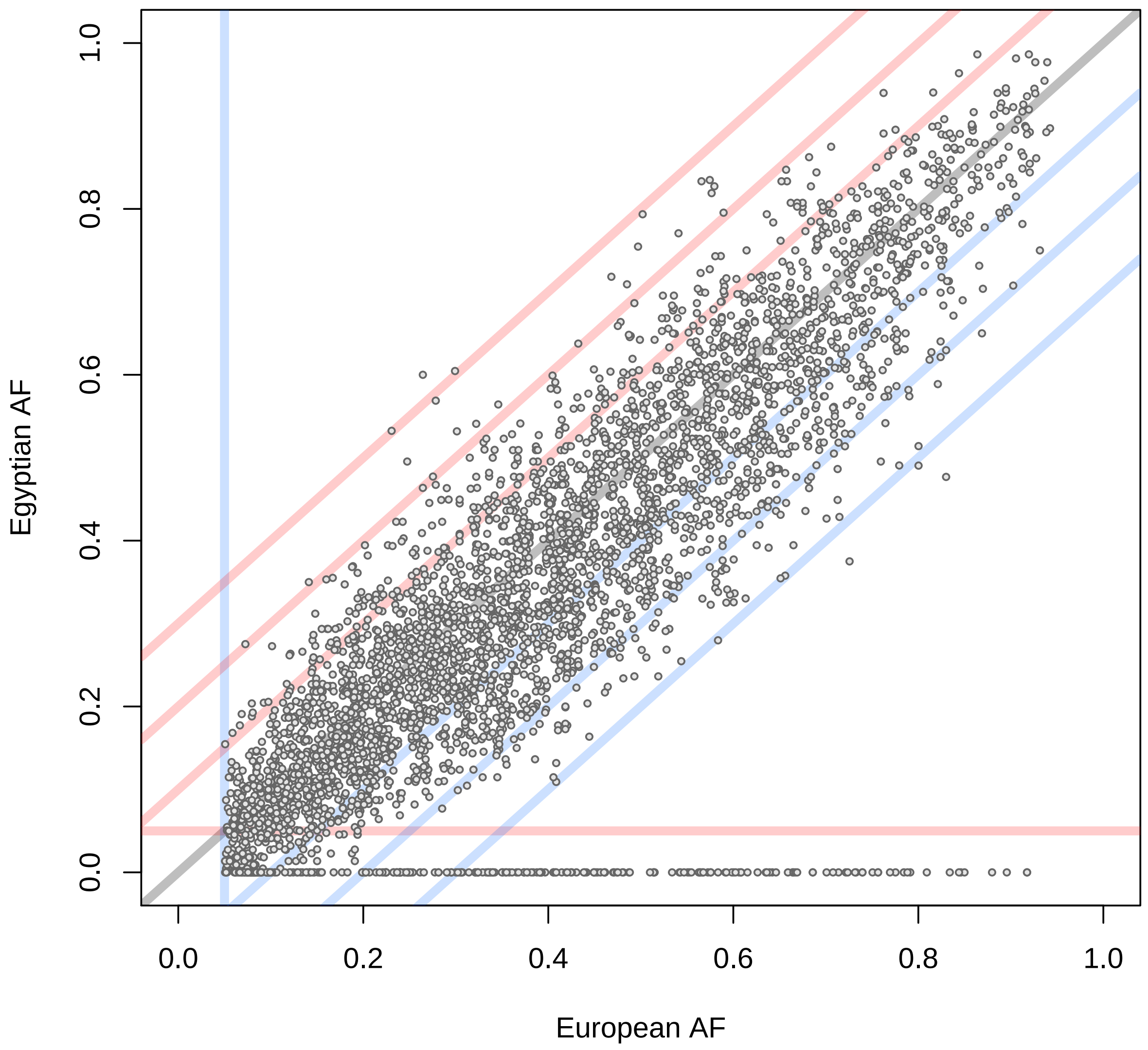

Supplementary Figure 46: Scatterplot of Egyptian AF versus European AF for GWAS tag SNPs which have genotypes for more than 100 Egyptian individuals. The blue vertical line denotes 5% European MAF. The red horizontal line denotes 5% Egyptian MAF. Diagonal lines denote 10, 20 and 30% AF difference. Note that there is a number of variants that are not present in the Egyptian data, i.e. have AF of 0%.

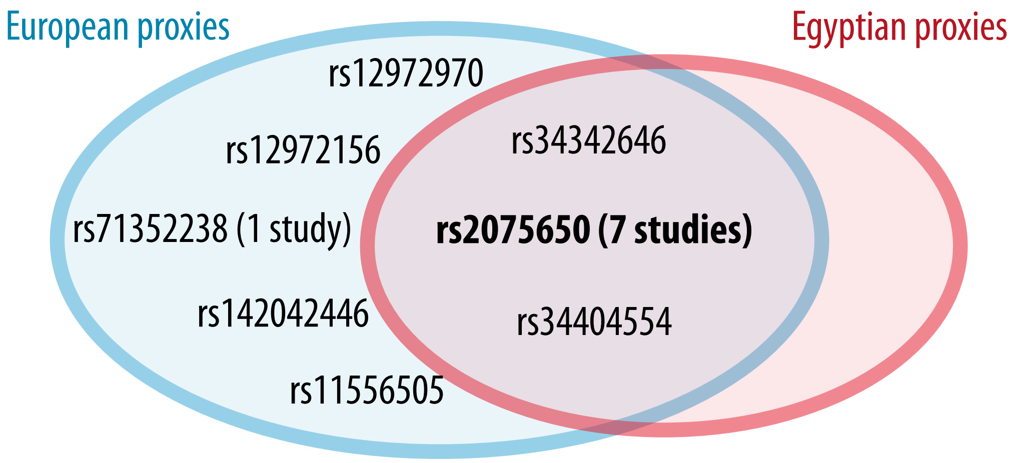

Supplementary Figure 47: Proxy SNP comparison for an Alzheimer's disease locus sometimes attributed to gene TOMM40. In seven Alzheimer's diesease GWAS, SNP rs207650 has been reported as tag SNP according to GWAS catalog. For this SNP, there are two shared proxy variants, which are in LD with rs207650 in Europeans as well as Egyptians ($R^2 \geq 0.8$). Further, there are five European-only proxies with Alzheimer's disease tag SNP rs207650. One variant, rs71352238, has also been reported in a study, illustrating that results of GWAS performed with European individuals may not be transferred to the Egyptian population because of LD differences.

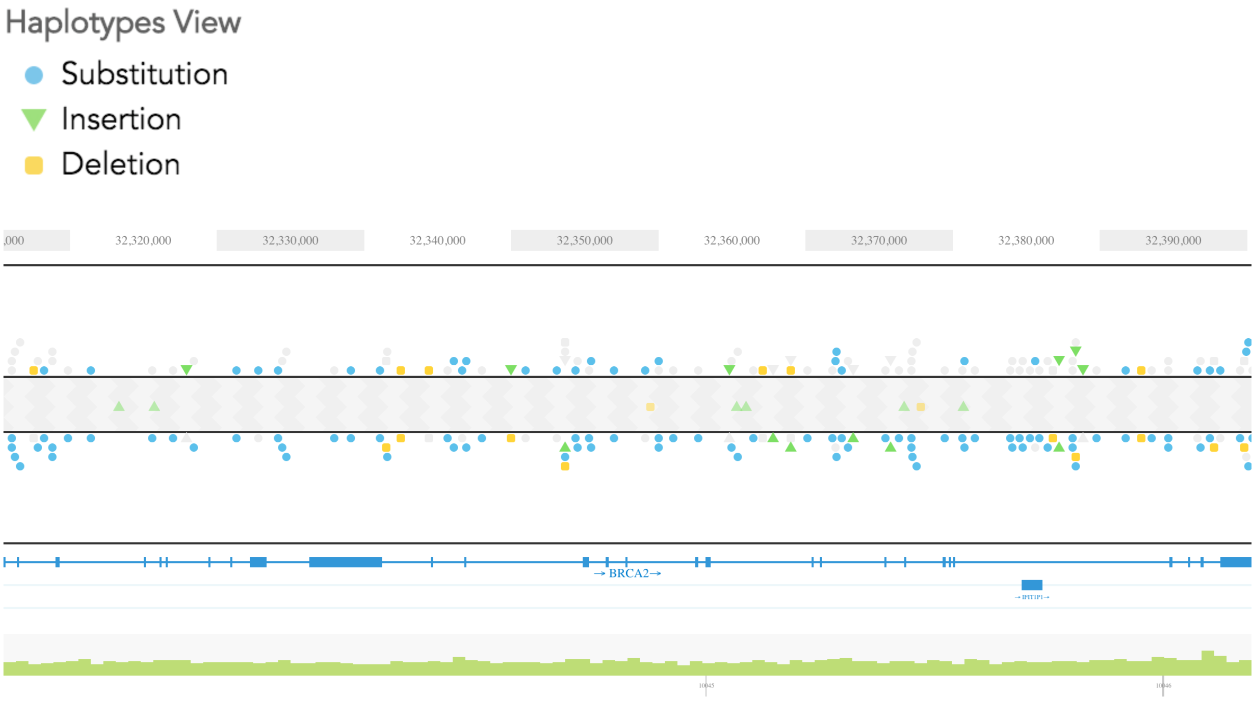

Supplementary Figure 48: View of variant phasing for the BRCA2 gene in the 10x Genomics LOUPE browser (version 2.1.2). The BRCA2 gene lies within a phasing block from chr13:25,831,216-33,523,430 (black box extending beyound the view). Each panel shows the variant allele on one chromosome, gray color within these two panels denotes a reference allele. The gray panel in the center contains unphased variants.

### Supplementary References

[1] Jue Ruan and Heng Li. Fast and accurate long-read assembly with wtdbg2. *bioRxiv*, page 530972, January 2019.

[8] Kishwar Shafin, Trevor Pesout, Ryan Lorig-Roach, Marina Haukness, Hugh E. Olsen, Colleen Bosworth, Joel Armstrong, Kristof Tigyi, Nicholas Maurer, Sergey Koren, Fritz J. Sedlazeck, Tobias Marschall, Simon Mayes, Vania Costa, Justin M. Zook, Kelvin J. Liu, Duncan Kilburn, Melanie Sorensen, Katy M. Munson, Mitchell R. Vollger, Evan E. Eichler, Sofie Salama, David Haussler, Richard E. Green, Mark Akeson, Adam Phillippy, Karen H. Miga, Paolo Carnevali, Miten Jain, and Benedict Paten. Efficient <em>de novo</em> assembly of eleven human genomes using PromethION sequencing and a novel nanopore toolkit. *bioRxiv*, page 715722, January 2019.

[9] Sergey Nurk, Brian P. Walenz, Arang Rhie, Mitchell R. Vollger, Glennis A. Logsdon, Robert Grothe, Karen H. Miga, Evan E. Eichler, Adam M. Phillippy, and Sergey Koren. HiCanu: accurate assembly of segmental duplications, satellites, and allelic variants from high-fidelity long reads. *bioRxiv*, page 2020.03.14.992248, January 2020.

[10] Kai Wang, Mingyao Li, and Hakon Hakonarson. ANNOVAR: functional annotation of genetic variants from high-throughput sequencing data. *Nucleic Acids Research*, 38(16):e164, September 2010.

[11] William McLaren, Laurent Gil, Sarah E. Hunt, Harpreet Singh Riat, Graham R. S. Ritchie, Anja Thormann, Paul Flicek, and Fiona Cunningham. The Ensembl Variant Effect Predictor. *Genome Biology*, 17(1):122, 2016.

[12] Philipp Rentzsch, Daniela Witten, Gregory M. Cooper, Jay Shendure, and Martin Kircher. CADD: predicting the deleteriousness of variants throughout the human genome. *Nucleic Acids Research*, 47(D1):D886–D894, January 2019.

[13] Ivan Adzhubei, Daniel M. Jordan, and Shamil R. Sunyaev. Predicting functional effect of human missense mutations using PolyPhen-2. *Current Protocols in Human Genetics*, Chapter 7:Unit7.20, January 2013.
